# Efficient assembly of plant genomes: A case study with evolutionary implications in *Ranunculus* (Ranunculaceae)

**DOI:** 10.1101/2023.08.08.552429

**Authors:** Kevin Karbstein, Nancy Choudhary, Ting Xie, Salvatore Tomasello, Natascha D. Wagner, Birthe H. Barke, Claudia Paetzold, John P. Bradican, Michaela Preick, Axel Himmelbach, Nils Stein, Argyris Papantonis, Iker Irisarri, Jan de Vries, Boas Pucker, Elvira Hörandl

**Affiliations:** University of Göttingen, Albrecht-von-Haller Institute for Plant Sciences, Department of Systematics, Biodiversity and Evolution of Plants (with Herbarium), Göttingen, Germany; Max Planck Institute for Biogeochemistry, Department of Biogeochemical Integration, Jena, Germany; TU Braunschweig, Institute of Plant Biology, Braunschweig, Germany; University Medical Center Göttingen, Institute of Pathology, Göttingen, Germany; Senckenberg Naturhistorische Sammlungen, Dresden, Germany; University of Potsdam, Institute for Biochemistry and Biology, Potsdam, Germany; Leibniz Institute of Plant Genetics and Crop Plant Research (IPK), Seeland, Germany; Center of integrated Breeding Research (CiBreed), Department of Crop Sciences, Göttingen, Germany; Museo Nacional de Ciencias Naturales (MNCN-CSIC), Department of Biodiversity and Evolutionary Biology, Madrid, Spain; University of Göttingen, Institute for Microbiology and Genetics, Department of Applied Bioinformatics, Göttingen, Germany; University of Göttingen, Campus Institute Data Science (CIDAS), Göttingen, Germany; University of Göttingen, Göttingen Center for Molecular Biosciences (GZMB), Department of Applied Bioinformatics, Göttingen, Germany

**Keywords:** gene evolution, large plant genomes, plastome, mitogenome, nuclear genome, Illumina vs. Nanopore vs. PacBio sequencing, *de novo* assembly strategies, *Ranunculus auricomus*, Ranunculaceae

## Abstract

Currently, it is still a challenge - in terms of laboratory effort and cost, as well as assembly quality - to unravel the sequence of large and complex genomes from non-model/crop plants. This often hampers the study of evolutionarily intricate species groups. The species-rich genus *Ranunculus* (Ranunculaceae) is an important angiosperm group for the study of polyploidy, apomixis, reticulate evolution, and biogeography. However, neither mitochondrial, nor high-quality nuclear genome sequences are available. This limits phylogenomic, functional, and taxonomic analyses thus far. Here, we tested Illumina short- read, Oxford Nanopore Technology (ONT) or PacBio/HiFi long-read, and hybrid-read assembly strategies. We used the diploid progenitor species *R. cassubicifolius* (*R. auricomus* complex), and selected the best assemblies in terms of completeness, contiguity, and quality scores. We first assembled the plastome (156 kbp, 85 genes) and mitogenome (1.18 Mbp, 40 genes) sequences using Illumina and Illumina-PacBio-hybrid strategies, respectively. We also present an updated plastome and the first mitogenome phylogeny of Ranunculaceae, including studies of gene loss (e.g., *infA*, *ycf15*, or *rps*) with evolutionary implications. For the nuclear genome, we favored a PacBio-based assembly three-times polished with filtered reads and subsequently scaffolded into 8 pseudochromosomes by chromatin conformation data (Hi-C) as the representative sequence. We obtained a haploid genome sequence with 2.69 Gbp, 94.1% complete BUSCO ‘embryophyta_odb10’ genes found, and 31,322 annotated genes. The genomic information presented here will improve phylogenomic analyses in this species complex, and will enable advanced functional, evolutionary, and biogeographic analyses for the genus and beyond Ranunculaceae in the future.

**Significance Statement:** The genus *Ranunculus* is a model system in flowering plants for polyploidy, apomixis, evolution, and biogeography research. We present the first nuclear and mitochondrial genome sequence, and the plastid evolution of Ranunculaceae. Using Illumina, ONT, and PacBio data, we developed an efficient assembly strategy that can be applied to other non-model plants. Results presented here are useful for improving population genomic and phylogenomic analyses, and enable better functional analyses at species, genus and family level.

## Introduction

Over the past decade, the scientific community has experienced a massive expansion in accessible sequencing technologies. For constructing genome assemblies, short-reads (e.g., Weisenfeld et al., 2018; Prjibelski et al., 2020), long-reads (e.g., Koren et al., 2017; Cheng et al., 2021), or a combination of both (hybrid-read; e.g., Di Genova et al., 2021; Gatter et al., 2021; Zimin and Salzberg, 2022) have frequently been used. Nevertheless, biologists have moved beyond short-read (100-250 bp) Illumina sequence data to perform genome assemblies. Short reads often produce incomplete assemblies, and lack the contiguity required to resolve repetitive regions that plague large genomes, leading to highly fragmented and incomplete *de novo* genome assemblies (Gatter et al., 2021; Rhie et al., 2021; Zimin and Salzberg, 2022). Today, long-read technologies, such as nanopore sequencing (Oxford Nanopore Technologies, ONT) or single molecule real time sequencing (SMRT; Pacific Biosciences), can bridge such gaps and improve the quality of genome assemblies. High-fidelity (HiFi) reads derived from SMRT sequencing are typically of high per-base quality (>99.5%) and up to 25 kbp in length. In contrast, ONT sequencing just crossed the 99% per-base accuracy threshold, read lengths are limited only by the input DNA fragment size thus reaching >1 Mbp, and can be easily performed in any laboratory (reviewed in Pucker et al., 2022).

HiFi or ONT sequencing, combined with proximity information obtained by chromosome conformation capture sequencing (Hi-C), has started to become prevalent for chromosome-scale, high-quality model but also non-model plant genome sequences (e.g., Mascher et al., 2017; Xie et al., 2020; Hoencamp et al., 2021; Qu et al., 2023). The quality and completeness of the assembly can be improved by polishing strategies using available, quality-filtered Illumina, ONT, or PacBio reads (e.g., Zimin and Salzberg, 2020; Dmitriev et al., 2021; Chang et al., 2023). The combination of low- to medium-sequencing coverage (that is, the proportion of the total assembly that is supported by the raw reads) long-read data with highly accurate, often already available Illumina sequences in hybrid assemblies is particularly attractive for improving genome assemblies when resources are limited. This is usually the case when working with non-model plants.

Producing “high-quality” genome sequences, that is genome assemblies characterized by > 90% completeness with respect to expected genome size, > 1-10 Mbp contiguity (N_50_), and > 90% recovery rate of conserved protein-coding genes (BUSCO; see assembly standards of the Earth Biogenome Project; Table S1 in Lawniczak et al., 2022), represent an important target for research groups. However, some metrics are rather adapted to animals. For example, flowering plants have undergone many ancient or recent polyploidization events (Van De Peer et al., 2017; Leebens-Mack et al., 2019; Hörandl, 2022) and are characterized by quite large, repetitive genomes with duplicated sequence content. Consequently, a BUSCO ‘embryophyta_odb10’ score of > 90% single-copy genes can often not be met because genes are often duplicated in the genome (Dmitriev et al., 2021) or is not informative for quality assessment in non-coding, repetitive DNA regions (e.g., LTR assembly index; Mokhtar et al., 2023). Irrespective of chosen metrics, efficient strategies in terms of laboratory effort, cost, and assembly quality are needed to reliably reconstruct large and complex high-quality genome sequences for non-model organisms. This often hampers the study of evolutionary, biogeographic, and ecological relationships within groups characterized by intricate evolutionary processes such as hybridization, polyploidy, and/or apomixis (Ennos et al., 2005; Karbstein et al., 2024).

*Ranunculus* represents a cosmopolitan genus with approximately 600 species, of which about 40% are polyploid (Hörandl et al., 2005) and have rather large genomes (ranging from 1.8 to 24.6 Gbp; Leitch et al., 2019). The *Ranunculus auricomus* complex is one of the largest polyploid apomictic plant species groups in Eurasia, with more than 840 described apomictic taxa (Hörandl and Raab-Straube, 2015). The complex is a model system for understanding the evolution and biogeography of young flowering plant groups formed by reticulate evolution and allopolyploidy (Paun et al., 2006; Hörandl et al., 2009; Pellino et al., 2013; Hodač et al., 2018; Karbstein et al., 2020b, 2020a, 2021, 2022; Tomasello et al., 2020; Karbstein, 2021; Bradican et al., 2023, 2024; Hodač & Karbstein et al., 2023). It has also been frequently studied for the molecular, cytogenetic, environmental, and evolutionary mechanisms of apomixis (i.e., the asexual reproduction via seeds; Hörandl et al., 2024).

Apomixis is relevant for agricultural applications (e.g., Schmidt, 2020), and it is often associated with hybridization and/or polyploidy, which are considered major evolutionary forces in plant evolution. At least 30% of flowering plant species are of recent polyploid origin, while > 25% of extant plant species are known to be involved in interspecific hybridization or introgression events (Mallet, 2005; Soltis et al., 2015; Van De Peer et al., 2017; Leebens-Mack et al., 2019; Hörandl, 2022). Consequently, improving (phylo)genomic analyses for di- and polyploids, and observing regulatory mechanisms of apomixis (e.g., Brukhin et al., 2019) with *R. auricomus* reference genome sequences will help develop a better understanding of plant evolution and biodiversity, and is therefore of general interest in biological research. Organellar genome reconstructions are also valuable targets for studying nuclear-organelle compatibility and the chimerism of cyto-nuclear complexes (Sloan, 2013; Møller et al., 2021). Little is known, for instance, about plant mitochondrial genomes, although mitochondrial-nuclear compatibility is important for speciation, evolution of crossing barriers, and emergence of cytoplasmic male sterility (Postel and Touzet, 2020; Møller et al., 2021). Elucidating cyto-nuclear complexes may be important for studying a variety of evolutionary processes. For instance, plastid information may provide additional insights into phylogenetic relationships (Tomasello et al., 2024), maternal progenitors of hybrids (Karbstein et al., 2022), or the ability of hybridization (Sobanski et al., 2019; Postel and Touzet, 2020), or mitochondrial information into plant sex evolution and determination (e.g., male sterility; (Havird et al., 2015; Park et al., 2021; Hörandl, 2024).

The *R. auricomus* complex contains five newly circumscribed, predominantly diploid sexual species (Karbstein et al., 2020b), which gave rise to the >800 described allopolyploid apomictic taxa within the last ca. 800 kyr (Tomasello et al., 2020; Karbstein et al., 2022). The diploid genome size of the *R. auricomus* complex (2n = 16) ranges between 6.12 and 6.30 pg DNA based on DNA image cytometry (Hörandl and Greilhuber, 2002), recalculated as 5.998-6.174 Gbp (Leitch et al., 2019). Therefore, *R. auricomus* genomes are large even at the diploid level. Increasing proportions of repetitive regions are responsible for genome size inflation (Bennetzen, 2005; Peška et al., 2019; Guo et al., 2021; Jayakodi et al., 2023), which could also be relevant for *R. auricomus* genomes (Hörandl et al., 2009; Hodač et al., 2018; Karbstein et al., 2022). However, so far, high-quality reference genome sequences for the species complex, but also for the entire genus *Ranunculus* are missing. In Ranunculaceae, the only other reference genome sequences are available from diploid *Aquilegia* and *Coptis* species (Xie et al., 2020; Liu et al., 2021; Mokhtar et al., 2023), but these genomes are much smaller (ca. 1 Gbp; Leitch et al., 2019), and no polyploid species complexes are known in these genera (TROPICOS database v3.4.2, http://www.tropicos.org; CCDB database v1.66, https://taux.evolseq.net/CCDB_web; Rice et al., 2015).

The evolutionary relationships among the diploid sexual progenitors of *R. auricomus* have recently been clarified (Karbstein et al., 2020b; Tomasello et al., 2020). A diploid *Ranunculus* reference genome sequence would be a major step forward in improving the reconstruction of the species by facilitating read mapping to orthologous loci, allele phasing, and subgenome assignments in hybrids; it would be the starting point for more sophisticated approaches such as reference-guided genome assemblies (e.g., Eaton and Overcast, 2020), polyploid phasing used for multilabeled species trees (e.g., Dauphin et al., 2018), or allopolyploid network reconstructions (e.g., Jones, 2017; Šlenker et al., 2021). In addition, the annotation of thousands of nuclear genes would also help enable sophisticated structural and functional analyses in the species complex. New developments allow the incorporation of external hints such as RNA-seq data or the annotation of homologous genes through knowledge transfer from other species (Pucker, 2024). Tools are also needed for automatically extracting and comparing group-specific annotations across genome sequences from databases, which are essential for evolutionary studies. In *Ranunculus*, and in Ranunculaceae, such large-scale studies are missing. Annotated high-quality genome sequences are essential for functional genomics in the context of apomixis, adaptation, and gene expression (e.g., Syngelaki et al., 2021), and for studying loci under selection and mutation-selection dynamics (e.g., Del Amparo et al., 2021) that could so far only be performed on (incomplete) transcriptomes (Pellino et al., 2013; Paetzold et al., 2022).

To generate plastid, mitochondrial, and high-quality nuclear genome assemblies of a diploid species of the *R. auricomus* complex, we applied different assembly strategies using short Illumina reads and long ONT or PacBio reads under low to medium sequencing coverage, with the best nuclear-genome assembly being scaffolded by Hi-C reads. This represents a typical scenario when working with non-model plant groups on a limited budget. We thus focused on applying different approaches based on the available sequencing data. To study organellar gene evolution in detail, we also downloaded and analyzed all available sequences from NCBI.

## Methods

### Plant Material

In the present study, analyses were performed on diploid sexual individuals (2n = 16) of *Ranunculus cassubicifolius* (*R. auricomus* complex; taxonomic treatment follows Karbstein et al., 2020b). Chromosome numbers, measurements of ploidy and mode of reproduction are published in Hörandl and Greilhuber (2002) under ‘*R. carpaticola*’, and in Karbstein et al. (2020b, 2021). Fresh leaf material was harvested in 2019 from individual EH8483/10 (NCBI BioSample SAMN27753242), sampled in Revúca, Slovakia (48°41’39"N, 20°07’42"E; Hörandl and Greilhuber, 2002; under ‘*R. carpaticola*’). High-molecular-weight DNA was extracted immediately after harvest and used for Illumina DNA sequencing. The same population was already used for transcriptome (Pellino et al., 2013) and population genetic analyses (Paun et al., 2006; Hodač et al., 2018). In autumn 2021, we harvested fresh, young leaf material for ONT sequencing, after 12-hour dark-adaptation at room temperature in the early morning from another conspecific individual collected nearby (LH040/02, SAMN27753242; 48°40’38"N, 20°06’24"E; Figure 1; Karbstein et al., 2020b) because the previously used individual died. In 2023, we used individuals from the same population for PacBio and Hi-C sequencing, LH040/06 and LH040/05+06 (SAMN44350262), respectively, because the previously harvested individual did not sprout again. All plants were cultivated at the Old Botanical Garden of the University of Göttingen.

**Figure 1.**
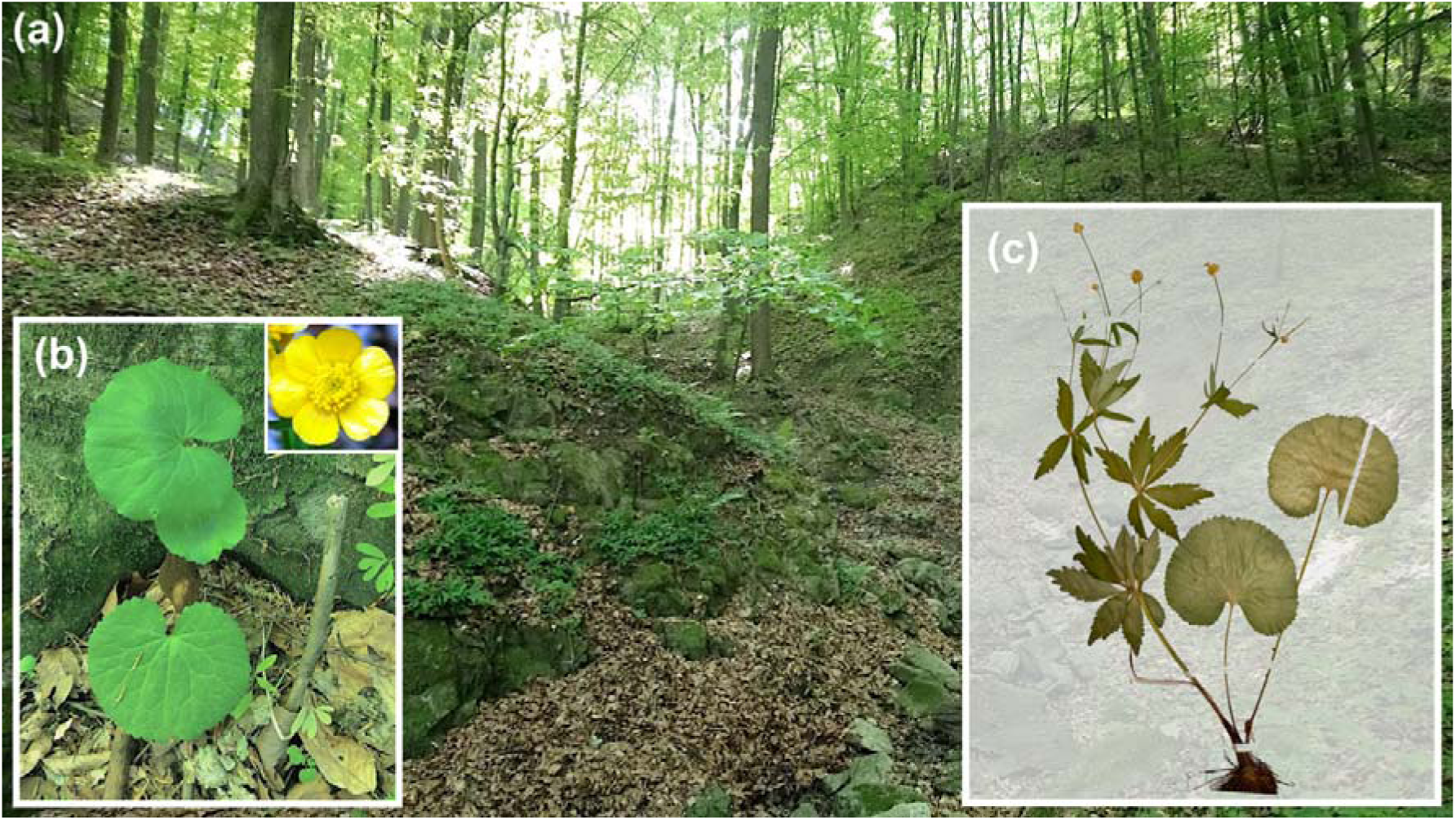
(a) Habitat of *Ranunculus cassubicifolius* s.l. (previously listed as ‘*R. carpaticola*’) population LH040 in a forest near Liptov, Slovakia (4^th^ May 2018, L. Hodač & K. Spitzer; details in Karbstein et al., 2020b). (b) Shown are the typical undissected basal leaves *in situ* and a typical *R. cassubicifolius* flower from a different population (LH009) and (c) the scan of a representative herbarium specimen from LH040. Field images: L. Hodač.

### Illumina DNA extraction and sequencing

For Illumina short-read sequencing, genomic DNA (gDNA) was directly extracted from ca. 200 mg of fresh leaf material from a single *R. cassubicifolius* s.l. individual (EH8483/10) using liquid nitrogen and the DNeasy PowerPlant Pro Kit (Qiagen, Hilden, Germany) according to the manufacturer’s instructions. DNA size and concentration were measured with the Qubit Fluorometer 2.0 and the Qubit dsDNA HS Assay Kit (ThermoFisher Scientific, Waltham, USA), and the Genomic DNA ScreenTape device (Agilent Technologies, Santa Clara, USA). The extraction yielded 16 ng/µl of DNA with a median fragment length of 19 kb.

Library preparation followed the 10x Chromium Controller Genomics Kit (10x Genomics, Pleasanton, USA) according to the manufacturer’s instructions. Library quality was assessed by a qPCR based on the NEBNext Library Quant Kit (New England Biolabs Inc., Ipswich, USA). Sequencing with standard Illumina primers was performed on a NextSeq 500/550 instrument (Illumina, San Diego, USA) with 2 x 150 bp reads.

We converted the raw output into FASTQ files using SUPERNOVA v2.1.1 (10x Genomics, San Francisco, USA; Marks et al., 2017; Weisenfeld et al., 2018) and BLC2FASTQ v2.20 (Illumina, San Diego, USA), resulting in 51 Gb raw FASTQ files. We conducted polyX-tail and adapter trimming, and quality filtering of Illumina raw reads using ATRIA v3.1.2 (Chuan et al., 2021) with default settings, and BBTools/BBMap v39.01 (‘repair.sh’; (Bushnell, 2022) to reorder filtered paired-end reads.

### Nanopore DNA extraction and sequencing

For ONT, gDNA extraction was mainly performed according to the Qiagen protocol for the isolation of gDNA from plants and fungi, and the manufacturer’s instructions of the Qiagen Genomic DNA Buffer Set using Genomic Tip 20/G (see details in Text S1). Several protocol modifications were applied to increase DNA yield and to optimize DNA quality (see also Vaillancourt and Buell, 2019; Li et al., 2020). Three gDNA extractions were performed in parallel from a single *R. cassubicifolius* s.l. individual (LH040). Fresh leaf material (ca. 75 mg) was frozen in liquid nitrogen and ground with a mortar and pestle, and transferred into a 2.0 ml Eppendorf tube containing 400 µl API Qiagen extraction buffer (Qiagen DNeasy Plant Mini Kit).

The mixture was incubated at 40°C for 1.5 hours with gentle agitation. Then, 4 µl Qiagen RNase A (100 mg/ml) was added, and the mixture was incubated again at 37°C for 30 minutes, followed by the addition of 18 µl Qiagen Proteinase K (20 mg/ml), and incubation at 50°C for 2 h with gentle agitation. The mixture was centrifuged at 12,000 x g for 20 min, and ca. 400 µl of the lysate was transferred with wide-bore tips to a buffer QBT-equilibrated Genomic Tip. DNA was washed with 5 x 1 ml QC Buffer, and eluted with 0.8 ml 55°C pre-warmed QF Buffer into a new 2.0 ml Eppendorf tube. DNA was precipitated by adding 70% room-temperature isopropanol and inverting the tube 20 times, followed by incubation at room temperature for 15 min. Using a glass rod, the DNA was rolled up and placed into a new 1.5 ml Eppendorf tube containing ca. 0.8 ml ice-cold 80% ethanol. The DNA was air dried, resuspended in 200 µl autoclaved ddH_2_O, and all three extractions were pooled into a single tube, stored at 4 °C until library preparation.

DNA concentration was assessed using the Qubit Fluorometer 3.0 and the Qubit dsDNA HS Assay Kit, and the DNA purity was checked with the NanoDrop 2000 Spectrophotometer (ThermoFisher Scientific, Waltham, USA; Text S1). DNA fragment length distribution and RNA absence were checked on a 2% agarose gel via electrophoresis (50 V, 90 min) and the Quick-Load 1 kb DNA Ladder (New England Biolabs Inc., Ipswich, USA; Text S1, Fig. S1). After several dilution steps, we obtained a final DNA yield of 61.2 µg (51.0 ng/µl * 1200 µl ddH_2_O).

Library preparation was conducted using the ONT Ligation Sequencing Kit SQK- LSK110 optimized for high throughput and long reads (ONT, Oxford, UK), and applicable for singleplex gDNA sequencing. We adjusted the DNA concentration to 1000 ng in 47 µl (ca. 21.5 µl/ng) and followed the manufacturer’s instructions for library preparation (protocol vGDE_9108_v110_revL_10Nov2020, available at community.nanoporetech.com) with a few modifications. Incubation times were increased up to 15 min, the concentration of ethanol wash buffer was increased to 80%, and DNA fragments > 3 kbp were enriched using the buffers provided with the kit. Active pores ranged from ca. 800-1500. Flow cell 1 was loaded with freshly prepared libraries, flow cell 2 with one freshly prepared and one library stored at -80°C for two days, and flow cells 3-6 with two libraries stored at -80°C for 1-2 weeks. Freezing of gDNA had no apparent effect on N_50_ fragment size (see also Results & Discussion and flow cell reports on FigShare). We used a MinION 101B device, the ONT software MinKNOW v21.11.7, 21.11.9, and 22.03.5 on a local Linux system, and six R9.4.1 flow cells following the manufacturer’s instructions for priming and loading. After 1-2 days of run time, the old library was exhausted and replaced with a second library using the supplied flow cell wash kit (protocol vWFC_9120_v1_revB_08Dec2020). Sequencing was run for 3-5 days per flow cell until the majority of pores were inactive (> 99%).

Reports per flow cell can be accessed via FigShare. Basecalling was performed with the ONT software GUPPY v6.0.1 specifying the configuration file “dna_r9.4.1_450bps_fast.cfg” for fast base-calling (in preruns, the difference of median per-base quality between ‘fast’ and the substantially slower ‘high accuracy’ was quite small, i.e., 97 vs. 98%, see FastQC files on Figshare). After that, NanoFilt v2.2.0 (De Coster et al., 2018) was applied to remove the first 50 bp of the reads that were usually of low quality.

### PacBio DNA extraction and sequencing

High-molecular-weight gDNA was isolated from fresh leaf material of a single *R. cassubicifolius* s.l. individual (LH040) with the NucleoBond HMW DNA kit (Macherey Nagel, Germany), following the manufacturer’s instructions. The integrity of the DNA was determined by the Femto Pulse System (Agilent Technologies Inc, CA, USA). The DNA concentration was measured with the Qubit dsDNA High Sensitivity assay kit (Thermo Fisher Scientific, MA, USA). For the construction of a HiFi library, 7.5 µg HMW DNA was fragmented (speed 29) using the Megaruptor 3 device (Diagenode SA, Seraing, Belgium).

In total, two HiFi libraries were constructed and sequenced into one Revio 25M SMRT cell. Libraries were generated according to the ‘Procedure & Checklist - Preparing whole genome and metagenome libraries using SMRTbell® prep kit 3.0’ manual (102-166- 600, Pacific Biosciences of California Inc., USA). For size fractionation, a SageELF (Sage Science, USA) device was employed. The final sequencing library size (16.2 kb) was measured using the Femto Pulse System (Agilent Technologies Inc, USA). Polymerase-bound SMRTbell complexes were formed according to standard manufacturer’s protocols. Sequencing (HiFi CCS) was conducted using the Pacific Biosciences Revio instrument (24 h movie time, loading concentration 230 pM, 2 h pre-extension time, diffusion loading, mean insert length according to SMRT link raw data report: 18.0 kb) following standard manufacturer’s protocols. The Revio 25M SMRT cell generated 82 Gb HiFi CCS data. All steps were done at IPK Gatersleben.

HiFiAdapterFilt v2.0.0 (Sim et al., 2022) was run with default settings to convert the HiFi reads from BAM to FASTQ format, and to remove reads with remnant PacBio adapter sequences. The quality of the sequenced reads was checked by FastQC v0.11.4 (Andrews, 2010) (reports on FigShare).

### Chromosome conformation capture (Hi-C) DNA extraction and sequencing

The Hi-C sequencing library was generated from fresh leaf material of two *R. cassubicifolius* s.l. individuals (LH040) with the *DpnII* enzyme, essentially following the method described in Padmarasu et al. (2019). The library was sequenced (paired-end: 2 x 111 cycles, two indexing reads: 8 cycles, v1.5 chemistry) using the NovaSeq6000 instrument (Illumina, Inc., San Diego, USA) according to standard manufacturer’s protocols. All steps were performed at IPK Gatersleben.

### Genomic Data Analysis

#### The plastid genome

The plastome sequence was assembled using Illumina short-read data and the software GetOrganelle v1.7.5.3 (Jin et al., 2020) as recommended by Freudenthal et al. (2020), specifying the ‘embplant_pt’ database, k-mer size range between 45 and 121, and as seed the *Ranunculus repens* plastome (154,247 bp, GenBank accession number NC_036976; Dann et al., 2017). The average embplant_pt base sequencing depth was 560x. The plastome was annotated using CPGAVAS2 (Shi et al., 2019) with default settings and 2,544 reference plastomes, and visualized with CPGView (Liu et al., 2023). We checked and edited the annotation results with Geneious R11 2023.1.2 (Kearse et al., 2012). Results were also visually inspected using Bandage v0.9.0 (Wick et al., 2015). The final annotated plastome sequence (Fig. 2) was deposited in GenBank (NC_077490). We applied MAFFT v7.490 to align and investigate detailed differences between *R. cassubicifolius* and available *Ranunculus* plastome sequences. The plastid raw data (SRA: PRJNA826743) and the annotated plastome (GenBank: NC_077490) are deposited in NCBI.

**Figure 2.**
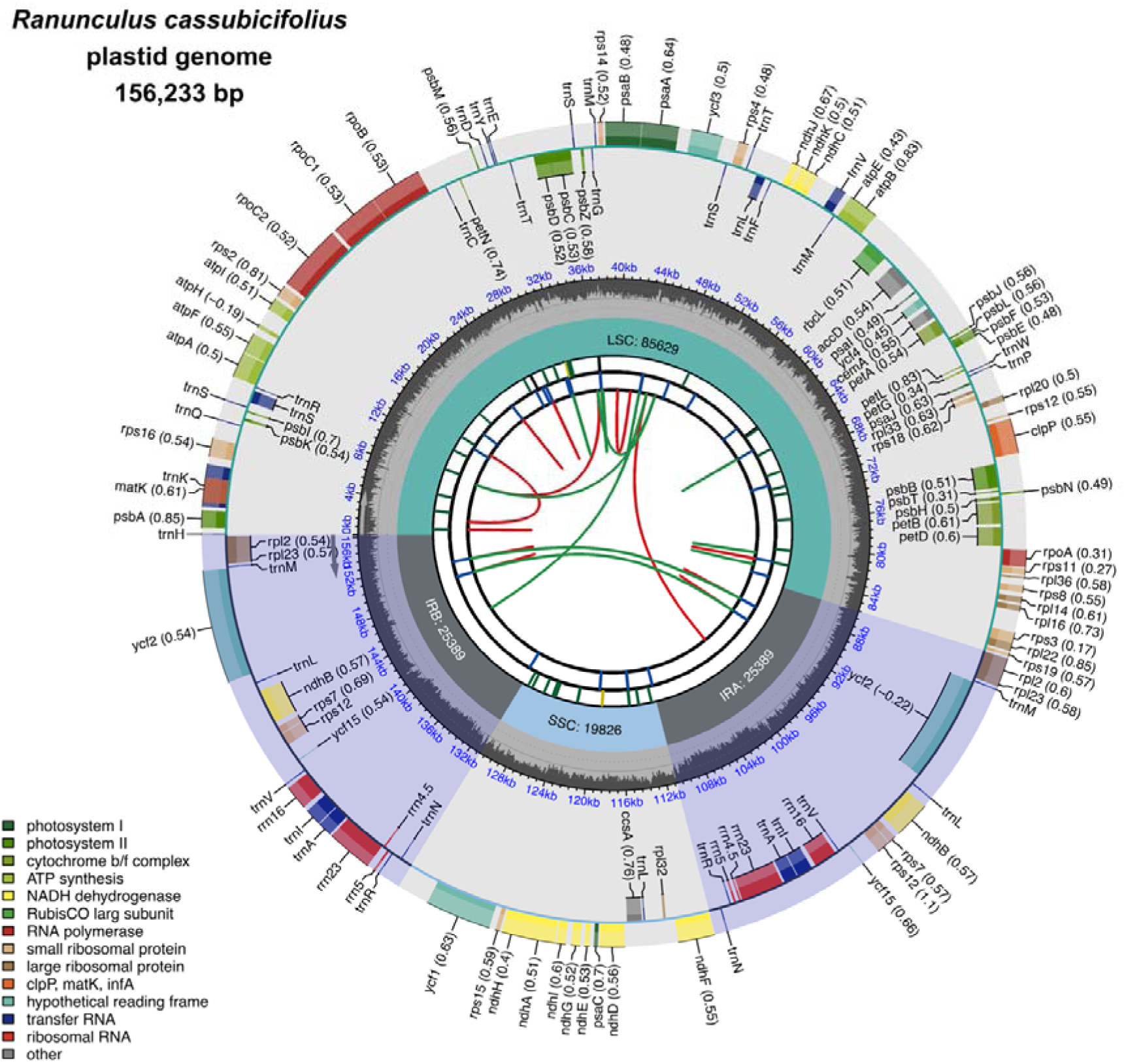
(a) The annotated plastome sequence of *Ranunculus cassubicifolius* (LH040). The plastome assembly is a 156,233 bp-long single circular contig. It contains 85 (78 unique) protein-coding genes, 37 (30 unique) tRNAs, and 8 (4 unique) rRNAs, as shown in the outer circle. Genes and t/rRNAs on the inside and outside of the circle are transcribed in the clockwise and counterclockwise directions, respectively (gene codon usage bias in brackets). The first inner circle represents GC content (%), and the second inner circle the plastome structure, i.e., a small single-copy region (SSC) and a large single-copy region (LSC) separated by two inverted repeat (IRA/B) regions. The third inner circle reflects repeat content, i.e., microsatellites (1-6 bp per repeat), tandem repeats (>6 bp per repeat), and dispersed repeats (transposable elements or satellite DNA; reports on FigShare). See Shi et al. (2019) and http://www.1kmpg.cn/cpgview/ for more details.

### The mitochondrial genome

Two alternative assembly strategies were followed. First, we used GetOrganelle with the same settings as described above, except that we specified “embplant_mt” and as seed used the closest available mitogenome (*Hepatica maxima*, GenBank accession number NC_053368.1; Park and Park, 2020). The average embplant_mt base sequencing depth (i.e., the average number of reads per position) of Illumina reads was estimated at 92x. To improve the fragmented mitogenome assembly, we first mapped ONT reads against the contigs derived from GetOrganelle and extracted the mapped reads using SAMtools v1.9 (‘faidx’ and ‘view’; Danecek et al., 2021), MINIMAP2 v2.27 (‘-ax map-ont’; Li, 2018), and BEDTools v2.29.1 (‘bamtofastq’; Quinlan and Hall, 2010). We applied MaSuRCA v4.0.9 (Zimin and Salzberg, 2022) to perform a hybrid assembly based on extracted ONT reads and “embplant_mt” hitting Illumina reads derived from the GetOrganelle assembly process.

Then, we retained an ONT-based mitogenome sequence in 18 contigs and a total length of 351,681 bp. We aligned contigs in Geneious to merge overlapping regions, which resulted in 4 contigs, onto which Illumina and ONT reads were mapped using minimap2 (‘-ax sr’ and ‘-ax map-ont’, respectively). Alignments were inspected with the Integrative Genomics Viewer (IGV) v2.16.1 (Thorvaldsdottir et al., 2013) and Geneious to ensure that previous gaps were spanned by raw reads (see alignments and information concerning contig borders on FigShare). Using Geneious, we inferred a mean Illumina depth of 62x (2.87% sites with no coverage) and ONT depth of 3640x (9.88% sites with no coverage), respectively. Remaining gaps were closed manually based on mapping results of Illumina and ONT reads. The assembly was annotated using MitoHiFi pipeline v3.2.1 (Uliano-Silva et al., 2023) including gene annotation with *H. maxima* as reference, and visualized with PMGmap (Zhang et al., 2024; http://www.1kmpg.cn/pmgmap) web browser tool. We ran RepeatModeler2 v2.0.5 to additionally check for dispersed repeats.

In a second assembly strategy, we used PacBio HiFi reads, which were subjected to the MitoHiFi all-in-one pipeline to assemble, annotate, and draw the mitogenome based on the PacBio HiFi reads. The pipeline represents a similar strategy as described before by first mapping length-filtered HiFi reads to the reference mitogenome, followed by assembly using Hifiasm and annotation with MitoFinder (Allio et al., 2020). The pipeline resulted in a very small mitogenome of about 125 kb (about 10% of the expected length). Therefore, we performed a hybrid assembly strategy with the HiFi reads as described above. This resulted in 40 raw contigs with 1,913,766 bp summarized into 9 contigs that were aimed to be arranged like the ones in the ONT hybrid mitogenome. We inferred a mean Illumina sequencing depth of 269x (1.96% sites with no coverage) and HiFi depth of 3324x (0.12% sites with no coverage). Finally, the mitogenome sequences were retained in a single contig based on ONT- and PacBio-hybrid assemblies. Due to substantial intragenomic rearrangements, we performed progressiveMauve (2015-02-25; Darling et al., 2010) instead of MAFFT in Geneious to investigate sequence differences between both assemblies, and between *R. cassubicifolius* and available Ranunculaceae assemblies.

Geneious was used to inspect and edit the annotation results. The resulting GeneBank file was converted into a feature table file for GenBank submission using the GB2SEQUIN tool (Lehwark and Greiner, 2019) within the MPI-MP CHLOROBOX webserver. Finally, we selected the PacBio-based mitogenome because of its (4x) higher sequencing depth of mapped Illumina reads and 1.5x and 82x fewer Illumina and long-read sites without coverage, although the gene content (but not the gene block arrangement, see FigShare) was the same compared to the ONT-based mitogenome. The annotated mitogenome is illustrated in Figure 3. The mitochondrial raw data (SRA: PRJNA831351) and annotated mitogenome (GenBank: PP657143) are deposited in NCBI.

**Figure 3.**
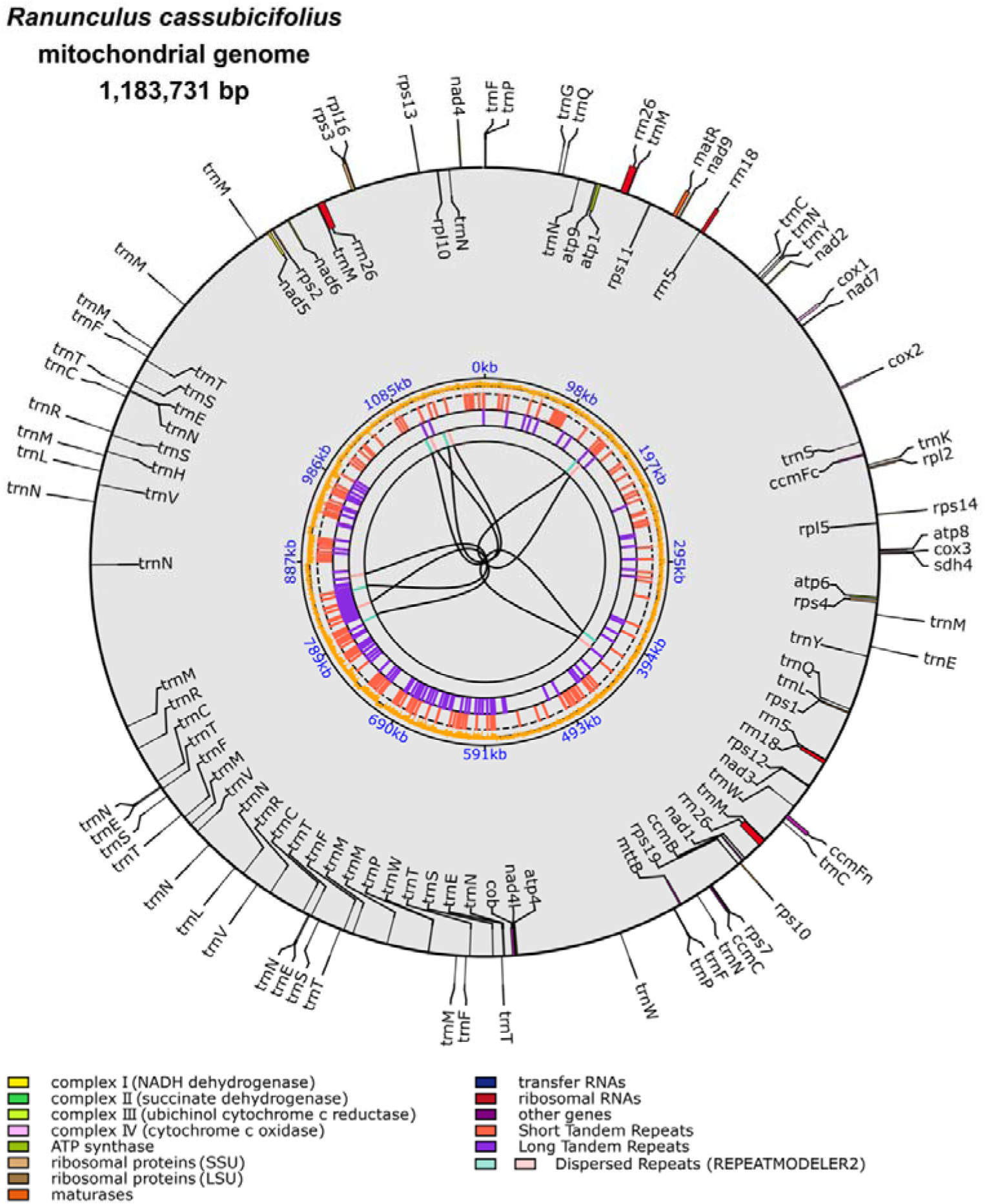
The mitogenome sequence of *Ranunculus cassubicifolius* (LH040). The mitogenome is 1,183,731 bp in size represented in a single, circular contig, and contains 40 (40 unique) protein-coding genes, 77 (unique 18) tRNAs, and 7 (unique 3) rRNAs, as shown in the outer circle. Genes and t/rRNAs on the inside and outside of the circle are transcribed in the clockwise and counterclockwise directions, respectively. The first inner circle represents GC content (%). The second inner circle reflects repeat content, i.e., short tandem repeats/microsatellites (1-6 bp per repeat), long tandem repeats (>6 bp per repeat), and dispersed repeats detected by RepeatModeler2 (reports on FigShare). See (Zhang et al., 2024) and http://www.1kmpg.cn/pmgmap for more details.

### Phylogenomics and organellar gene evolution within Ranunculacaeae

To study gene evolution within Ranunculaceae, we downloaded all available organellar genome sequences from NCBI using a custom Python script ‘findGenome.py’ (https://github.com/KK260/NCBI-Genome-Tools/). Applying the following specifications “complete genome” AND “Ranunculaceae” AND “chloroplast”/ “mitochondrion”, we found 1161 plastome and 16 mitogenome sequences on NCBI that were initially filtered by duplicate sequence removal, a maximum of 1 individual per species and 2 species per genus, and selection of the latest RefSeq ‘NC’ records. Taxon names were checked for synonyms using information from the Catalogue of Life on GBIF. This yielded 308 plastomes including 6 outgroup taxa of Berberidaceae and Papaveraceae (295 taxa), and 10 mitogenomes including 1 outgroup taxon of Papaveraceae (none available for Berberidaceae, 10 taxa) (see Table S1).

Plastome sequences were aligned using MAFFT with mode ‘FFT-NS-2’ (Katoh and Standley, 2013), following the deletion of sites with less than 50%, 70%, and 90% of available samples using Goalign v0.3.7 (Lemoine and Gascuel, 2021). We then ran ModelTest-NG v0.1.7 (Darriba et al., 2020), which determined ‘TVM+I+G4’ (unfiltered), ‘TIM1+I+G4’ (min50), and ‘GTR+I+G4’ (min70, min90) as best-fit evolutionary models for the datasets. Maximum likelihood trees were inferred with RAxML-NG v1.2.1 (Kozlov et al., 2019) specifying the estimated sequence model and 100 pseudoreplicates of non-parametric bootstrapping, and transfer bootstrap expectation (TBE) metric. To further assess branch support, Quartet Sampling (QS) v1.3.1 (Pease et al., 2018; https://www.github.com/fephyfofum/quartetsampling) scores were calculated with default settings and visualized using the supplied R script (https://github.com/ShuiyinLIU/QS_visualization). Following the approach of Karbstein et al. (2020b), the final phylogenetic tree was selected according to the highest mean TBE support and optimal QS scores (QC=1, QD=0, QI=1) using the R package PHYTOOLS v2.0-3 (Revell, 2012, 2024; https://github.com/liamrevell/phytools). Finally, we chose ‘min90’ (126 kbp) with the lowest number of gaps excluding poorly assembled non-coding DNA regions, and the highest TBE and QS values in the phylogeny as the optimal alignment (see details in Figs. S2-S4 and Text S2, and alignments on FigShare). Branches of all tribes except *Ranunculeae* were collapsed with FigTree v1.4.4 (Rambaut, 2014), gene gain and loss events per genus were mapped onto the tree (Fig. 4a).

**Figure 4.**
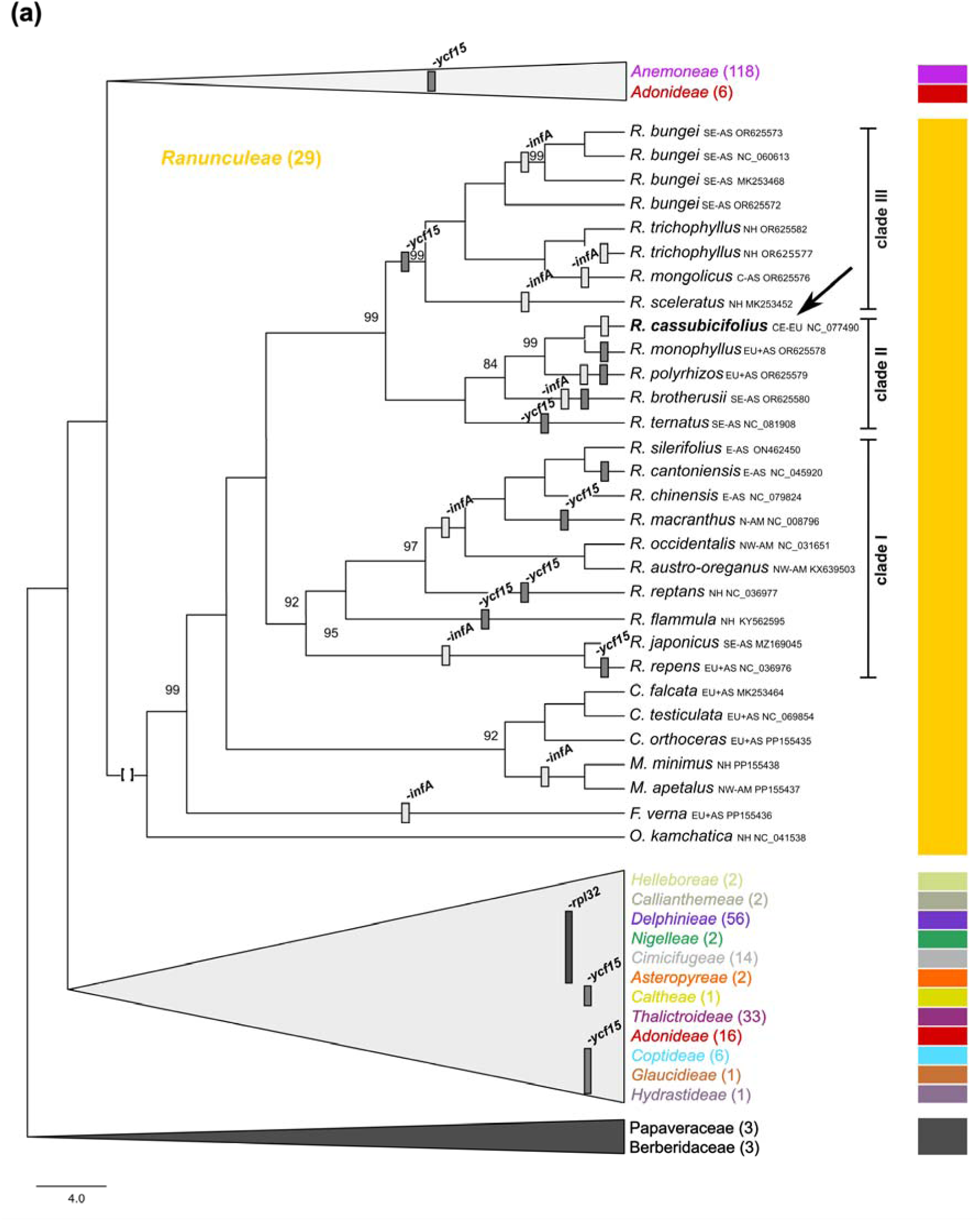
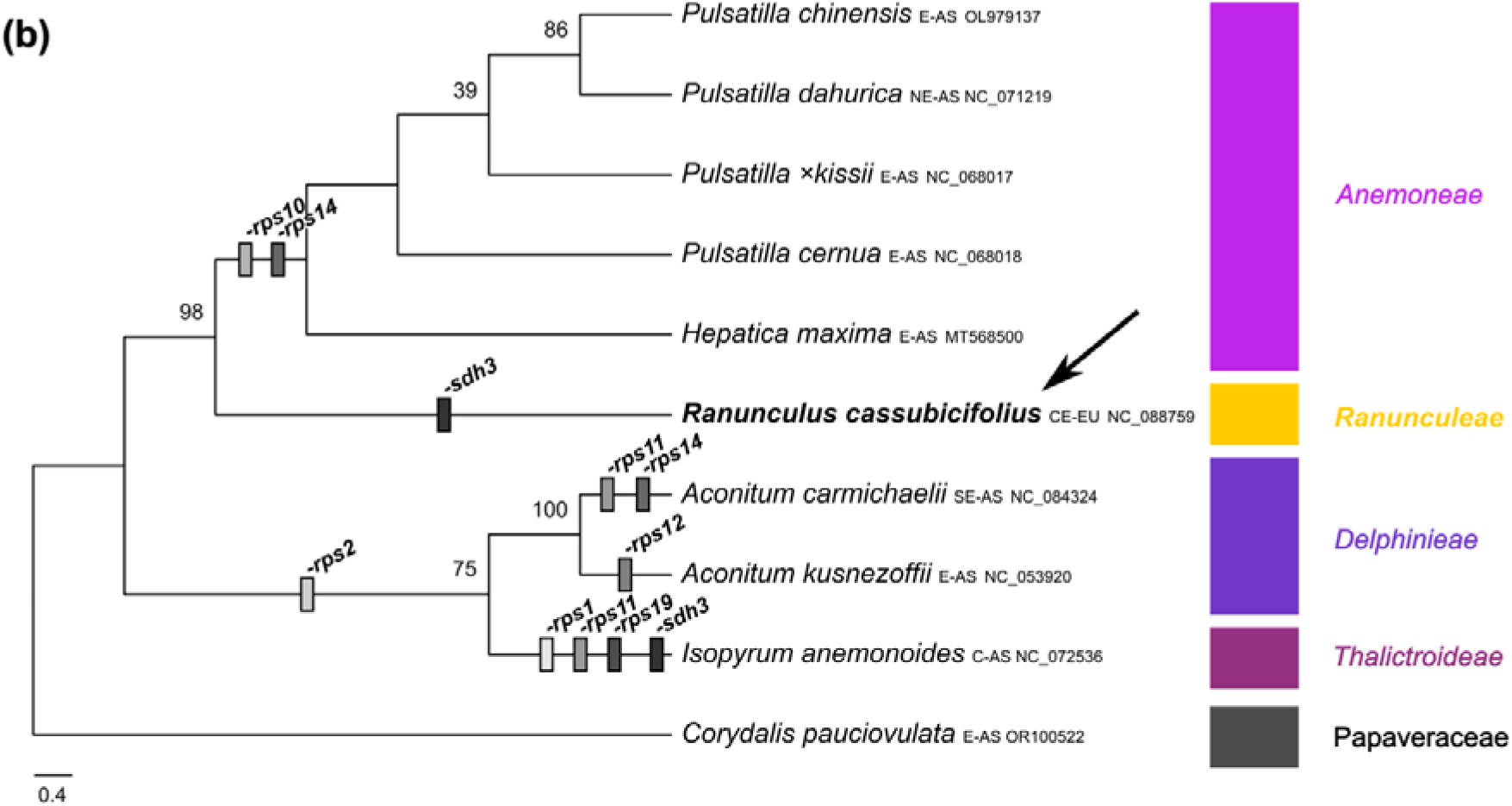
Phylogenetic relationships and gene evolution within Ranunculaceae. (a) Maximum-likelihood phylogeny based on 308 plastome sequences (295 taxa) and the min90 alignment (126 kbp) of the plant family Ranunculaceae. Branches of tribes except for *Ranunculeae* were collapsed, and gained or lost genes are shown in different colored gray bars along the tree. All branches received full (1) transfer bootstrap expectation (TBE) values, unless otherwise shown. See also Quartet Sampling (QS) metrics in Fig. S2, the complete phylogeny in Fig. S4a,b, and more technical details in Text S3. (b) Coalescent-based phylogeny of 10 mitogenome sequences and 42 genes of Ranunculaceae. All branches received full (100%) Felsenstein Bootstrap Proportions (FBP) values, unless otherwise shown. All NCBI species names were checked by gbif.org, and synonyms were replaced by accepted names (synonyms in brackets). The color of clades corresponds to tribe/subfamily names (tribal classification follows Wang et al. (2016) and Zhai et al. (2019). *Ranunculus cassubicifolius* is highlighted in bold with an arrow. Abbreviations for native distribution ranges of taxa: EU = Europe, AS = Asia, AM = America, NH = Northern Hemisphere, C = Central, E = Eastern, N= Northern, S = Southern, and W = Western.

The mitogenome phylogeny was built based on coding regions due to substantial genomic rearrangements and size differences present among sequences. Using a custom Python script ‘assembleGenes.py’ (https://github.com/KK260/NCBI-Genome-Tools/), we run Astral v5.7.8 with 100 bootstraps based on collapsed gene trees of low support previously inferred by RAxML_NG with 100 bootstraps, and mapped gene features onto the resulting tree. In Ranunculaceae, the loss of genes per group derived from the feature table of ‘assembleGenome.py’ was manually verified by mapping extracted genes from closely related Ranunculaceae species to the sequence of interest in Geneious, and edited if necessary. In *R. cassubicifolius*, we aimed to find lost organellar genes by mapping extracted genes from closely related *Ranunculus* species to the nuclear genome sequence trees (Fig. 4b).

### The nuclear genome

We applied different genome assembly approaches with default settings and a haploid genome size of 3.2 Gbp and RAM of at least 1.5 TB, unless otherwise specified: (i) using Illumina short-reads, we performed analyses with Supernova (incorporation of all reads, acceptance of extreme coverages) and SPAdes v3.13.2 (Prjibelski et al., 2020); (ii) using Illumina short-reads and ONT or PacBio/HiFi long-reads, we run SPAdes (untrusted Supernova contigs for gap filling), MaSuRCA v4.1.0 (Zimin et al., 2017), MuCHSALSA v0.02 (Gatter et al., 2021) and WENGAN (M) v0.2 (Di Genova et al., 2021); (iii a) using ONT long-reads, we performed analyses with Canu v2.1.1 (Koren et al., 2017) and Flye v2.9.1 (Kolmogorov et al., 2019) option ‘--nano-raw’), followed by the best polishing strategy of Dmitriev et al. (2021) of two-times via filtered ONT reads using Racon v1.4.3 (Vaser et al., 2017) and Medaka v0.7.1 (https://github.com/nanoporetech/medaka), and once using Illumina reads via the POLCA function (Zimin and Salzberg, 2020); (iii b) using HiFi long-reads, we run Canu, Flye, and Hifiasm v0.19.8 (Cheng et al., 2021), followed by the previous polishing strategy except for the use of polishCLR v1.1.0 (Chang et al., 2023) instead of Medaka. To ensure a fair comparison between assemblies based on ONT or PacBio reads within the same tool, we downsampled PacBio reads to 16x coverage using reformat.sh from BBTools and reran all analyses.

Statistics for genome assemblies were calculated using QUAST v5.0.1 (Gurevich et al., 2013), and are summarized in Table 1 (reports on FigShare). Assembly completeness in terms the expected single-copy genes in inherent to land plants was estimated by ‘Benchmarking Universal Single-copy Orthologs’ (BUSCO v5.0.0; Waterhouse et al., 2018; Manni et al., 2021) with the ‘embryophyta_odb10’ 2020-09-10 dataset containing 60 species and 1375 core genes. The assembly with the highest completeness, largest contig size, and best BUSCO scores (Table 1, Fig. 3, reports on FigShare) was selected for scaffolding with Hi-C data and genome annotation. Filtered Hi-C reads were mapped to the HiFi polished contigs using Juicer v1.6 (Durand et al., 2016) with default parameters. Paired reads mapping to different contigs were used for the Hi-C-associated scaffolding. We applied the 3D-DNA pipeline v170123 (Dudchenko et al., 2017) to order and orient the clustered contigs with sizes > 15,000 bp to produce chromosome-level scaffolds. Misassembles were identified and fixed manually using assembling mode in the Juicebox software, which was manually corrected based on the neighboring interactions. The resulting Hi-C contact map is shown in Fig. S5. To fulfill NCBI quality standards, we screened for non-plant contigs using FCS-GX v0.5.4 (Astashyn et al., 2024;https://github.com/ncbi/fcs) and for mitochondrial contigs using seqkit v2.8.2 (Shen et al., 2016) removing 552 short contigs in total. The functional annotation was performed using InterProScan5 (Jones et al., 2014) and eggNOG-mapper (Cantalapiedra et al., 2021) using the Funannotate “annotate” function.

**Table 1.** QUAST statistics for different genome assembly strategies of *R. cassubicifolius* (LH040) (see also reports on Figshare). SUPERNOVA, and partially SPAdes, MaSuRCA, and WENGAN used Illumina short-reads (‘Illumina’). Canu, Flye, and Hifiasm, and partially SPAdes, MaSuRCA, and WENGAN used Oxford Nanopore (ONT) or PacBio long-reads. Coverage is given per read type. Assembly polishing included the following steps: polishing via filtered ONT reads using Racon and Medaka / via filtered PacBio reads using Racon and polishCLR, respectively, and via filtered Illumina reads using POLCA (MaSuRCA pipeline). The best assembly in terms of completeness and contig length are highlighted in bold. N_50_ describes the shortest contig length required to summarize at least half (50%) of the bases of the entire assembly; L_50_ is defined as the minimum number of contigs that encompass half (50%) of the bases of the entire assembly. See Table S3 for comparisons with not downsampled PacBio reads. - means no results produced.

| Metric | SUPERNOVA<br>(Illumina22x) | SPAdes<br>(Illumina22x) | SPAdes<br>(Illumina22x +<br>ONT16x/<br>PacBio16x) | MaSuRCA<br>(Illumina22x +<br>ONT16x/<br>PacBio16x) | WENGAN<br>(Illumina22x +<br>ONT16x/<br>PacBio16x) | Canu<br>(ONT16x/<br>PacBio16x) | Canu<br>(ONT16x/<br>PacBio16x)<br>+3x Polish | Flye<br>(ONT16x/<br>PacBio16x) | Flye<br>(ONT16x/<br>PacBio16x)<br>+3x Polish | HIFIASM<br>(PacBio25x/<br>PacBio16x) | HIFIASM<br>(PacBio25x)<br>+3x Polish |
| --- | --- | --- | --- | --- | --- | --- | --- | --- | --- | --- | --- |
| **Total**<br>**assembly**<br>**length, Mbp** | 2,76<br>(86%) | 0,557<br>(17%) | 0.408/<br>(13%)/<br>0.464<br>(14%) | 0.076/<br>(2.4%)/<br>- | 0.005<br>(0.16%)/<br>1.21<br>(38%) | 0.80<br>(25%)/<br>5.05<br>(158%) | 0.81<br>(25%)/<br>5.04<br>(157%) | 3.58<br>(112%)/<br>4.90<br>(153%) | 3.45<br>(108%)/<br>4.90<br>(153%) | 3.22<br>(101%)/<br>3.62<br>(113%) | **3.21**<br>(100%) |
| **No. contigs** | 6,920,808 | 1,231,317 | 1,184,036/<br>529,574 | 38,650/<br>- | 368/<br>6,565,544 | 30,612/<br>16,262 | 32,959/<br>15,711 | 65,232/<br>12,514 | 54,310/<br>12,356 | 1,453/<br>3,449 | **1,319** |
| **No. contigs >**<br>**1 Mbp** | 562,716 | 122,407 | 110,594/<br>108,266 | 12,799/<br>- | 368/<br>5,046 | 30,612/<br>16,262 | 31,072/<br>15,711 | 64,618/<br>12,506 | 53,572/<br>12,356 | 1,453/<br>3,449 | **1,319** |
| **Largest**<br>**contig, Mbp** | 0.039 | 0.069 | 0.096/<br>0.112 | 0.145/<br>- | 0.095/<br>0.009 | 0.540/<br>14.3 | 0.542/<br>14.3 | 0.976/<br>6.57 | 0.974/<br>6.57 | 84/<br>32 | **84** |
| **GC, %** | 40 | 39 | 39/<br>39 | 40/<br>- | 40 /<br>39 | 40/<br>42 | 40/<br>42 | 43/<br>42 | 43/<br>42 | 42/<br>42 | **42** |
| **N<sub>50</sub>, Mbp** | 0.002 | 0.003 | 0.005/<br>0.005 | 0.010/<br>- | 0.021/<br>0.006 | 0.037/<br>0.617 | 0.038/<br>0.618 | 0.103/<br>0.853 | 0.104/<br>0.852 | 20.0/<br>2.89 | **20.0** |
| **L<sub>50</sub>** | 259,069 | 29,900 | 23,910/<br>23,533 | 1,507/<br>- | 76/<br>28,344 | 6,935/<br>2,374 | 6,937/<br>2,366 | 10,305/<br>1728 | 9,889/<br>1727 | 45/<br>350 | **45** |

The genome assembly was then subjected to the identification of tandem repeats (TRs) and transposable elements (TEs). Tandem Repeats Finder v4.10.1 (Benson, 1999) using parameters ‘Match=2, Mismatch=5, Delta=7, PM=80, PI=10, Minscore=5, and MaxPeriod=2000’, and EDTA v2.2.1. The EDTA pipeline integrates the results of multiple high-performing tools: LTR-FINDER (Xu and Wang, 2007; Ou and Jiang, 2019), LTR_retriever (Ou and Jiang, 2018), Generic Repeat Finder (Shi and Liang, 2019), TIR-Learner (Xiong et al., 2014; Su et al., 2019), HelitronScanner (Xiong et al., 2014; Zhang et al., 2022), and TEsorter, to achieve accurate and efficient TE identification and classification.

Given the limited availability of RNA-seq reads from *R. cassubicifolius*, we used a multi-species RNA-seq mapping approach. This strategy included RNA reads from multiple *Ranunculus* species (37 individuals, Table S1), retrieved using prefetch and fasterq-dump from NCBI SRA toolkit v3.1.0 (https://github.com/ncbi/sra-tools). RNA data was mapped onto the genome using HISAT2 v2.2.1 (Kim et al., 2019). Structural annotation of protein-coding genes used a combination of three approaches.

i. First, homology-based gene prediction using GeMoMa v1.9 (Keilwagen et al., 2016, 2018), using annotations from three Ranunculaceae and eleven Ranunculales species (Table S2). The coding genes from these 14 species were aligned to the *R. cassubicifolius* genome sequence using MMseqs2 v15-6f452 (Steinegger and Söding, 2017) following the default GeMoMa setting. GeMoMa is based on the conservation of amino acid sequences and intron position conservation among closely related species. To refine intron boundaries, RNA-seq data from five recently published diploid *R. auricomus* transcriptomes (Paetzold et al., 2022) were incorporated. The resulting fourteen gene annotation sets were merged and filtered using the GeMoMa Annotation Filter (GAF) with parameters: f="start==’M’ and stop==’*’ and (isNaN(score) or score/aa>=1.50 or (score/aa>0.8 and avgCov>50)) and avgCov>0.0 and (iAA>=0.5 or pAA>=0.5) and (evidence>1 or tpc>=0.8)" and atf="tie==1 and sumWeight>4".
ii. Second, we used the BRAKER3 pipeline (Gabriel et al., 2024), which uses ab initio gene prediction using AUGUSTUS v3.5.0 (Stanke et al., 2006, 2008) and GeneMark-ETP v1.02 (Brůna et al., 2024), which is corrected by extrinsic evidence provided by RNA-seq and protein homology. For protein evidence, we used ‘Viridiplantae’ library (Kuznetsov et al., 2023) and annotated NCBI genomes of Ranunculaceae species (*Aquilegia caerulea* (GCA_002738505.1), *Thalictrum thalictroides* (GCA_013358455.1), and *Coptis chinensis* (GCA_015680905.1)). The resulting BRAKER3 annotation was filtered using GeMoMa Annotation Filter (GAF) with parameters: f="start==’M’ and stop==’*’ and avgCov>0.0 and tpc>0.5" and atf="tie==1 or sumWeight>1".
iii. Third, we used Funannotate v1.8.15 (https://github.com/nextgenusfs/funannotate; Palmer and Stajich, 2022), initially developed for fungi but now capable of handling larger genomes. Repetitive contigs were cleaned from the genome using minimap2 v2.26 and simple repeats were masked using TANTAN (‘-s arabidopsis’; Frith, 2011). Approximately 8.04% (217 Mbp) of the genome sequence was masked. Funannotate’s gene prediction involved three steps: train, predict, and update. In all steps, the parameter ‘--max_intronlen 1000000’ was used. The filtered genome and the transcriptome sequences were used to train the annotation process by a genome-guided transcriptome assembly using Trimmomatic v0.39 (Bolger et al., 2014), Trinity v2.8.5 (Grabherr et al., 2011), HISAT2, kallisto v0.46.1 (Bray et al., 2016), and PASA v2.5.3 (Haas et al., 2008) to filter, normalize, and cluster (genome-guided) transcriptomic data. The sorted alignments, transcripts, UniProt protein library, and transcriptome annotations were the input for the gene prediction process performed by AUGUSTUS v3.5.0, GlimmerHMM (Majoros et al., 2004), and SNAP (Korf, 2004), and results were summarized with EVidenceModeler v1.1.1 (Haas et al., 2008). Additional parameters ‘--busco_db eudicots_odb10 --organism other --busco_seed_species arabidopsis’ were used in this step.

The annotations resulting from GeMoMa, BRAKER3, and Funannotate were compared based on BUSCO completeness scores. Based on these comparisons, the GeMoMa and BRAKER3 gene predictions were merged using GeMoMa Annotation Filter (GAF) with the following parameters: f="start==’M’ and stop==’*’ and avgCov>0 and tpc>0 and aa >=50 and ((isNaN(score) and (tie==1 or isNaN(tie) and tpc>0.5)) or (tie>0.5 or tpc>0.5))" and atf= " tie==1 and tpc>0.5 and (isNaN(score) or sumWeight>2)". Gene models in disagreement with RNA evidence data were fixed manually. The annotation quality was evaluated using BUSCO v5.6.1 with the embryophyta_odb10 database.

The nuclear genome sequences and annotated protein-coding genes (Fig. 6) are deposited in NCBI under SAMN44350262 in the Bioproject PRJNA831351 (until publication in FigShare). Non-coding RNA features, including rRNAs, tRNAs, and short ncRNAs were identified using Infernal v1.1.5 (Nawrocki and Eddy, 2013) cmscan, which scanned covariance models from the Rfam database v15 (Ontiveros-Palacios et al., 2024). Additionally, tRNAscan-SE v2.0.12 (Chan et al., 2021) and barrnap v0.9 (available at https://github.com/tseemann/barrnap) were used to enhance the detection of tRNAs and rRNAs, respectively.

All data analyses were performed on the HPC cluster of the GWDG (Göttingen, Germany), except for the genome annotation which was run on the de.NBI cloud.

## Results and Discussion

More than 504 million filtered Illumina reads (150 bp) were generated of *R. cassubicifolius*, resulting in 70.7 Gbp raw data and genome coverage of 22x (mean base accuracy of 99.97%), assuming a haploid genome size of 3.2 Gbp. In addition, the final ONT long-read dataset contains 51.6 Gbp and 5.83 million reads (N_50_=22.6 kbp), yielding a coverage of 16x (mean base accuracy of 96.62%). The final HiFi (PacBio) long-read dataset harbors 79.1 Gbp and 4.40 million reads (N_50_=17.9 kbp), resulting in a coverage of 25x (mean base accuracy of 99.98%).

### Plastome features and gene evolution within *Ranunculus* and Ranunculaceae

Based on short Illumina reads, we retained a circular plastid genome sequence with 156,233 bp in two states caused by the differing orientations of the inverted repeat (IR) regions (“flip-flop”; Stein et al., 1986; Walker et al., 2015). The plastome shows a typical tetrapartite structure of a small single-copy region (SSC, length: 19,826 bp) and a large single-copy region (LSC, length: 85,629 bp) separated by two IR regions (length: 2 x 25,389 bp). The plastome contains 85 (78 unique) protein-coding genes, 37 (30 unique) tRNAs, and 8 (4 unique) rRNAs, resulting in a coding content of 53.2%. We also observed 35 SSR, 32 long tandem, and 34 dispersed repeat motifs (2.2% repeat content). For example, in comparison with the closely related *R. sceleratus* (nested within the sister clade to sect. *Auricomus*) and the more distantly related *R. repens* (nested within a different subgenus within Ranunculaceae), *R. cassubicifolius* shows a similar number of unique protein-coding genes (78/77/78), tRNAs (30/28/29), and rRNAs (4/3/4). The plastome size of *R. cassubicifolius* was similar to that of *R. sceleratus* (+96 bp, 156,329 bp), but larger compared to *R. repens* (- 1,986 bp; 154,247 bp), mainly due to additional mono-, oligo-, or polynucleotide insertions between different protein-coding genes (e.g., see positions at 15,754 bp or 126,944 bp in alignment file(s) on FigShare).

Within Ranunculaceae, the plastome sequence of *R. cassubicifolius* is quite average, considering the longest (167 kbp; *Helleborus atrorubens*) and shortest (150 kbp; *Myosurus apetalus*) plastomes. A clear phylogenetic trend of size evolution in Ranunculaceae is not recognized (Fig. 4). With similar gene contents, length variation can be mainly attributed to non-coding regions (Fig. 4a, and feature tables and alignments on FigShare). However, Long et al. (2024) observed in *Myosurus* that gene length and number can also be responsible for smaller plastomes, possibly reflecting adaptation to new environments, or an effect of annual life cycles, as known from nuclear genomes (e.g., Lunkova and Ivanov, 2022). In general, compared to other flowering plant lineages, we detected usual size variation and no substantial plastome rearrangement among investigated plastome sequences of *Ranunculus* and Ranunculaceae, typical for an obligate autotrophic lifestyle (Petersen et al., 2018; Konupková et al., 2024).

The maximum likelihood plastome phylogeny with 292 taxa revealed a fully resolved and well-supported phylogeny of Ranunculaceae (TBE>85; Figs. 4 and S4, Text S3). Low branch support occurs within species-rich genera such as *Anemone, Hepatica,* or *Aconitum* and is probably due to fast radiation and the slow evolutionary rates of plastomes (Wang et al., 2016; Zhai et al., 2019). Almost all subfamilies or tribes are monophyletic, and branching is widely consistent with previous studies on plastome sequences (35 taxa) or four nuclear and plastid loci (105 taxa) (Wang et al., 2009; Zhai et al., 2019). Within *Ranunculeae*, *Oxygraphis*, *Ficaria*, *Myosuru*s, and *Ceratocephala* represent close relatives of *Ranunculus*, in agreement with Emadzade et al. (2010). Compared to Zhai et al. (2019) and Ji et al. (2023), the here presented plastome phylogeny is the most recent and taxon-rich to date of Ranunculaceae. The plastome phylogeny of *Ranunculus* widely fits the one presented in Ji et al. (2023), that is, an Eurasian-American (clade 1), European-Asian clade including *R.* sect. *Auricomus* (clade 2), and the predominant Asian clade including aquatic taxa (clade 3). The phylogeny is also largely congruent with combined plastid-nuclear marker trees in Hörandl and Emadzade (2012), but based on a smaller taxon sampling and therefore not representative for this big genus. Further plastome sequencing efforts would allow a more taxon-rich and - balanced phylogenomic reconstruction of Ranunculaceae.

Regarding gene evolution within Ranunculaceae and particularly *Ranunculus*, we frequently did not detect some specific genes (*infA*, *pbf1*, *rpl32*, *rps16*, and *ycf15*) suggesting a potential loss (Fig. 4a; see feature tables on FigShare). Gene transfers from the chloroplast to the nucleus have been reported for *infA* but also for *accD, rpl22, rpl20, rpl32, rpl23, rps7, rps16, ycf1*, and *ycf2* in seed plants (Park et al., 2015; Zhou et al., 2023). In our study, the plastid gene *rpl32* was lost in the common ancestor of *Caltheae, Asteropyreae*, *Cimifugeae,* and *Nigelleae*, and *Delphinieae* and reappeared in *Callianthemeae* and all remaining subfamilies/tribes. The ribosomal gene *rpl32* is involved in protein biosynthesis and plastid function. Its putative loss and functional transfer to the nuclear genome have been observed in other clades within Ranunculaceae (Park et al., 2015; Zhai et al., 2019; Park and Park, 2020). The gene *rpl32* may be involved in rapid adaptation to novel environments, such as in *Myosurus* adapting from mesophytic to wet habitats during climate change over the last 5 Ma (Long et al., 2024). In *R. cassubicifolius, rpl32* shows a typical length of 162 bp and is not reduced as in *Myosurus* (138 bp), or in alpine *Clematis alpina* (69 bp).

The plastid gene *infA* is present in the outgroups and all tribes of Ranunculaceae but appears to have been lost several times (22x in *Ranunculus*), including *R. cassubicifolius*. Independent losses and transfers to the nuclear genome are already documented for various angiosperm lineages, making *infA* one of the most mobile plastome genes in plants (Millen et al., 2001). In the case of *R. cassubicifolius*, we were even able to confirm the loss of *infA* in the plastome sequence and its transfer to the nuclear genome (two presumably functional, size-reduced (ca. 150-160 bp) copies exist in chromosomes 1 and 4), a result not previously reported in any other plant. The gene *infA* encodes the translation initiation factor 1 important for ribosome assembly, translation initiation, and thus cell viability (Cummings and Hershey, 1994). Transfer to the nucleus may reduce the metabolic burden of the chloroplast, enhance the functional integration and regulation of plastid- and nucleus-encoded proteins, or even escape from the accumulation of deleterious mutations in asexual organellar genomes (Muller’s Ratchet; Millen et al., 2001). Studies have shown that *infA* plays a crucial role in adaptation to cold shocks (*Escherichia coli*) or to different temperatures (slipper orchids; Giangrossi et al., 2007; Hu et al., 2022). The transfers of *infA* into the nuclear genome of *R. cassubicifolius* could thus, even more, promote the adaptation of sexual diploids, and particularly of allopolyploid apomicts distributed from Arctic to Mediterranean climates (Karbstein et al., 2020b, 2021, 2022; Hodač & Karbstein et al., 2023).

In addition, the gene *ycf15* appears to have been lost early in the evolution of Ranunculaceae (e.g., *Hydrastideae*, *Caltheae*), and regained several times in the plastome also within *Ranunculus* (12x in *Ranunculus)* including *R. cassubicifolius*. The gene *ycf15* probably originated at the roots of the angiosperms and has been pseudogenized or lost many times, and compared to other (pseudo)genes like *infA* or *rpl32*, no transfer to the nuclear genome was observed across many angiosperm lineages (Shi et al., 2013). For the genus *Ranunculus*, the complete loss of this gene had been assumed (Shi et al., 2013) but the recent study by Kim et al. (2023) for *R. austro-oreganus* and *R. occidentalis,* and the results presented here contradict this assumption. Although the gene is only 54 bp long in *R. cassubicifolius* and thus apparently degraded (e.g., compare to 198 bp in *R. austro-oreganus*), and seemingly without essential function to photosynthesis and survival of plants (but see its application for species delimitation; Shi et al., 2013; Li et al., 2021), we found two copies of *ycf15* in the IRA and IRB regions with regular ATG start and TGA stop codons, suggesting that it is functional in *R. cassubicifolius*. However, the functions of *ycf15* and the co-transcribed large plastid gene *ycf2* are still unclear, although *ycf2* and *accD* influence plastid structure and competitiveness and thus may play a role in uniparental vs. biparental inheritance of plants that is known to shape the ability of hybridization (Drescher et al., 2000; Shi et al., 2013; Liu et al., 2017; Sobanski et al., 2019; Postel and Touzet, 2020). Its role in the hybridization of sexual progenitors that produced the hundreds of allopolyploid *R. auricomus* nothotaxa (Karbstein et al., 2022) needs further investigation.

Other genes frequently known to be transferred into the nuclear genome like *accD, yc1*, or *ycf2* remain in the plastomes of Ranunculaceae. Noteworthy, we observed that annotations of (pseudo)genes are sometimes missing (e.g., *clpp*, *psbG*, *rpl22*, or *rpl32*) or renamed (*pbsN* vs. *pbf1*), which complicates plastome feature comparisons and requires additional manual work such as mapping gene sequences from closely related species to the plastome sequence of interest. Therefore, we advocate for a more careful annotation of plastome sequences through cross-species alignments of homologous genes.

### Mitogenome features and gene evolution within *Ranunculus* and Ranunculaceae

Based on the Illumina-PacBio hybrid assembly, we assembled a mitochondrial *R. cassubicifolius* genome sequence of 1,183,731 bp. It contains 40 (40 unique) protein-coding genes, 77 (18 unique) tRNAs, and 7 (3 unique) rRNAs, resulting in a coding content of 8.0%. This is consistent with observations in angiosperms (Skippington et al., 2015; Dong et al., 2018; Park and Park, 2020; Jiang et al., 2023). In comparison with *Hepatica maxima* (the closest available reference within Ranunculaceae, see Fig. S4a; Zhai et al., 2019), both mitogenomes show similar numbers of unique protein-coding genes (40/39), t-RNAs (18/18), and rRNAs (3/3), but the *R. cassubicifolius* mitogenome is slightly longer (+61,185 bp). In Ranunculaceae, variation in mitogenome size is high, ranging from 0.207 Mbp in *Isopyrum anemonoides* to 1.184 Mbp in our study species *R. cassubicifolius*. In angiosperms, mitogenomes can even range from 66 kbp to 11.7 Mbp (Skippington et al., 2015; Putintseva et al., 2020). The mitogenome of *R. cassubicifolius* is therefore relatively small, but one of the largest known for Ranunculaceae.

Size differences in the mitogenomes of Ranunculaceae can only be explained to a limited extent by variation in repeat content. In *R. cassubicifolius*, many short (187) and long (168) tandem repeats and a few dispersed repeats were detected, leading to 2.1% repeat content. This is lower compared to *H. maxima* (6.9%) and substantially lower (28x) than the smallest known mitogenome of *I. anemonoides* (0.207 Mbp, 56%). In contrast to plastomes (and animal mitogenomes), plant mitogenomes are recombinationally active, can exist as populations of alternative structures even within individuals, and are considered to be organized in noncircular molecules (Sloan, 2013; Møller et al., 2021). Sequence differences despite similar gene content are thus mainly attributable to sequence recombination and rearrangements in non-coding regions, supported by whole genome alignment results in the present study (Fig. S5a). Significant structural differences were also found between the PacBio and ONT mitogenome assemblies (Fig. S5b), which might stem from the fact that different individuals were sequenced rather than from the use of different sequencing technologies (see Table 1). In addition, despite reads bridging contig boundaries, assembly results may encourage the presence of at least 3 chromosomes instead of a single circular molecule in the active cell state (Skippington et al., 2015; Feng et al., 2021).

The coalescent-based mitogene phylogeny of Ranunculaceae was well-supported (FBP>85%; Fig. 4b), marking the first comprehensive one for the family and one of the first for plants (e.g., Lin et al., 2022). Like in the plastome tree, low branch support occurs especially within fast radiating, species-rich genera such as *Pulsatilla* (*Anemoneae*).

Branching of tribes is also widely consistent with the here presented and previously described plastome phylogenies (Fig. 4a; Wang et al., 2009; Zhai et al., 2019), that is, *Thalictroideae*, *Delphinieae*, *Ranunculeae*, and *Anemoneae*. Regarding mitochondrial gene evolution within Ranunculaceae and particularly *Ranunculus*, we observed that some genes (*rps1*, *rps2*, *rps10*, *rps11*, *rps12*, *rps14*, *rps19*, or *sdh3*) were frequently lost (Fig. 4b; see raw feature table on FigShare). In flowering plants, gene loss and transfer to the nuclear genome have been frequently observed for *rps*, *sdh3,* and *sdh4* (Adams et al., 2000, 2001; Bi et al., 2020).

In our study, the mitochondrial genes *rps10* and *rps14* appeared to be lost from *Ranunculeae* to *Anemoneae* and putatively transferred to the nuclear genome (no reference genome sequence available). The *rps* genes encode small ribosomal subunit proteins and are thus essential for mitochondrial ribosome function, as well as vegetative and reproductive plant development (Adams et al., 2000, 2001; Robles and Quesada, 2017). These two genes were shown to be under evolutionary positive selection in certain plant groups (e.g., *rps1*, *rps10, rps14* in legumes: Bi et al., 2020; *rps13* in rosids: Liu and Adams, 2008). In addition, *rps1*, *rps2*, *rps11*, and *rps19* were not found in *I. anemonoides*; *rps2* and *rps12* in *Aconitum kusnezoffii*; and *rps2*, *rps11*, and *rps14* in *A. carmichaelii*. In plant evolution, the majority of mitochondrial genes have already been transferred to the nuclear genome. This enhances regulatory host control of these genes and integration with nuclear-encoded mitochondrial proteins, and has important consequences for ATP synthesis, metabolism, and abiotic cold or heat and biotic stress responses (Adams et al., 2000; Bonen and Calixte, 2006; Woodson and Chory, 2008; Saha et al., 2017; Fakih et al., 2023).

Interestingly, only in *R. cassubicifolius,* all ribosomal genes are present. In contrast to our obligate self-incompatible species of interest (Hörandl, 2008; Karbstein et al., 2020a), all other genera examined, such as *Anemone*, *Pulsatilla*, and partially *Aconitum*, are known to be largely self-compatible (Lindell, 1998; Zhigang et al., 2006; Liao et al., 2009). Selfing (and clonal reproduction), in the absence of deleterious mutation accumulation in the mitogenome, is known to increase the transfer of mitochondrial genes to the nuclear genome and provides an adaptive advantage due to fixation of co-adapted gene combinations (Brandvain and Wade, 2009). Therefore, it would be worth investigating whether self-compatible, apomictic polyploids retain as many *rps* genes in their mitogenomes as their sexual diploid *R. auricomus* progenitors, such as *R. cassubicifolius*. However, also the putative transfer of all *rps10* and *rps14* genes to the nuclear genome within the fast radiating, species-rich clade Anemoneae (e.g., only *Anemone* and *Clematis* with >100 and >400 accepted species on gbif.org) needs further investigation. In Ranunculaceae, the lost and putatively transferred *rps* genes are known to be regulated under temperature heat and cold stress (Alafari and Abd-Elgawad, 2021; Fakih et al., 2023), and in the investigated taxa it could play a crucial role in adaptation to a broad range of subtropical to subarctic and lowland to mountainous habitats in central and (south)eastern Asia. Similar to its common forest understory ancestor, *R. cassubicifolius* grows in temperate habitats in humid forests or along stream-sides (Tomasello et al., 2020; Karbstein et al., 2021), facing low selective environmental pressure, which, along with its reproductive traits, may explain the retention of all *rps* genes in its mitogenome sequence. Whether this pattern is consistent across the 600 *Ranunculus* species with worldwide distribution in various environments, needs further investigation.

The protein-coding gene *sdh3* was detected neither in *R. cassubicifolius* nor *I. anemonoides* (see raw feature table on FigShare). It encodes the subunit 3 of the succinate dehydrogenase and plays a role in electron transfer in complex II of the respiratory chain, and is thus important for energy production (Gebert et al., 2011). Ancient and recent events of *sdh*3 loss with and without transfer to the nuclear genome have been observed in several flowering plant lineages (>40x; (Adams et al., 2001; Choi et al., 2019; Huang et al., 2019). Surprisingly, transfer of *sdh3* to the nuclear genome could not be confirmed in *R. cassubicifolius*, and for *I. anemonoides* no reference genome sequence is available. Loss of *sdh3* might reflect optimization of cellular efficiency, and that the gene is not essential in the investigated plant lineages.

### Comparison of nuclear genome assembly strategies

This study presents a systematic and comprehensive comparison of assembly strategies for large non-model plant genomes based on short- and long-read sequencing data. Related studies have mainly focused on model plants with high coverage and few tested tools (e.g., Mascher et al., 2021), or non-model plants with small-sized genomes (e.g., Murigneux et al., 2020). Here, the 18 genome assembly strategies yielded substantial differences concerning completeness, continuity, and quality (Table 1, Fig. 5). SUPERNOVA and SPAdes assemblies based on Illumina reads delivered mixed results in terms of completeness (86/17%), with too many (6.92/1.23 million contigs), short (N_50_: 2/3 kbp) contigs. BUSCO genes were also highly fragmented (F: 36/28%), with low levels of complete single-copy (S: 60/20%) and duplicated (D: 11/2%) genes, and many missing genes (M: 34/10%). This is in line with studies showing highly fragmented, gappy, low-quality assemblies of large genomes when only short Illumina reads are available (Gatter et al., 2021; Rhie et al., 2021; Zimin and Salzberg, 2022; Hotaling et al., 2023). Especially in *R. cassubicifolius* with 86% repetitive regions, short reads with moderate coverage cannot resolve these long and complex patterns.

**Figure 5.**
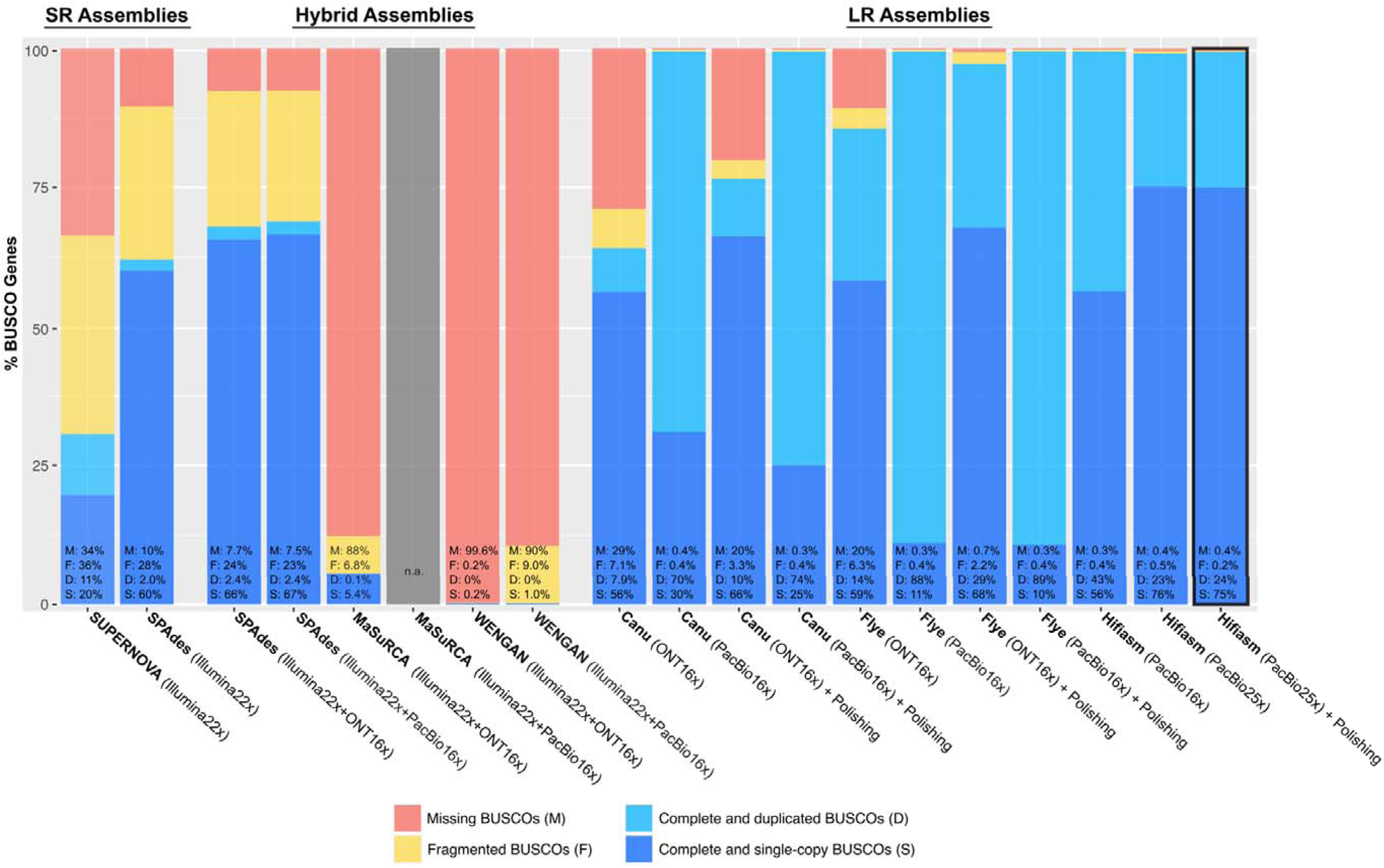
BUSCO assessments for different genome assembly strategies of *Ranunculus cassubicifolius* (LH040). SUPERNOVA, and partially SPAdes, MaSuRCA, and WENGAN used Illumina short-reads (SR). Canu, Flye, and HIFIASM, and partially SPAdes, MaSuRCA, and WENGAN used ONT/PacBio long-reads alone (LR) or in combination with short reads (Hybrid). The best assembly in terms of complete BUSCO genes (C+D) is highlighted with a black square (i.e., ‘HIFIASM (PacBio) + Polishing’). Analyses are based on 1,614 BUSCO groups searched. See the Materials & Methods section, and Table 1 and Table S4 (Excel Sheet) for more details.

Hybrid assemblies of SPAdes, MaSuRCA, and WENGAN based on Illumina and ONT or PacBio reads performed slightly better in terms of contiguity, with fewer but larger contigs (368-1.18/0.530-6.57 million contigs, with N_50_: 5-21/5-6 kbp), but not in terms of genome completeness (0.16-13/14-38%) and BUSCO completeness as many genes were often missing (M: 7.7-99.6 / 7.5-90%, F: 0.2-24 / 9-23 %, D: 0-2.4 / 0-2.4%, S: 0.2-66 / 1-67%).

SPAdes produced the best assemblies in terms of completeness, contig size, and BUSCO scores, despite being developed and tested for very small microbial genomes (<0.1 Gbp; Prjibelski et al., 2020). MaSuRCA and WENGAN (MuCHSALSA, results not shown) performed worse. Many hybrid assemblers are considered by developers to be particularly useful under low long-read coverage (<10-15x) or large/complex genomes (e.g, Zimin et al., 2017; Di Genova et al., 2021; Gatter et al., 2021). However, this cannot be confirmed in our study. These tools have been tested on model species like *Arabidopsis thaliana* with small genomes (0.135 Gbp) and very low levels of TE (ca. 21%, reviewed in Quesneville, 2020) or in human data with similar genome size (3.2 Gbp) to *R. cassubicifolius* but also lower levels of repeat content (ca. 50%, reviewed in Liao et al., 2023), or they need Illumina data at higher coverage (>30-40x) than we provide to build initial robust reads for subsequent long read mapping, which may explain their poorer performance here.

Compared to already discussed strategies, the ONT or PacBio-based assemblies conducted with Canu, Flye, and Hifiasm were generally superior in terms of completeness (25-158%), contig number (1,453- 65,232), median contig size (N_50_: 0.037-20.0 Mbp), and quality as the majority of BUSCO genes were found (M: 0.2-29%, F: 0.3-7.1%, D: 7.9-88%, S: 11-76 %). This is also consistent with the recent review by Hotaling et al. (2023), which concludes that long reads allow for more complete and contiguous genome assemblies of higher quality, particularly for large repeat-rich genomes. However, we also observed that PacBio delivered better assemblies than ONT using Canu and Flye. PacBio produced the longer (N_50_: 617/853 vs. 37-103 kbp) and fewer contigs (16,262/12,506 vs. 30,612/64,618). The lower base accuracy of ONT compared to PacBio sequence data (97 vs. 98%), despite a slightly larger median sequence length (23 vs. 18 kbp), can explain this observation (Hotaling et al., 2023). It is important to note that the ONT read N_50_ length of the runs in this study was ca. 20 kbp, while other studies have achieved runs with more than twice this value when using fresh libraries based on different DNA extraction protocols (Hakim et al., 2024; Horz et al., 2024; Nowak et al., 2024). Using the latest ONT flow cell generation and fresh DNA extracts from different kits would probably deliver much longer, higher quality reads and thus improved assemblies (see Text S5 for further discussion). However, the PacBio-based assemblies generated by Canu and Flye were far too large considering the expected haploid genome size (158/153 vs. 25/112%) and characterized by too many duplicated BUSCO genes (70/88 vs. 8/14%). This suggests uncollapsed heterozygous regions in the assembly.

However, Hifiasm – both using the full PacBio 25x coverage or the 16x downsampled set – outperformed Canu and Flye by being closest to expected genome size (101/113 vs. 158/153%), median contig length (2,890/20,000 vs. 617/853 kbp) and contig number (3449/1453 vs. 16,262/12,506), and runtime (2-3 vs. 7-14 days), but all delivered similar BUSCO gene completeness (99/99% vs. 99/99%). This agrees with recent benchmark analyses on large and complex eukaryotic and synthetic genomes (Cheng et al., 2021; Yu et al., 2024). Interestingly, our polishing strategy slightly improved assembly in terms of missing and fragmented BUSCO scores (mean M: 8.41 vs. 3.67%, and F: 2.51 vs. 1.16%), and contig number of >1 Mbp (mean 21.9 vs. 19.8 Mbp). Percentage of single-copy BUSCO genes improved significantly only for ONT-Canu assemblies (56 vs. 66%). After polishing, genome completeness (mean 118 vs. 117%) and N_50_ contig length (mean 4.13 vs. 4.13 Mbp) remained constant. We generally observed that polishing improved BUSCO statistics more for lower-quality assemblies rather than for good assemblies. Thus, sequencing errors, misassemblies, and duplications can be improved by polishing, but existing contigs are not significantly enlarged. The need for polishing thus depends on the obtained assembly quality, and is particularly attractive when working under low coverage scenarios (<30x). In addition, 25x compared to downsampled 16x coverage PacBio data delivered assemblies of substantially fewer and larger contigs (mean 26,873 vs. 21,268, and 0.832 vs. 1.83 Mbp), more single-copy BUSCO genes (mean 37.6 vs. 31.7%) and less duplicated (mean 40 vs. 46%), fragmented (mean 5.23 vs. 5.63%) and missing ones (mean 8.4 vs. 16%), but had similar genome completeneness (mean 105 vs 106%) across all tested hybrid and long-read strategies (Tables 1, S4). Coverage much below 25x is therefore not recommended for either strategy.

The PacBio-based Hifiasm assembly polished twice by PacBio and once more by Illumina reads delivered the best results among all tested strategies (100% completeness, 1,319 contigs, 20 Mbp N_50_, and 99% complete BUSCO genes). This represents the best assembly strategy for the large and complex *R. cassubicifolius* nuclear genome with low to moderate sequencing depths, and agrees with findings in smaller non-model plant assemblies (e.g., Dmitriev et al., 2021).

### The nuclear genome features

Based on the best PacBio + Hi-C assembly, we obtained a nuclear *R. cassubicifolius* genome sequence of 2,691,150,139 bp. Approximately 98.4% (2.65 Gbp) of the scaffolds were anchored to form 8 pseudochromosomes based on Hi-C data. This high-quality, chromosome-level assembly has an N_50_ of 355 Mbp, with the longest pseudochromosome spanning 447 Mbp (Fig. 6, Table 2). BUSCO analysis revealed that 94.1% (78.8% single copy, 15.3% duplicated) of the BUSCO embyophyta_odb10 genes were present, indicating the completeness of the assembly. The BUSCO score is slightly lower than before Hi-C scaffolding (99%). The Hi-C scaffolding procedure automatically removes contigs below 1,500 bp, but all contigs prior to scaffolding were far above this threshold, so contig removal does not explain the reduced BUSCO gene score. More likely are misassemblies or incorrect gene annotations based on the original assembly. The retained genome sequences excel the recommended minimum standard of the Earth Biogenome Project in terms of continuity (scaffolds >10 Mbp), functional completeness (>90% BUSCO genes), and chromosome status (>80% contigs assigned to chromosomes; https://www.earthbiogenome.org/report-on-assembly-standards, Lawniczak et al., 2022).

**Figure 6.**
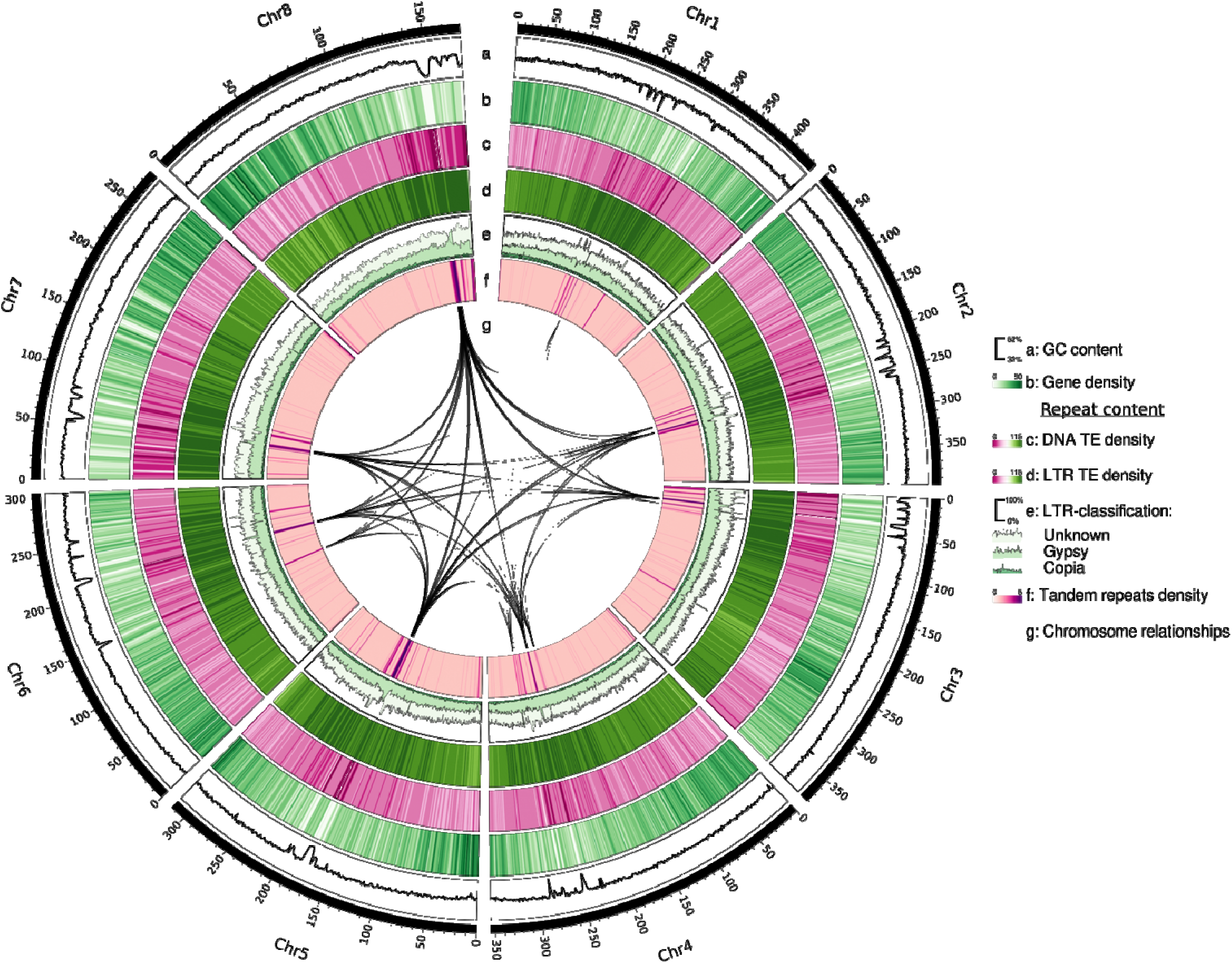
Circos plot of the chromosome-scaffolded nuclear genome of *Ranunculus cassubicifolius* (LH040). The genome is 2.69 Gbp in size, has 8 pseudochromosomes and 31,322 annotated genes. Illustrated are (from the outer to the inner circle): Chromosome number, chromosome size in Mbp, GC content (a), protein-coding gene density (b), DNA transposon element density (c), Long terminal repeat (LTR) retrotransposon density (d), percentage of unknown, Gypsy, and copia LTR-type TEs (e), tandem repeat density (f), and major interchromosomal synteny (g).

**Table 2.**
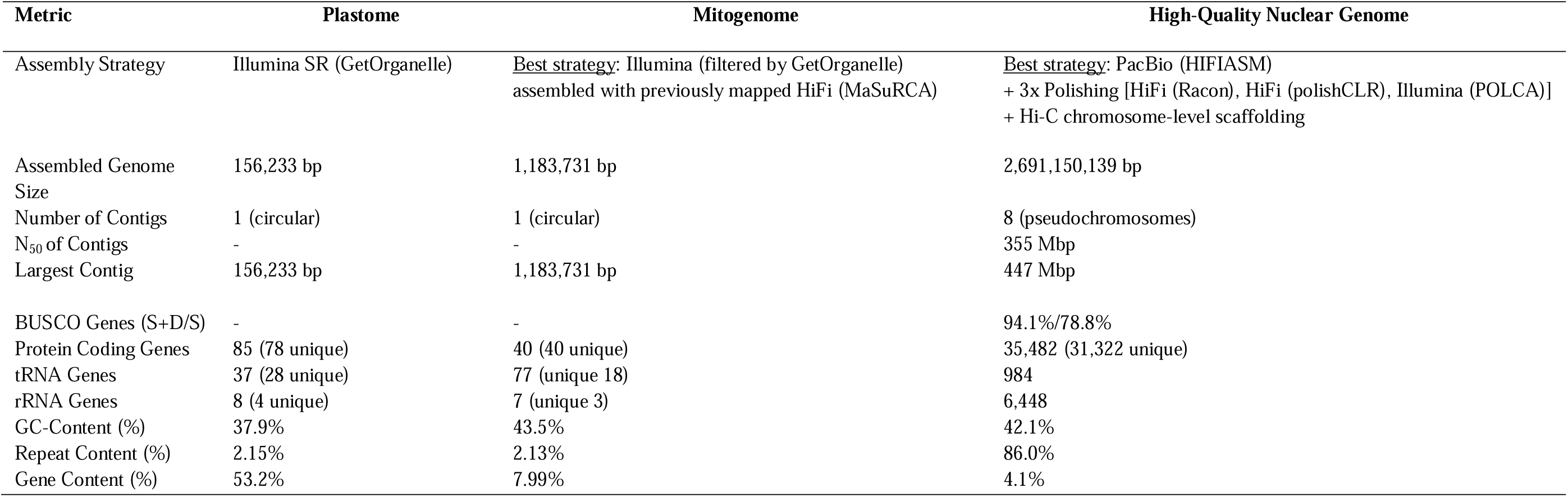
Assembly statistics of the annotated plastome, mitogenome, and chromosome-scaffolded nuclear genome of *Ranunculus cassubicifolius* (LH040). Statistics were calculated with Geneious and R. See Table 4 for a detailed repeat content of the nuclear genome, Materials and Methods, and Results section for more details.

The genome sequence contains 35,482 (31,322 unique) protein-coding genes with an average gene length of 3,548 bp and an average coding sequence (CDS) length of 1,169 bp. The gene annotation showed a high level of BUSCO completeness (94.5% using embryophyta_odb10). We predicted 984 tRNA, 99 spliceosomal RNA (snRNA), and 6,448 rRNA (3,013 5S rRNA, 1,077 5.8S rRNA, 1,154 18S rRNA, and 1,204 28S rRNAs) gene models. Further annotation details are available on Github. In comparison with *Aquilegia kansuensis* (293 Mbp; Xie et al., 2020), *Coptis chinensis* (959 Mbp; Chen et al., 2021), and *Coptis teeta* (932 Mbp; Wang et al., 2024) as the only available references within Ranunculaceae, the *R. cassubicifolius* nuclear genome is up to 10-fold larger in size (+2.42 Gbp / +1.75 Gbp /+1.78 Gbp) but has similar numbers of chromosomes (8 vs. 7/9/9). *Ranunculus*, however, has mostly the long, strongly winding chromosome type (R-type) of the family, while *Aquilegia* and *Coptis* have small, simply curved chromosome types (T-type like *Thalictrum*) (Tamura, 1993). Within *Ranunculus*, our target species shows an ancestral karyotype with large chromosomes (4 metacentric and 4 submetacentric, Baltisberger and Hörandl, 2016). Comparing all available genome sequences, *Coptis teeta* shows the most genes (43,979 in Hap1 /46,311 in Hap2), followed by *C. chinensis* (34,109), *R. cassubicifolius* (31,322), and *A. kansuensis* (25,571), suggesting that despite different gene annotation pipelines, gene content and genome size are probably decoupled (Michael, 2014; Li et al., 2017). In angiosperms, haploid genome size varies ca. 2400-fold from 0.063 to 149 Gbp (median = 0.6 / mean = 5.7 Gbp, Dodsworth et al., 2015). The nuclear genome of *R. cassubicifolius* is larger than most flowering plants, and one of the largest assembled yet for Ranunculaceae. Within diploid *Ranunculus* species, haploid genome sizes measured with karyological methods were estimated to range from 1.86 to 8.43 Gbp (Leitch et al., 2019). *R. cassubicifolius* thus represents a typical, middle-sized genome reference for the genus.

The low proportion of duplicated BUSCO genes (15%) hints at no recent polyploidization event. In ancestral angiosperms, a baseline of 12,000 – 14,000 protein-coding genes is postulated (Sterck et al., 2007). In Liu et al. (2021) and He et al. (2022), ancient whole genome duplication (WGD) events were detected in Beberidaceae and Papaveraceae as close outgroups to Ranunculaceae, and also shared and lineage-specific ones (e.g., *Coptis chinensis* with ca. 39,500 genes, Coptideae) at the base of Ranunculaceae. *Ranunculus* is nested within a far later branching clade (Ranunculeae) within Ranunculaceae, but *R. cassubicifolius* shows only slightly higher gene number. Either no further shared WGD event occurred and gene number remained quite conserved in the several million years of Ranunculaceae evolution, or re-diploidization masked another, more recent WGD event. Interestingly, the *A. kansuensis* (Thalictroideae) genome with only ca. 25,000 genes is placed shortly after *Coptis* and before *Ranunculus* in the Ranunculaceae phylogeny. Whether this re- diploidisation represents a lineage-specific or general trend followed by subsequent WGD events needs to be tested using higher quality genome assemblies and additional analysis tools (e.g., Sun et al., 2022b).

Almost 86% of the *R. cassubicifolius* assembly consists of repetitive elements (Table 4), among the highest in flowering plants (85% in *Zea mays*; Schnable et al., 2009; Wang et al., 2021). Long terminal repeat (LTR) retrotransposons are most abundant, making up to ca. 77% of the genome. Additionally, DNA transposons, and long interspersed nuclear elements (LINEs) and short interspersed nuclear elements (SINEs) retrotransposons account for 5.7%, 0.81%, and 0.2% of the genome, respectively. In angiosperms, genome size correlates well with repetitive element content and particularly with LTR-retrotransposon content but not with polyploidization (Li et al., 2017; Wang et al., 2021). Size differences between Ranunculaceae genomes can be largely explained by variation in LTR-retrotransposon contents. In *R. cassubicifolius*, LTR-RTs account for 77.6% of the genome whereas they constitute ca. 36% of the genome of *Aquilegia kansuensis* (Xie et al., 2020). In *Coptis teeta,* LTR-RTs contribute to ca. 36% of genome (Wang et al., 2024), and in *Coptis chinensis*, it is 50% (Chen et al., 2021). Interestingly, LTR retrotransposon bursts have recently been shown to have occurred during the last three million years (Pleistocene) characterized by severe climatic variability, potentially promoting adaptation through novel genomic functions and altered expression patterns (Wang et al., 2021). The progenitor species of the *R. auricomus* complex also speciated by vicariance processes due to range contraction and expansion during cold and warm phases 830-580 kya in Europe (Tomasello et al., 2020). Therefore, climatic variations and the need to adapt to them may be related to the high LTR content in *R. cassubicifolius*.

**Table 4.** Repeat content of the chromosome-scaffolded nuclear genome of *Ranunculus cassubicifolius* (LH040). Long Interspersed Nuclear Elements (LINEs), Short Interspersed Nuclear Elements (SINEs), Long Terminal Repeats (LTRs), and Terminal Inverted Repeats (TIR). A detailed explanation of repeat classes can be found in Wicker et al. (2007). Annotation reports are deposited in Github.

|  | Total Sequences | Total Length |  |
| --- | --- | --- | --- |
|  | 320 | 2,691,150,139 |  |
| Class | Count | bpMasked | %masked |
| **Class I (retrotransposons)** |  |  |  |
| **LINE** |  |  |  |
| L1 | 31,842 | 21,678,526 | 0.81% |
| **LTR** |  |  |  |
| Copia | 147,394 | 80,810,425 | 3.00% |
| Gypsy | 1,105,626 | 654,391,560 | 24.32% |
| unknown | 2,075,881 | 1,353,935,235 | 50.32% |
| **SINE** |  |  |  |
| tRNA | 309 | 114,573 | 0.00% |
| unknown | 347 | 444,295 | 0.02% |
| **Class II (DNA transposons)** |  |  |  |
| **TIR** |  |  |  |
| CACTA | 85,851 | 32,242,719 | 1.20% |
| Mutator | 134,007 | 38,209,632 | 1.42% |
| PIF_Harbinger | 86,910 | 30,275,684 | 1.13% |
| Tc1_Mariner | 32,184 | 12,038,336 | 0.45% |
| hAT | 133,119 | 40,436,252 | 1.50% |
| **nonTIR** |  |  |  |
| helitron | 123,174 | 38,735,549 | 1.44% |
| **rDNA** |  |  |  |
| 45S | 191 | 42,204 | 0.00% |
| **Repeat Fragment** | 44,541 | 11,822,897 | 0.44% |
| **Satellite DNA** | 186 | 83,887 | 0.00% |
| total interspersed | 4,001,562 | 2,315,261,774 | 86.04% |
| **Total** | **4,001,562** | **2,315,261,774** | **86.04%** |

Given the overall high completeness of our assembly (94.1% of BUSCO genes), we think it could represent a very important resource for exploring the genetic background of reproductive mode shifts from sex to apomixis in *R. cassubicifolius* and beyond. The apomixis candidate genes under selection in apomictic hybrids (*ASY1, XRI1*, and *APC1*) recognized by Paetzold et al., 2022) based on RNA-seq data were also detected in the genome of *R. cassubicifolius* at pseudochromosomes 2 and 5. However, the apomixis candidate gene *MSP1* was not found and needs further investigation. The annotated genes will be also an important resource for comprehensive analysis of loci under selection in the genus (e.g., via dN/dS ratios; (Pellino et al., 2013), which could give insights into general genome evolution, adaptation, and speciation.

### Conclusions

In this study, we assembled the complete plastome, mitogenome, and large nuclear genome sequences of *R. cassubicifolius,* a diploid sexual progenitor species within the *Ranunculus auricomus* complex, using different combinations of Illumina short-reads and ONT and PacBio long-reads under low to medium sequencing depth scenarios. We generated a plastid and the first intricate mitogenome sequences for *Ranunculus* based on Illumina and Illumina + PacBio data, respectively. An updated plastome phylogeny and the first mitogenome phylogeny of Ranunculaceae indicated frequent gene losses (e.g., *infA*, *ycf15*, and *rps* genes), potentially in the context of environmental adaptation and reproductive features. To assemble the large and complex nuclear genome, we performed 18 assemblies using short-, long-, and hybrid-read strategies, followed by gap filling and/or polishing with various state-of-the-art bioinformatic tools. The best result in terms of contiguity and completeness was achieved by Hifiasm using PacBio data, polished twice with filtered PacBio reads (Racon, polishCLR) and once with Illumina reads (POLCA), representing the first high-quality, chromosome-scaffolded genome sequence within *Ranunculus* and the fourth within Ranunculaceae (although the two currently available genome sequences on NCBI are below 1 Gbp genome size; Xie et al., 2020b; Chen et al., 2021). The large *R. cassubicifolius* genome with 2.69 Gb size, containing 31,322 annotated genes and 86% repetitive elements, represents a typical medium-sized plant genome and is probably representative for *Ranunculus*.

The obtained results and in particular the new nuclear genome assembly are useful for (i) improving read mapping (see Text S4), orthologous gene/locus assembly, and allele phasing of subgenomic reduced-representation sequencing (RRS) (e.g., RAD-Seq, target enrichment), transcriptome (RNA-seq), and whole genome (re)sequencing data (e.g., WGS, WGR), (ii) as a basis for understanding cyto-nuclear compatibility for both plastid-nuclear and mitochondrial-nuclear interactions; (iii) as diploid reference for the analysis of the more complex allo- and autopolyploid nuclear genomes within the *Ranunculus,* (iii) as reference for further studies of genome evolution, loci under selection, adaptation, and speciation, as well as (iv) for improving functional analyses (e.g., gene expression studies) related to apomixis, adaptation, and many other traits.

## Data Availability

The basic data supporting the findings are available within the manuscript and Supporting Information. Raw sequencing data and the assembled sequences are deposited at the National Center for Biotechnology Information (NCBI) database under the BioProject accession numbers: PRJNA826743 for Illumina raw reads (https://www.ncbi.nlm.nih.gov/bioproject?LinkName=biosample_bioproject&from_uid=2758 <u>2367</u>), PRJNA831351 for filtered ONT, PacBio, and Hi-C reads, filtered ONT and PacBio mitochondrial reads, and filtered mitochondrial Illumina reads (https://www.ncbi.nlm.nih.gov/bioproject/PRJNA831351/); SAMN44350262 the assembled and annotated genome (https://www.ncbi.nlm.nih.gov/datasets/genome/SAMN44350262/); NC_077490 for the assembled and annotated plastome (https://www.ncbi.nlm.nih.gov/search/all/?term=NC_077490), PP657143 for the assembled and annotated mitogenome (https://www.ncbi.nlm.nih.gov/search/all/?term=PP657143). The non-coding genes and transposable elements annotation are available on Github at https://github.com/NancyChoudhary28/Ranunculus-genomics. We deposited software assembly reports, statistical results, and genome alignments on FigShare (https://doi.org/10.6084/m9.figshare.20488908; private link: https://figshare.com/s/b7fba31a6f2444123906), and the used Python scripts on GitHub at https://github.com/KK260/NCBI-Genome-Tools.

## Author Contributions

KK and EH designed research; EH collected plant materials; KK, AH, MP, ST, JPB, and BHB performed lab work; KK, NC, XT, BP, and ST analyzed data; KK and EH wrote the paper, with contributions from NC, II, BP, JdV, XT, AH, NS, ST, AP, NW, BHB, JPB, CP, and MP.

## Acknowledgments

We acknowledge the excellent technical assistance of Susanne König, Sylvia Swetik, and Manuela Knauft during PacBio and Hi-C library preparation and DNA sequencing, Jennifer Krüger for laboratory help during ONT DNA sequencing, Silvia Friedrichs and Gabriele Ließmann for garden work, Ladislav Hodač for material sampling and field image supply, and Magnus Nordborg and Robin Burns for bioinformatic advice. This work was supported by the BMBF-funded de.NBI Cloud within the German Network for Bioinformatics Infrastructure (de.NBI) (031A532B, 031A533A, 031A533B, 031A534A, 031A535A, 031A537A, 031A537B, 031A537C, 031A537D, 031A538A).

## Supplementary Material

The Supplementary Material for this article can be found attached to this article.

