## Supporting Information for "Efficient assembly of plant genomes: A case study with evolutionary implications in *Ranunculus* (Ranunculaceae)"

Kevin Karbstein 0000-0003-1424-6557

Nancy Choudhary 0000-0002-2562-7905

Ting Xie 0000-0002-6648-7697

Salvatore Tomasello 0000-0001-5201-1156

Natascha D. Wagner 0000-0001-6623-7623

Birthe H. Barke 0000-0001-5379-6055

Claudia Paetzold 0000-0002-4128-6645

John P. Bradican 0000-0001-5650-6172

Axel Himmelbach 0000-0001-7338-0946

Nils Stein 0000-0003-3011-8731

Argyris Papantonis 0000-0001-7551-1073

Iker Irisarri 0000-0002-3628-1137

Jan de Vries 0000-0003-3507-5195

Boras Pucker 0000-0002-3321-7471

Elvira Hörandl 0000-0002-7600-1128

**Text S1.** Extraction of genomic DNA (gDNA) for Oxford Nanopore Technology (ONT) sequencing performed at the University of Göttingen. We have included information on subsequent DNA quantity and quality control.

**gDNA extraction** (Qiagen Genomic Tip 20/G, high-molecular-weight DNA up to 150 kb; Qiagen, Hilden, Germany)

**Protocol based on**

- https://www.qiagen.com/de/resources/download.aspx?id=cb2ac658-8d66-43f0-968e-7bb0ea2c402a&lang=en
- the manufacturer´s instructions of the Qiagen Genomic DNA Buffer Set
- Vaillancourt and Buell (2019), Li et al. (2020)
- own experience

1) **Harvest young(!), fresh, dark-adapted leaves from your samples** (young expanding leaf and/or shoot material is optimal and should be dark-treated for 12 ~ 24 h before harvesting to reduce photosynthetic by-products; use a Falcon tube wrapped in aluminum foil for fresh leaf sampling (see also Li et al., 2020)
2) **Place autoclaved mortar and pestle (one set per sample) in the freezer just prior to gDNA extraction.** This avoids immediate melting of ground frozen leaf powder and ensures high quality of extracted DNA (see below).
3) **Be sure to wear a lab coat, safety glasses, and gloves for any subsequent lab work!**
4) **Finally (!) grind 75 mg fresh leaf material** (should result in **at least** 30 ng/µl in 200 µl dd$H_2O$, e.g., Qiagen AE buffer leads to bad Nanodrop 260/230 values due to various ingredients, like e.g. salt, which can be problematic in subsequent DNA sequencing; → 30 ng/µl * 200 µl= 6000 ng → 6 µg → Genomic-tip 20/G: up to 20 µg) **using liquid nitrogen and autoclaved mortar and pestle. The amount of fresh material is species-specific,** and 75 mg is needed to recover 6 µg of DNA from *Ranunculus auricomus* plants. **Pour a little liquid nitrogen into the empty mortar** and wait for it to evaporate (to cool), then add a little liquid nitrogen and then the leaves and **start grinding immediately**. Repeat this step if necessary. In the end, you should have a **finely ground powder that is still frozen**.
5) **Immediately(!) transfer the leaf powder** with a spatula into a 2.0 ml Eppendorf tube containing 400 µl of pre-warmed (40°C) Quiagen AP1 Extraction Buffer.

6) Incubate for 1.5 h at 40°C with gentle agitation (ca. 350 rpm).
7) Add **4 µl DNase-free Qiagen RNase A** (100 mg/ml), invert 20 times, and incubate for 30 minutes at 37°C.
8) Add **18 µl Qiagen Proteinase K** (20 mg/ml), invert 20 times, and incubate for 2 h at 50°C with gentle agitation.
9) **Centrifuge at 12,000 x g for 20 minutes** to pellet insoluble debris.
10) **Be sure to use wide-bore pipettes** to avoid disrupting the DNA molecules!
11) **Equilibrate the Qiagen Genomic Tip 20/G with 1 ml of QTB Buffer,** and allow the Qiagen Genomic Tip to empty by gravity flow.
12) Transfer **400 µl of the clarified lysate** (do not include the foam that formed during incubation time) to the appropriate buffer **QBT-equilibrated Qiagen Genomic Tip 20/G**. Allow it to enter the resin (Granulat, Harz) by gravity flow. **Important note: Keep the residue!** If you use too much fresh material, it is possible that the DNA won't be retained by the tip filters. You can recognize this problem in Step 15 because there is no precipitate. Just use the residue from this step and continue with step 15. Since you are transferring only precipitated DNA to your glass tube, no contaminants will be transferred to the next tube.
13) Wash with **QC Buffer** (Genomic-tip 20/G: **5 x 1 ml) (Important Note:** Keep the residue as well!).
14) Elute with prewarmed (55°C) **Buffer QF** (Genomic-tip 20/G: **0.8 ml**) into a new 2.0 ml Eppendorf Tube.
15) **Precipitate the DNA** by adding 0.7 volumes (ca. 0.5 ml) of room temperature isopropanol by inverting the tube 20 times, and waiting 15 minutes (DNA should usually precipitate immediately; **if this is not the case, the DNA was already solubilized in the previous steps → see comments above**).
16) **Spool out DNA on a glass rod** (Alternatively: centrifuge at 5,500 g at 4°C for 30 minutes, and remove isopropanol).
17) Prepare a 1.5 ml Eppendorf tube with ca. 0.8 ml ice-cold 80% ethanol, and **wash the DNA (will be kept on the glass rod).**
18) **Air-dry the DNA (30 seconds) and gently resuspend the DNA in 200 µl Qiagen TE Buffer overnight in the refrigerator (4 °C**) (Alternatively, gently pipette the solution up and down several times 4-5 hours after extraction).
19) **Store at 4°C in the refrigerator for** several weeks or months. **Freeze the DNA sample only in case of long-term storage** (Freezing → DNA degradation!).

**Nanodrop (DNA Quality)**

(1) Turn on the computer and click on the NanoDrop 2000 Spectrophotometer icon on the desktop.
(2) Click on the Nucleic Acid tab.
(3) Blank the machine before measuring concentrations (e.g., with 1.5 µl ddH2O/AE water).
(4) Add 1.5 µl sample volume (shake very gently before).

**Gel electrophoresis (DNA Quality and Quantity)**

**Gel preparation:**

(1) Weigh 1 g agarose and add to 70 mL 1 x TAE buffer (2%).
(2) Boil (microwave).
(3) Cool under running water.
(4) Pour the gel solution into the gel stand and remove bubbles.
(5) Place combs on the gel stand.
(6) Place the hardened gel (after approx. 30 minutes) with the gel slide in the gel chamber.

**Sample preparation:**

(1) Mix 5 µl of the extracted DNA with 1 µl application buffer (6x ROTI Load DNAstain; Carl Roth, Germany), respectively.

**Gel electrophoresis:**

(1) Pipette samples into the gel chambers, and note the sample order.
(2) Pipette 3 µl DNA size standard (Quick-Load 1 kb Extended DNA Ladder BioLabs) mixed with 1 µl loading buffer (6x ROTI Load DNAstain) into an empty chamber.
(3) Gel run: 50 V, 100 minutes.
(4) Take an image of the gel under UV light.

**Qubit (DNA Quantity;** Thermo Fisher Scientific, Waltham, USA)

**Sample preparation:**

(1) All reagents should be at room temperature!
(2) "Working solution" for MasterMix: **per sample and standard measurement, 199 µl** dsDNA HS Buffer + **1 µl** dsDNA HS („high sensitive") / BR ("broad range") Reagent.
(3) Qubit tubes: pipette **2 µl** DNA extract into the tube (vortex DNA before!) and add **198 µl** HS/BR Buffer → immediately vortex!
(4) Pipette **10 µl** S1 and S2 HS/BR standard into a tube and add **190 µl** HS/BR Buffer → immediately vortex!
(5) Incubate all qubit tubes protected from light for **at least 5 minutes**.

**Qubit measurements:**

(1) Set DS-DNA (high sensitive/broad range) in the Qubit Fluorometer
(2) Calibration: first measure Standard S1, then measure Standard S2
(3) Sample measurements (invert tube 2-3 times before measurement)

**Text S2**. Determining the optimal DNA sequence alignment for phylogenetic analyses.

Filtering settings (i.e., the minimum number of samples per alignment site) resulted in different alignment lengths: 343,436 bp (min0), 155,794 (min50), 147,062 (min70), and 126,707 bp (min90). With increasing strictness of data filtering (min0 to min90), we observed an increase in mean FBP (86.09, 88.06, 88.31, and 91.21) and TBE (93.43, 94.36, 93.92, and 95.56) values. As expected for large phylogenies, TBE was higher than FBP (88.42 vs. 94.32). Mean values of QS metrics such as QC (0.66, 0.67, 0.66, and 0.67) and QI (0.90, 0.91, 0.91, and 0.90) showed relatively high levels of concordant patterns with taxa usually “placed” correctly within the phylogeny and high branch informativeness across filtering settings, although a drop in median QC for min0 (QC=0.92, all others with QC and QI = 1) was observed. The raw (min0) and less strictly filtered alignments (min50, min70) contained various gaps due to substantial variation in non-coding plastome regions. While this information can be useful for discriminating closely related taxa, it has also been shown that random or too high levels of missing data negatively affect phylogenetic reconstructions and bootstrap values (Eaton et al., 2017; Karbstein et al., 2020). Therefore, we chose the min90 setting (126,707 bp) with the lowest number of gaps in the alignment, excluding poorly assembled non-coding DNA regions, and the highest average bootstrap support in the phylogeny.

**Text S3**. Detailed results of plastome-based phylogenies in Ranunculaceae.

The results of the best maximum likelihood plastome analyses indicate a fully resolved and well supported phylogeny (TBE>95; Fig. S4a,b). Low branch support occurs particularly in the species-rich genera *Anemone, Clematis, Hepatica,* or *Aconitum* probably due to fast radiation and slow rates of plastome sequence evolution (Wang et al., 2016; Zhai et al., 2019). Almost all subfamilies/tribes are monophyletic (except for *Anemoneae* and *Adonideae*; also a few species non-monophyletic like *Clematis fruticosa* or *terniflora*). Convergent evolution with multiple origins of taxonomically informative morphological characters has been shown for Ranunculaceae (Zhai et al., 2019) and may explain non-monophyletic relationships observed particularly for *Adonideae*. The ambiguous position of the local endemic of the Qinling Mountains in China *Anemone taipaiensis*, which leads to the non-monophyly of *Anemoneae* (but monophyletic in less well-supported min50/70 phylogenies, Fig. S4b,c), needs further investigation.

The tribes branch successively as follows: *Glaucideae*, *Hydrastideae*, and *Coptideae* (i.e., core Ranunculaceae; Wang et al., 2009, 2016), *Adonideae* (incl. *Trollius* and *Calathodes*) as sister to *Thalictroideae*, *Caltheae* as sister to *Asteropyreae*, *Cimifugeae*, *Nigelleae* as sister to *Delphinieae*, *Calanthemeae*, *Helleboreae*, *Ranunculeae*, and *Adonideae* (incl. *Adonis*) as sister to *Anemoneae*. Observed phylogenetic patterns among tribes are largely consistent across min0 to min90 datasets (except for 'jumping' tribes such as *Cimifugeae* and a part of *Thalictroideae*; Fig. S4a-d), indicating a robust phylogenetic framework for Ranunculaceae. Branching of tribes is also widely consistent with a recent plastome study based on fewer taxa or a multi-locus-based nuclear and plastid phylogeny (Wang et al., 2009; Zhai et al., 2019; except for non-monophyly of *Adonideae* and sister relationships among "central" tribes such as *Delphinieae*, *Nigelleae, Cimifugeae, Caltheae*, and *Thalictroideae*).

**Text S4**. Mapping off-target reads the differently related reference plastomes.

Mapping target enrichment (TEG) off-target reads to reference plastome sequences from sampled or herbarium specimens representy a common strategy for constructing phylogenies in evolutionary studies (Folk et al., 2015; Tomasello et al., 2020). Therefore, we followed the procedures described in the study by Karbstein et al. (2022) to examine the effect of read mapping and lengths of the resulting alignments using the plastome sequences of the distantly related *R. repens*, the *R. cassubicifolius* as a closely related reference, and 87 di- and polyploid, sexual and apomictic *R. auricomus* TEG samples (SRA BioProject PRJNA628081; Tomasello et al., 2020; Karbstein et al., 2022).

Mapping off-target plastid reads against the coding regions of both references yielded slightly better assemblies for the closely-related *R. cassubicifolius* with 24.30% compared to *R. repens* with 24.07% plastome sequence completeness (assembly lengths) resulting in 827 bp longer alignments on average (37,968 vs. 37,141 bp; see plastome mapping reports on FigShare). Read mapping to the closest possible reference ensures reliable locus assembly, particularly in non-coding regions often characterized by insertions and deletions even among individuals of a species (Freeland et al., 2011; Folk et al., 2015; Scheunert et al., 2020). This might be even more effective when no close reference plastome sequence is available within the genus, with the potential to improve subsequent phylogenomic analysis due to increased sequence length and information content.

**Text S5.** Impact of using frozen libraries for ONT DNA sequencing.

Sequencing of ONT libraries of flow cells 3-6 once stored at -80°C compared to fresh libraries of flow cell 1 (and partly 2) did not yield decreased output quality in terms of $N_{50}$ read fragment size (flow cells 1-6: 20.1, 21.67, 21.6, 20.5, 19.6, and 23.3 kbp), and read number and sequencing output (flow cells 1-6: 1.1, 0.7, 0.9, 0.5, 1.8, and 0.7 M / 10.8, 6.6, 7.3, 3.3, 11.7, and 5.7 Gbp) (see flow cell reports on FigShare). Therefore, short freezing and thawing of the DNA libraries did not lead to any apparent fragmentation of the DNA reads, supported by observations for dried material or general recommendations of other studies (Doyle and Dickson, 1987; Russo et al., 2022). This may be attributed to the used standard elution buffer that stabilizes DNA in solution, and the rapid freezing at -80°C and slow thawing of libraries.

**Table S1.** Selected (a) plastome and (b) mitogenome sequences from NCBI.

**(a)**

| Genbank ID | Taxon | Sequence Length (bp) |
|---|---|---|
| PP407389 | *Aconitum alboviolaceum* | 155652 |
| NC_036357 | *Aconitum angustius* | 156109 |
| KY407559 | *Aconitum austrokoreense* | 155892 |
| MK253470 | *Aconitum barbatum* | 156761 |
| NC_041579 | *Aconitum brachypodum* | 155651 |
| OK323949 | *Aconitum bulleyanum* | 155791 |
| KY407560 | *Aconitum carmichaelii* | 155880 |
| NC_031420 | *Aconitum ciliare* | 155832 |
| MG678803 | *Aconitum contortum* | 155653 |
| NC_031421 | *Aconitum coreanum* | 157029 |
| OM289058 | *Aconitum delavayi* | 155733 |
| NC_061703 | *Aconitum duclouxii* | 155479 |
| NC_038096 | *Aconitum episcopale* | 151214 |
| NC_036358 | *Aconitum finetianum* | 155625 |
| NC_056280 | *Aconitum flavum* | 155654 |
| MZ959044 | *Aconitum forrestii* | 155869 |
| OK539525 | *Aconitum habaense* | 155800 |
| NC_038095 | *Aconitum hemsleyanum* | 155684 |
| KT820669 | *Aconitum jaluense subsp. jaluense* | 155926 |
| KT820670 | *Aconitum japonicum subsp. napiforme* | 155878 |
| MK253471 | *Aconitum kusnezoffii* | 155832 |
| NC_035894 | *Aconitum longecassidatum* | 155524 |
| NC_080244 | *Aconitum macrorhynchum* | 155913 |
| NC_031423 | *Aconitum monanthum* | 155688 |
| NC_061321 | *Aconitum nagarum* | 155732 |
| NC_061553 | *Aconitum ouvrardianum* | 155799 |
| MK782812 | *Aconitum paniculigerum var. wulingense* | 155774 |
| MK782813 | *Aconitum paniculigerum var. wulingense* | 155813 |
| NC_053848 | *Aconitum pendulum* | 155597 |
| NC_058692 | *Aconitum piepunense* | 155836 |
| NC_035892 | *Aconitum pseudolaeve* | 155628 |
| MN967020 | *Aconitum puchonroenicum* | 155631 |
| NC_057536 | *Aconitum quelpaertense* | 155636 |
| NC_061700 | *Aconitum ramulosum* | 155841 |
| MF186593 | *Aconitum reclinatum* | 157354 |
| MW817090 | *Aconitum scaposum* | 157688 |
| PP407390 | *Aconitum sczukinii* | 155819 |
| NC_036359 | *Aconitum sinomontanum* | 157215 |
| NC_061555 | *Aconitum stapfianum* | 155858 |
| NC_061704 | *Aconitum stylosum* | 155475 |
| NC_050689 | *Aconitum tanguticum* | 157114 |
| ON751949 | *Aconitum transsectum* | 155872 |
| NC_066973 | *Aconitum tschangbaischanense* | 155881 |
| NC_072898 | *Aconitum umbrosum* | 157227 |
| NC_038094 | *Aconitum vilmorinianum* | 155761 |
| KU556690 | *Aconitum volubile* | 155872 |
| NC_061702 | *Aconitum weixiense* | 155872 |
| NC_041525 | *Actaea asiatica* | 159638 |
| NC_081474 | *Actaea biternata* | 159761 |
| NC_077574 | *Actaea cimicifuga* | 159761 |
| NC_041533 | *Actaea dahurica* | 159370 |
| NC_042253 | *Actaea heracleifolia* | 159578 |
| NC_081475 | *Actaea japonica* | 159736 |
| NC_081476 | *Actaea purpurea* | 159497 |
| MN623225 | *Actaea simplex* | 159624 |
| NC_041543 | *Actaea vaginata* | 159110 |
| NC_056353 | *Adonis amurensis* | 157032 |
| MK253469 | *Adonis coerulea* | 157033 |
| NC_077573 | *Adonis mongolica* | 157521 |
| NC_065037 | *Adonis pseudoamurensis* | 156917 |
| NC_072342 | *Adonis ramosa* | 157057 |
| NC_041474 | *Adonis sutchuenensis* | 157603 |
| NC_037194 | *Anemoclema glaucifolium* | 160400 |
| NC_063528 | *Anemone alpina* | 162638 |
| NC_064389 | *Anemone altaica* | 160806 |
| NC_082177 | *Anemone coronaria* | 159564 |
| NC_060982 | *Anemone flaccida* | 157213 |
| NC_039465 | *Hepatica henryi* | 159276 |
| NC_045910 | *Hepatica asiatica* | 160157 |

| | | |
|---|---|---|
| NC_045878 | *Hepatica nobilis var. japonica* | 160988 |
| NC_066445 | *Anemone hortensis* | 158962 |
| NC_045909 | *Hepatica maxima* | 160876 |
| NC_045879 | *Anemone narcissiflora* | 158557 |
| NC_066446 | *Anemone nemorosa* | 160895 |
| NC_063529 | *Anemone occidentalis* | 161678 |
| MN025345 | *Anemone patens subsp. multifida* | 161936 |
| MN025344 | *Pulsatilla grandis* | 161376 |
| NC_041526 | *Anemone raddeana* | 160493 |
| NC_054280 | *Anemone reflexa* | 160910 |
| OR251125 | *Anemone rivularis var. flore-minore* | 162991 |
| NC_069812 | *Anemone shikokiana* | 159286 |
| NC_050873 | *Anemone taipaiensis* | 162084 |
| OR251126 | *Anemone thomsonii var. thomsonii* | 160058 |
| NC_039451 | *Anemone tomentosa* | 160945 |
| NC_039456 | *Anemone trullifolia* | 157096 |
| MK860686 | *Anemone turczaninovii* | 162795 |
| NC_056381 | *Anemone vitifolia* | 162203 |
| NC_041527 | *Anemonopsis macrophylla* | 158787 |
| NC_053820 | *Aquilegia barnebyi* | 161954 |
| NC_041528 | *Aquilegia coerulea* | 161429 |
| NC_041529 | *Aquilegia ecalcarata* | 161795 |
| NC_058528 | *Aquilegia kansuensis* | 162090 |
| NC_046738 | *Aquilegia rockii* | 162123 |
| NC_058527 | *Aquilegia yabeana* | 162094 |
| NC_058529 | *Aquilegia yangii* | 161820 |
| NC_041530 | *Asteropyrum cavaleriei* | 163660 |
| NC_045850 | *Asteropyrum peltatum* | 164455 |
| NC_041531 | *Beesia calthifolia* | 158355 |
| NC_072729 | *Beesia deltophylla* | 157506 |
| OY747139 | *Berberis vulgaris* | 166264 |
| NC_041475 | *Calathodes oxycarpa* | 160415 |
| MK253466 | *Callianthemum alatavicum* | 156538 |
| NC_041476 | *Callianthemum taipaicum* | 156939 |
| NC_041532 | *Caltha palustris* | 156167 |
| MK253464 | *Ceratocephala falcata* | 150821 |
| PP155435 | *Ceratocephala orthoceras* | 151407 |
| NC_069854 | *Ceratocephala testiculata* | 150820 |
| NC_046829 | *Chelidonium majus* | 159734 |
| NC_039844 | *Clematis acerifolia* | 159552 |
| NC_065272 | *Clematis acerifolia var. elobata* | 159690 |
| MT876505 | *Clematis alpina subsp. ochotensis* | 159631 |
| MT876514 | *Clematis alpina subsp. sibirica* | 159675 |
| NC_039577 | *Clematis alternata* | 159476 |
| NC_081049 | *Clematis apiifolia* | 159682 |
| NC_069848 | *Clematis armandii* | 159353 |
| NC_069849 | *Clematis aureolata* | 159548 |
| NC_065250 | *Clematis brachiata* | 159678 |
| NC_042793 | *Clematis brachyura* | 159532 |
| NC_039579 | *Clematis brevicaudata* | 159583 |
| NC_065251 | *Clematis cadmia* | 159714 |
| NC_066977 | *Clematis calcicola* | 159658 |
| NC_065264 | *Clematis campestris* | 159671 |
| NC_065265 | *Clematis canescens* | 159752 |
| NC_066718 | *Clematis chinensis* | 159497 |
| NC_065266 | *Clematis chrysocoma* | 159584 |
| NC_065267 | *Clematis connata* | 159768 |
| NC_065268 | *Clematis crassifolia* | 159312 |
| NC_065269 | *Clematis crispa* | 159304 |
| NC_065270 | *Clematis delavayi* | 159659 |
| NC_065271 | *Clematis drummondii* | 159729 |
| NC_058885 | *Clematis florida* | 159606 |
| NC_065273 | *Clematis fruticosa* | 159796 |
| NC_060524 | *Clematis fusca* | 159624 |
| NC_065274 | *Clematis glaucophylla* | 159302 |
| NC_081050 | *Clematis gouriana* | 159681 |
| NC_081051 | *Clematis grandidentata* | 159634 |
| NC_081052 | *Clematis gratopsis* | 159643 |
| NC_050373 | *Clematis guniuensis* | 159682 |
| NC_065275 | *Clematis haenkeana* | 156007 |
| NC_070190 | *Clematis henryi* | 159707 |
| NC_039845 | *Clematis heracleifolia* | 159565 |
| NC_065276 | *Clematis hexapetala* | 159534 |
| NC_065277 | *Clematis huchouensis* | 159590 |
| NC_081053 | *Clematis integrifolia* | 159630 |
| NC_065278 | *Clematis intricata* | 159593 |

| | | |
|---|---|---|
| NC_065279 | *Clematis lancifolia* | 159447 |
| NC_065286 | *Clematis lasiandra* | 159688 |
| NC_065281 | *Clematis leschenaultiana* | 159622 |
| NC_065282 | *Clematis ligusticifolia* | 159600 |
| NC_039690 | *Clematis loureiroana* | 159624 |
| NC_041477 | *Clematis macropetala* | 159647 |
| NC_079957 | *Clematis mandshurica* | 159564 |
| NC_057507 | *Clematis montana* | 159523 |
| NC_065283 | *Clematis nannophylla* | 159814 |
| NC_071961 | *Clematis orientalis* | 159543 |
| NC_069850 | *Clematis otophora* | 159534 |
| NC_081054 | *Clematis parviloba* | 159664 |
| NC_063559 | *Clematis patens* | 159603 |
| NC_085220 | *Clematis patens subsp. tientaiensis* | 159644 |
| NC_081055 | *Clematis peterae* | 159676 |
| NC_065284 | *Clematis petriei* | 159425 |
| NC_065248 | *Clematis pinnata* | 159652 |
| NC_058760 | *Clematis potaninii* | 159691 |
| NC_081056 | *Clematis psilandra* | 159622 |
| NC_065285 | *Clematis pubescens* | 159391 |
| NC_065287 | *Clematis quinquefoliolata* | 159724 |
| NC_081994 | *Clematis ranunculoides* | 159741 |
| NC_069851 | *Clematis rehderiana* | 159582 |
| NC_039578 | *Clematis repens* | 159507 |
| NC_065288 | *Clematis reticulata* | 159284 |
| NC_065289 | *Clematis rutoides* | 159678 |
| NC_060523 | *Clematis serratifolia* | 159648 |
| NC_065290 | *Clematis songorica* | 159822 |
| NC_081062 | *Clematis speciosa* | 159753 |
| NC_081057 | *Clematis stans* | 159671 |
| ON411443 | *Clematis stans var. austrojaponensis* | 159775 |
| NC_081058 | *Clematis subumbellata* | 159687 |
| NC_080984 | *Clematis tangutica* | 159584 |
| NC_028000 | *Clematis terniflora* | 159528 |
| NC_069852 | *Clematis tibetana* | 159540 |
| NC_065291 | *Clematis tomentella* | 159818 |
| NC_081059 | *Clematis tsugetorum* | 159611 |
| NC_065249 | *Clematis tubulosa* | 159653 |
| OP649578 | *Clematis tubulosa var. ichangensis* | 159645 |
| NC_039846 | *Clematis uncinata* | 159524 |
| NC_065292 | *Clematis urophylla* | 159696 |
| NC_081060 | *Clematis urticifolia* | 159720 |
| NC_065293 | *Clematis virginiana* | 159561 |
| NC_065294 | *Clematis viridis* | 159847 |
| NC_081061 | *Clematis vitalba* | 159710 |
| NC_065295 | *Clematis williamsii* | 159645 |
| NC_065296 | *Clematis xiangguiensis* | 159575 |
| NC_041534 | *Consolida ajacis* | 155900 |
| NC_047292 | *Consolida orientalis* | 155915 |
| NC_036485 | *Coptis chinensis* | 155484 |
| OM202495 | *Coptis chinensis var. brevisepala* | 154641 |
| MK569483 | *Coptis chinensis* | 154849 |
| NC_064406 | *Coptis deltoidea* | 154538 |
| NC_054329 | *Coptis japonica* | 154985 |
| NC_054330 | *Coptis omeiensis* | 154573 |
| NC_037759 | *Coptis quinquesecta* | 154549 |
| NC_054331 | *Coptis teeta* | 154156 |
| MK253461 | *Delphinium anthriscifolium* | 155077 |
| NC_051554 | *Delphinium brunonianum* | 153926 |
| MW246165 | *Delphinium candelabrum var. monanthum* | 153995 |
| MK253460 | *Delphinium ceratophorum* | 154245 |
| NC_049872 | *Delphinium grandiflorum* | 157339 |
| NC_047293 | *Delphinium maackianum* | 154484 |
| NC_056321 | *Delphinium yunnanense* | 154053 |
| MK253459 | *Dichocarpum dalzielii* | 153110 |
| NC_041478 | *Dichocarpum fargesii* | 153134 |
| MK253458 | *Dichocarpum sutchuenense* | 155390 |
| NC_041535 | *Enemion raddeanum* | 151916 |
| NC_053526 | *Epimedium parvifolium* | 157201 |
| NC_066652 | *Eranthis byunsanensis* | 160324 |
| NC_041536 | *Eranthis stellata* | 159251 |
| MK281585 | *Eschscholzia californica* | 160201 |
| PP155436 | *Ficaria verna* | 156439 |
| NC_041539 | *Glaucidium palmatum* | 156791 |
| NC_033341 | *Gymnaconitum gymnandrum* | 157327 |
| NC_062141 | *Helleborus atrorubens* | 166695 |

| | | |
|---|---|---|
| NC_041540 | *Helleborus thibetanus* | 155525 |
| NC_060983 | *Hepatica acutiloba* | 159497 |
| NC_060984 | *Hepatica americana* | 159805 |
| NC_060985 | *Hepatica falconeri* | 161075 |
| MG001340 | *Hepatica henryi* | 159276 |
| NC_062086 | *Hepatica insularis* | 160377 |
| MG952899 | *Hepatica maxima* | 160876 |
| NC_060986 | *Hepatica nobilis* | 160635 |
| MG952898 | *Hepatica nobilis var. japonica* | 160988 |
| NC_060987 | *Hepatica transsilvanica* | 161005 |
| NC_034702 | *Hydrastis canadensis* | 160000 |
| NC_041541 | *Isopyrum manshuricum* | 151243 |
| NC_041542 | *Leptopyrum fumarioides* | 157448 |
| NC_066184 | *Mahonia bodinieri* | 165697 |
| NC_012615 | *Megaleranthis saniculifolia* | 159924 |
| PP155437 | *Myosurus apetalus* | 150380 |
| PP155438 | *Myosurus minimus* | 150431 |
| NC_039542 | *Naravelia pilulifera* | 159513 |
| NC_039580 | *Naravelia zeylanica* | 159568 |
| NC_041537 | *Nigella damascena* | 155218 |
| NC_083219 | *Nigella sativa* | 154120 |
| NC_041538 | *Oxygraphis glacialis* | 155434 |
| NC_037831 | *Papaver rhoeas* | 152905 |
| NC_041479 | *Paraquilegia anemonoides* | 164383 |
| NC_067755 | *Paraquilegia microphylla* | 164390 |
| NC_061038 | *Pulsatilla campanella* | 162322 |
| NC_045908 | *Pulsatilla cernua var. koreana* | 162709 |
| NC_039452 | *Pulsatilla chinensis* | 162052 |
| MK860685 | *Pulsatilla dahurica* | 162450 |
| OP729488 | *Pulsatilla saxatilis* | 162659 |
| NC_061398 | *Pulsatilla tongkangensis* | 163442 |
| KX639503 | *Ranunculus austro-oreganus* | 154493 |
| MK253468 | *Ranunculus bungei* | 156082 |
| NC_045920 | *Ranunculus cantoniensis* | 155154 |
| NC_077490 | *Ranunculus cassubicifolius* | 156233 |
| NC_079824 | *Ranunculus chinensis* | 155289 |
| KY562595 | *Ranunculus flammula* | 156061 |
| MZ169045 | *Ranunculus japonicus* | 156981 |
| NC_008796 | *Ranunculus macranthus* | 155129 |
| NC_065303 | *Ranunculus membranaceus* | 156028 |
| NC_031651 | *Ranunculus occidentalis* | 154474 |
| NC_060613 | *Ranunculus pekinensis* | 156139 |
| NC_036976 | *Ranunculus repens* | 154247 |
| NC_036977 | *Ranunculus reptans* | 157239 |
| NC_080350 | *Ranunculus sceleratus* | 156329 |
| ON462450 | *Ranunculus silerifolius var. silerifolius* | 155368 |
| NC_081908 | *Ranunculus ternatus* | 156003 |
| NC_063514 | *Ranunculus yunnanensis* | 156050 |
| NC_039743 | *Semiaquilegia adoxoides* | 158340 |
| NC_057495 | *Semiaquilegia guangxiensis* | 164047 |
| NC_047294 | *Staphisagria macrosperma* | 155905 |
| NC_058830 | *Thalictrum aquilegiifolium* | 156253 |
| MW816628 | *Thalictrum aquilegiifolium var. sibiricum* | 156244 |
| MZ962406 | *Thalictrum baicalense* | 156196 |
| NC_061927 | *Thalictrum cirrhosum* | 155969 |
| NC_026103 | *Thalictrum coreanum* | 155088 |
| NC_085600 | *Thalictrum elegans* | 155864 |
| NC_070058 | *Thalictrum fargesii* | 155929 |
| NC_053570 | *Thalictrum foeniculaceum* | 155923 |
| NC_058920 | *Thalictrum foliolosum* | 155764 |
| OM501079 | *Thalictrum minus var. hypoleucum* | 156258 |
| NC_041544 | *Thalictrum minus* | 156217 |
| MK253449 | *Thalictrum petaloideum* | 155876 |
| NC_068627 | *Thalictrum simplex* | 156211 |
| MK253448 | *Thalictrum tenue* | 156103 |
| NC_039433 | *Thalictrum thalictroides* | 154924 |
| NC_058831 | *Thalictrum viscosum* | 155984 |
| NC_084427 | *Trollius altaicus* | 159986 |
| NC_084428 | *Trollius asiaticus* | 160002 |
| NC_084429 | *Trollius buddae* | 159688 |
| NC_031849 | *Trollius chinensis* | 160191 |
| NC_084430 | *Trollius dschungaricus* | 159816 |
| NC_050872 | *Trollius farreri* | 160611 |
| NC_084431 | *Trollius japonicus* | 160061 |
| NC_084432 | *Trollius ledebouri* | 160025 |
| NC_084433 | *Trollius lilacinus* | 160202 |

| | | |
|---|---|---|
| NC_059918 | *Trollius macropetalus* | 160094 |
| NC_084434 | *Trollius micranthus* | 159597 |
| NC_084435 | *Trollius pumilus* | 159598 |
| NC_084436 | *Trollius ranunculoides* | 159616 |
| NC_084437 | *Trollius taihasenzanensis* | 160019 |
| NC_084438 | *Trollius vaginatus* | 159723 |
| NC_084439 | *Trollius yunnanensis* | 159717 |
| NC_039744 | *Urophysa henryi* | 158303 |
| NC_039742 | *Urophysa rockii* | 158512 |

**(b)**

| **Genbank ID** | **Taxon** | **Sequence Length (bp)** |
|---|---|---|
| NC_084324 | *Aconitum carmichaelii* | 425319 |
| NC_053920 | *Aconitum kusnezoffii* | 440720 |
| NC_053368 | *Hepatica maxima* | 1122546 |
| OR100522 | *Corydalis pauciovulata* | 675483 |
| NC_072536 | *Paropyrum anemonoides* | 206722 |
| NC_068018 | *Pulsatilla cernua* | 747621 |
| NC_068017 | *Pulsatilla chinensis* | 878988 |
| OL979137 | *Pulsatilla chinensis var. kissii* | 684203 |
| NC_071219 | *Pulsatilla dahurica* | 824625 |
| NC_088759 | *Ranunculus cassubicifolius* | 1183731 |

**Table S2.** RNA-seq data (37) of Ranunculaceae from SRA/NCBI used for *R. cassubicifolius* genome annotation. See Materials and Methods for annotation details.

| SRA ID | Taxon |
|---|---|
| SRR21162053 | *Ranunculus carpaticola x Ranunculus notabilis* |
| SRR21162054 | *Ranunculus carpaticola x Ranunculus notabilis* |
| SRR21162055 | *Ranunculus notabilis* |
| SRR21162056 | *Ranunculus notabilis* |
| SRR21162057 | *Ranunculus carpaticola* |
| SRR11487683 | *Ranunculus caucasicus* |
| SRR11487684 | *Ranunculus acris* |
| SRR11487685 | *Ranunculus acris* |
| SRR13060786 | *Ranunculus japonicus* |
| SRR1737526 | *Ranunculus cantoniensis* |
| SRR1822558 | *Ranunculus brotherusii* |
| SRR3291759 | *Ranunculus sceleratus* |
| SRR958803 | *Ranunculus carpaticola x Ranunculus cassubicifolius* |
| SRR958819 | *Ranunculus carpaticola x Ranunculus cassubicifolius* |
| SRR958841 | *Ranunculus notabilis* |
| SRR958846 | *Ranunculus cassubicifolius* |
| SRR1822529 | *Ranunculus bungei* |
| SRR18512371 | *Ranunculus bungei* |
| SRR18512372 | *Ranunculus bungei* |
| SRR18512373 | *Ranunculus bungei* |
| SRR18512374 | *Ranunculus bungei* |
| SRR18512375 | *Ranunculus bungei* |
| SRR18512376 | *Ranunculus bungei* |
| SRR18512381 | *Ranunculus bungei* |
| SRR18512383 | *Ranunculus bungei* |
| SRR18512384 | *Ranunculus bungei* |
| SRR18512385 | *Ranunculus bungei* |
| SRR18512386 | *Ranunculus bungei* |
| SRR18512387 | *Ranunculus bungei* |
| SRR18512388 | *Ranunculus bungei* |
| SRR18512389 | *Ranunculus bungei* |
| SRR18512390 | *Ranunculus bungei* |
| SRR18512391 | *Ranunculus bungei* |
| SRR18512392 | *Ranunculus bungei* |
| SRR18512398 | *Ranunculus bungei* |
| SRR18512400 | *Ranunculus bungei* |
| SRR18512401 | *Ranunculus bungei* |

**Table S3.** Genome annotation data (14) of Ranunculales from NCBI used for genome annotation of *R. cassubicifolius*.

| Tool | Species | Family | Order | NCBI Accession | Reference |
|---|---|---|---|---|---|
| Funanotate and Braker | *Aquilegia coerulea* | Ranunculaceae | Ranunculales | GCA_002738505.1 | https://www.ncbi.nlm.nih.gov/datasets/genome/GCA_002738505.1/ |
| Funanotate and Braker | *Coptis chinensis* | Ranunculaceae | Ranunculales | GCA_015680905.1 | https://doi.org/10.1038/s41467-021-23611-0 |
| Funanotate and Braker | *Thalictrum thalictroides* | Ranunculaceae | Ranunculales | GCA_013358455.1 | 10.1002/aps3.11407 |
| Funanotate | *Kingdonia uniflora* | Circaeasteraceae | Ranunculales | GCA_014058105.1 | 10.1016/j.isci.2020.101124 |
| Funanotate | *Macleaya cordata* | Papaveraceae | Ranunculales | GCA_002174775.1 | https://doi.org/10.1016/j.molp.2017.05.007 |
| Funanotate | *Papaver armeniacum* | Papaveraceae | Ranunculales | GCA_023531295.1 | https://doi.org/10.1038/s41467-022-30856-w |
| Funanotate | *Papaver atlanticum* | Papaveraceae | Ranunculales | GCA_023531105.1 | https://doi.org/10.1038/s41467-022-30856-w |
| Funanotate | *Papaver bracteatum* | Papaveraceae | Ranunculales | GCA_023529315.1 | https://doi.org/10.1038/s41467-022-30856-w |
| Funanotate | *Papaver californicum* | Papaveraceae | Ranunculales | GCA_023531435.1 | https://doi.org/10.1038/s41467-022-30856-w |
| Funanotate | *Papaver nudicaule* | Papaveraceae | Ranunculales | GCA_023529015.1 | https://doi.org/10.1038/s41467-022-30856-w |
| Funanotate | *Papaver somniferum* | Papaveraceae | Ranunculales | GCF_003573695.1 | 10.1126/science.aat4096 |
| Funanotate | *Stephania cephalantha* | Menispermaceae | Ranunculales | GCA_039657325.1 | https://doi.org/10.1038/s41467-024-45690-5 |
| Funanotate | *Stephania japonica* | Menispermaceae | Ranunculales | GCA_039657345.1 | https://doi.org/10.1038/s41467-024-45690-5 |
| Funanotate | *Stephania yunnanensis* | Menispermaceae | Ranunculales | GCA_039657365.1 | https://doi.org/10.1038/s41467-024-45690-5 |

**Table S4.** QUAST statistics for different genome assembly strategies of the diploid sexual species *R. cassubicifolius* (LH040) (see also reports on Figshare). SUPERNOVA, and partially SPAdes, MaSuRCA, and WENGAN used Illumina short-reads (22x 'Illumina'). Canu, Flye, and Hifiasm, and partially SPAdes, MaSuRCA, and WENGAN used ONT or PacBio long-reads (16x 'ONT', 25x 'PacBio'). The polishing of the Canu, Flye, and Hifiasm assemblies includes the following steps: polishing via filtered ONT reads using Racon and Medaka / filtered PacBio reads using Racon and polishCLR, respectively, and via filtered Illumina reads using POLCA (MaSuRCA pipeline). Key parameters and the best assembly in terms of completeness and contig length are highlighted in bold. $N_{50}$ describes the shortest contig length required to summarize at least half (50%) of the bases of the entire assembly; $L_{50}$ is defined as the minimum number of contigs that encompass half (50%) of the bases of the entire assembly. See Table 1 for comparisons with downsampled (16x) PacBio reads. - = no results produced.

| **Metric** | **SUPERNOVA** (Illumina22x) | **SPAdes** (Illumina22x) | **SPAdes** (Illumina22x + ONT16x /PacBio25x) | **MaSuRCA** (Illumina22x + ONT16x /PacBio25x) | **WENGAN** (Illumina22x + ONT/ PacBio25x) | **Canu** (ONT16x /PacBio25x) | **Canu** (ONT16x /PacBio25x) +3x Polish | **Flye** (ONT16x /PacBio25x) | **Flye** (ONT16x /PacBio25x) +3x Polish | **Hifiasm** (PacBio25x/ PacBio16x) | **Hifiasm** (PacBio25x) +3x Polish |
|---|---|---|---|---|---|---|---|---|---|---|---|
| **Total assembly length, Mbp** | 2,76 (86%) | 0,557 (17%) | 0.408/ (13%)/ 0.464 (15%) | 0.076/ (2.4%)/ - | 0.005 (0.16%)/ 1.21 (38%) | 0.80 (25%)/ 5.12 (160%) | 0.81 (25%)/ 5.12 (160%) | 3.58 (112%)/ 4.93 (154%) | 3.45 (108%)/ - | 3.22 (101%)/ 3.62 (113%) | **3.21 (100%)** |
| **No. contigs** | 6,920,808 | 1,231,317 | 1,184,036/ 555,617 | 38,650/ - | 368/ 6,565,547 | 30,612/ 5,840 | 32,959/ 5,208 | 65,232/ 8,104 | 54,310/ - | 1,453/ 3449 | **1319** |
| **No. contigs > 1 Mbp** | 562,716 | 122,407 | 110,594/ 103,406 | 12,799/ - | 368/ 5,056 | 30,612/ 5,840 | 31,072/ 5,208 | 64,618/ 8,094 | 53,572/ - | 1,453/ 3,449 | **1319** |
| **Largest contig, Mbp** | 0.039 | 0.069 | 0.096/ 0.111 | 0.145/ - | 0.095/ 0.009 | 0.540/ 5.10 | 0.542/ 51.6 | 0.976/ 12.6 | 0.974/ - | 84/ 32 | **84** |
| **GC, %** | 40 | 39 | 39/ 39 | 40/ - | 40 / 39 | 40/ 42 | 40/ 42 | 43/ 42 | 43/ - | 42/ 42 | **42** |
| **$N_{50}$, Mbp** | 0.002 | 0.003 | 0.005/ 0.005 | 0.010/ - | 0.021/ 0.006 | 0.037/ 3.18 | 0.038/ 3.19 | 0.103/ 1.73 | 0.104 -/ | 20.0/ 2.89 | **20.0** |
| **$L_{50}$** | 259,069 | 29,900 | 23,910/ 22,261 | 1,507/ - | 76/ 28,344 | 6,935/ 469 | 6,937/ 467 | 10,305/ 842 | 9,889 -/ | 45/ 350 | **45** |

**Table S5.** BUSCO statistics of nuclear genome sequence assemblies. (a) Short-read (Illumina), (b) hybrid-read (Illumina + ONT/PacBio), and (c) long-read (ONT/PacbBio) assemblies. Total BUSCO groups searched = 1614. Complete and single-copy BUSCOs (S), complete and duplicated BUSCOs (D), fragmented BUSCOs (F), and missing BUSCOs (M). See Figure S7 and further details in the Materials and Methods sections.

**(a)**

| | Assemblies | | | |
|---|---|---|---|---|
| **BUSCO scores** | **SUPERNOVA_ Illumina22x** | **SUPERNOVA_ Illumina22x_perc** | **SPAdes_ Illumina22x** | **SPAdes_ Illumina22x_perc** |
| S | 317 | 19.64 | 969 | 60.04 |
| D | 178 | 11.03 | 32 | 1.98 |
| F | 576 | 35.69 | 444 | 27.51 |
| M | 543 | 33.64 | 169 | 10.47 |

**(b)**

| | **Assemblies** | | | | | |
|---|---|---|---|---|---|---|
| **BUSCO scores** | **SPAdes Illumina22x ONT16x** | **SPAdes Illumina22x ONT16x_perc** | **MaSuRCA Illumina22x ONT16x** | **MaSuRCA Illumina22x ONT16x_perc** | **WENGAN Illumina22x ONT16x** | **WENGAN Illumina22x ONT16x _perc** |
| S | 1059 | 65.61 | 87 | 5.39 | 3 | 0.19 |
| D | 38 | 2.35 | 2 | 0.12 | 0 | 0.00 |
| F | 393 | 24.35 | 109 | 6.75 | 4 | 0.25 |
| M | 124 | 7.68 | 1416 | 87.73 | 1607 | 99.57 |

| | **SPAdes Illumina22x PacBio16x** | **SPAdes Illumina22x PacBio16x_perc** | **MaSuRCA Illumina22x PacBio16x** | **MaSuRCA Illumina22x PacBio16x_perc** | **WENGAN Illumina22x PacBio16x** | **WENGAN Illumina22x PacBio16x_perc** |
|---|---|---|---|---|---|---|
| S | 1082 | 67.04 | - | - | 16 | 0.99 |
| D | 39 | 2.42 | - | - | 0 | 0.00 |
| F | 372 | 23.05 | - | - | 145 | 8.98 |
| M | 121 | 7.50 | - | - | 1453 | 90.02 |

| | **SPAdes Illumina22x PacBio25x** | **SPAdes Illumina22x PacBio25x_perc** | **MaSuRCA Illumina22x PacBio25x** | **MaSuRCA Illumina22x PacBio25x_perc** | **WENGAN Illumina22x PacBio25x** | **WENGAN Illumina22x PacBio25x_perc** |
|---|---|---|---|---|---|---|
| S | 1108 | 68.65 | - | - | 16 | 0.99 |
| D | 34 | 2.11 | - | - | 0 | 0.00 |
| F | 341 | 21.13 | - | - | 141 | 8.74 |
| M | 131 | 8.12 | - | - | 1457 | 90.27 |

**(c)**

| | Assemblies | | | | | | | |
|---|---|---|---|---|---|---|---|---|
| **BUSCO scores** | **Canu ONT16x** | **Canu ONT16x perc** | **Canu ONT16x Polished** | **Canu ONT16x Polished _perc** | **Flye ONT16x** | **Flye ONT16x perc** | **Flye ONT16x Polished** | **Flye ONT16x Polished _per** |
| S | 907 | 56.20 | 1068 | 66.17 | 959 | 59.42 | 1093 | 67.72 |
| D | 127 | 7.87 | 168 | 10.41 | 225 | 13.94 | 475 | 29.43 |
| F | 114 | 7.06 | 54 | 3.35 | 102 | 6.32 | 35 | 2.17 |
| M | 466 | 28.87 | 324 | 20.07 | 328 | 20.32 | 11 | 0.68 |

| | **Canu PacBio16x** | **Canu PacBio16x perc** | **Canu PacBio16x Polished** | **Canu PacBio16x Polished perc** | **Flye PacBio16x** | **Flye PacBio16x perc** | **Flye PacBio16x Polished** | **Flye PacBio16x Polished perc** | **Hifiasm Pacbio16x** | **Hifiasm PacBio16x perc** |
|---|---|---|---|---|---|---|---|---|---|---|
| S | 488 | 30.24 | 403 | 24.97 | 909 | 56.32 | 167 | 10.35 | 909 | 56.32 |
| D | 1113 | 68.96 | 1199 | 74.29 | 694 | 43.00 | 1435 | 88.91 | 694 | 43.00 |
| F | 7 | 0.43 | 7 | 0.43 | 7 | 0.43 | 7 | 0.43 | 7 | 0.43 |
| M | 6 | 0.37 | 5 | 0.31 | 4 | 0.25 | 5 | 0.31 | 4 | 0.25 |

| | **Canu PacBio25x** | **Canu PacBio25x perc** | **Canu PacBio25x Polished** | **Canu PacBio25x Polished perc** | **Flye PacBio25x** | **Flye PacBio25x perc** | **Flye PacBio25x Polished*** | **Flye PacBio25x Polished Perc*** | **Hifiasm Pacbio25x** | **Hifiasm PacBio25x perc** | **Hifiasm PacBio25x Polished** | **Hifiasm PacBio25x Polished _perc** |
|---|---|---|---|---|---|---|---|---|---|---|---|---|
| S | 468 | 29.00 | 354 | 21.93 | 464 | 28.75 | | | 1232 | 76.33 | 1217 | 75.40 |
| D | 1138 | 70.51 | 1252 | 77.57 | 1141 | 70.69 | | | 368 | 22.80 | 386 | 23.92 |
| F | 5 | 0.31 | 5 | 0.31 | 6 | 0.37 | | | 8 | 0.50 | 4 | 0.25 |
| M | 3 | 0.19 | 3 | 0.19 | 3 | 0.19 | | | 6 | 0.37 | 7 | 0.43 |

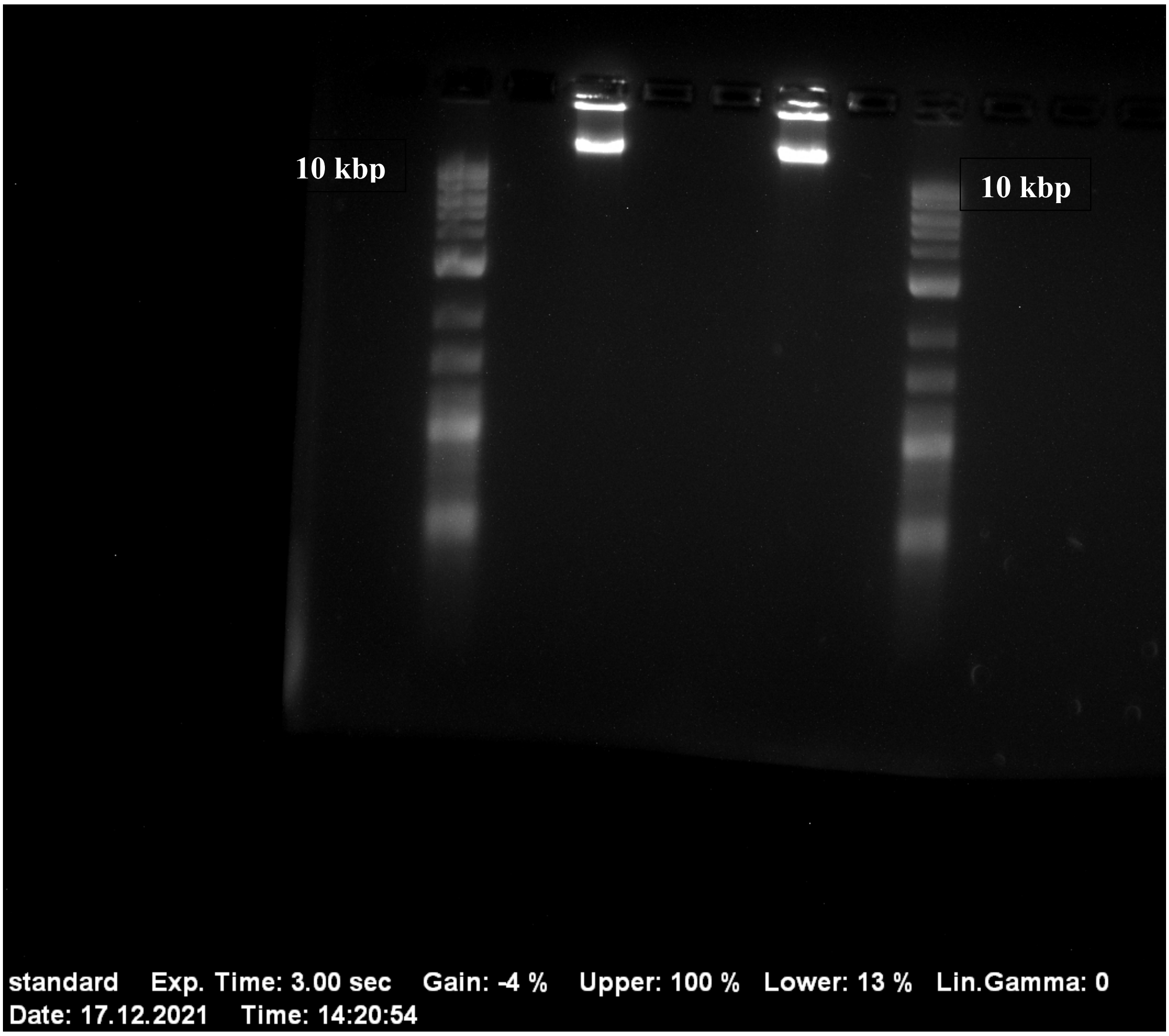


**Figure S1.** Gel electrophoresis of gDNA extractions from 17th December 2021 using 1 kb DNA Ladder (New England Biolabs Inc., Ipswich, USA; 500 bp - 10 kb) as size standard. The gDNA extract was applied in the central lanes, surrounded on the left and right by the DNA ladder. The top DNA fragment of the ladder is 10 kb. The extracted gDNA is substantially larger than 10 kb, and some very large fragments were not even able to move in the gel. The ONT analyses showed a median fragment length distribution of $N_{50}$ = ca. 23 kb, ranging up to 236 kb.

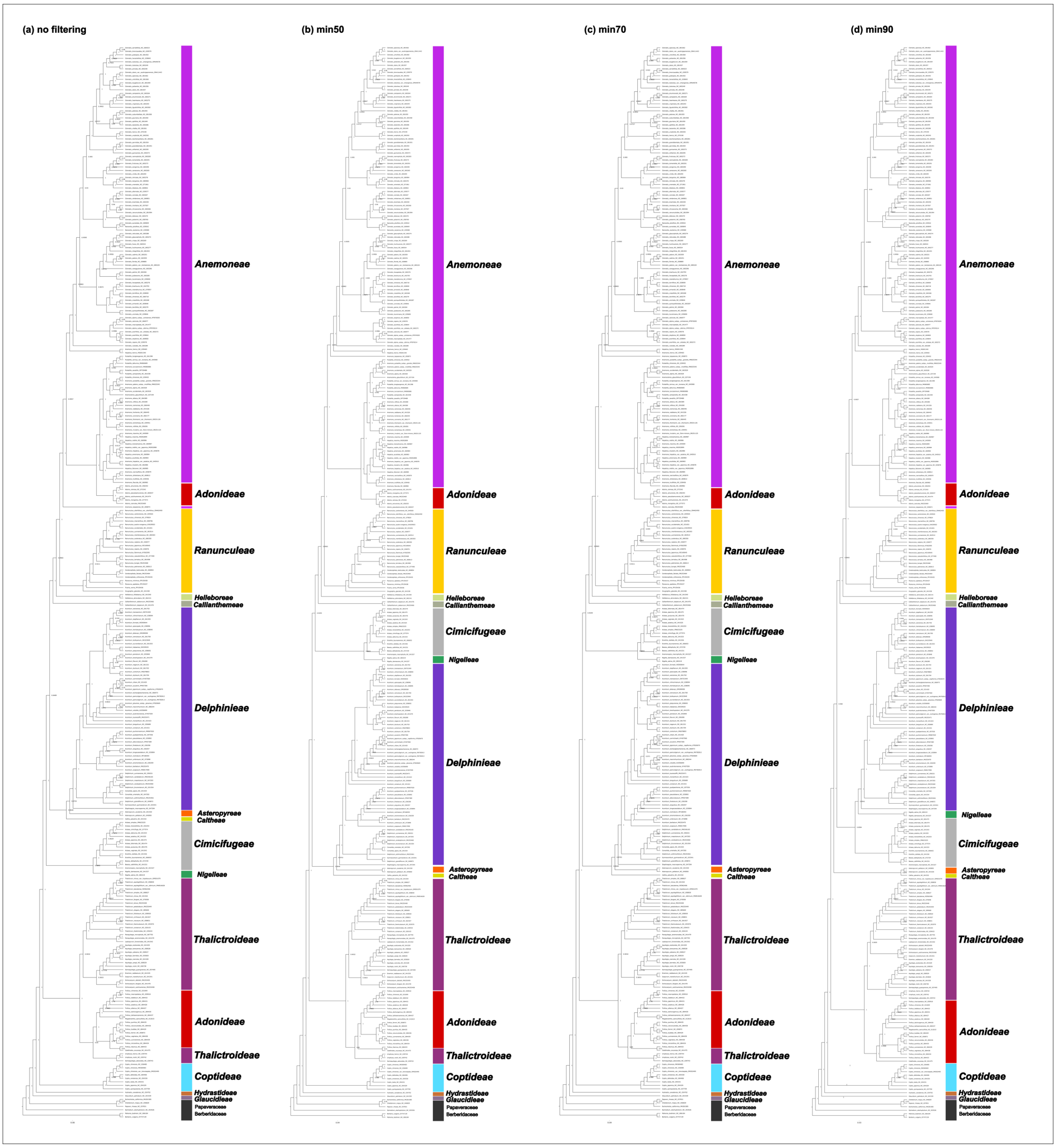


**Figure S2.** (a-d) Maximum-likelihood phylogeny based on min0 (no filtering), min50, min70, and min90 alignments of 306 plastomes (taxa) of the plant family Ranunculaceae. Transfer expectation values (TBE) are shown. See the Materials & Methods section for more details. The file is attached to the Supplements.

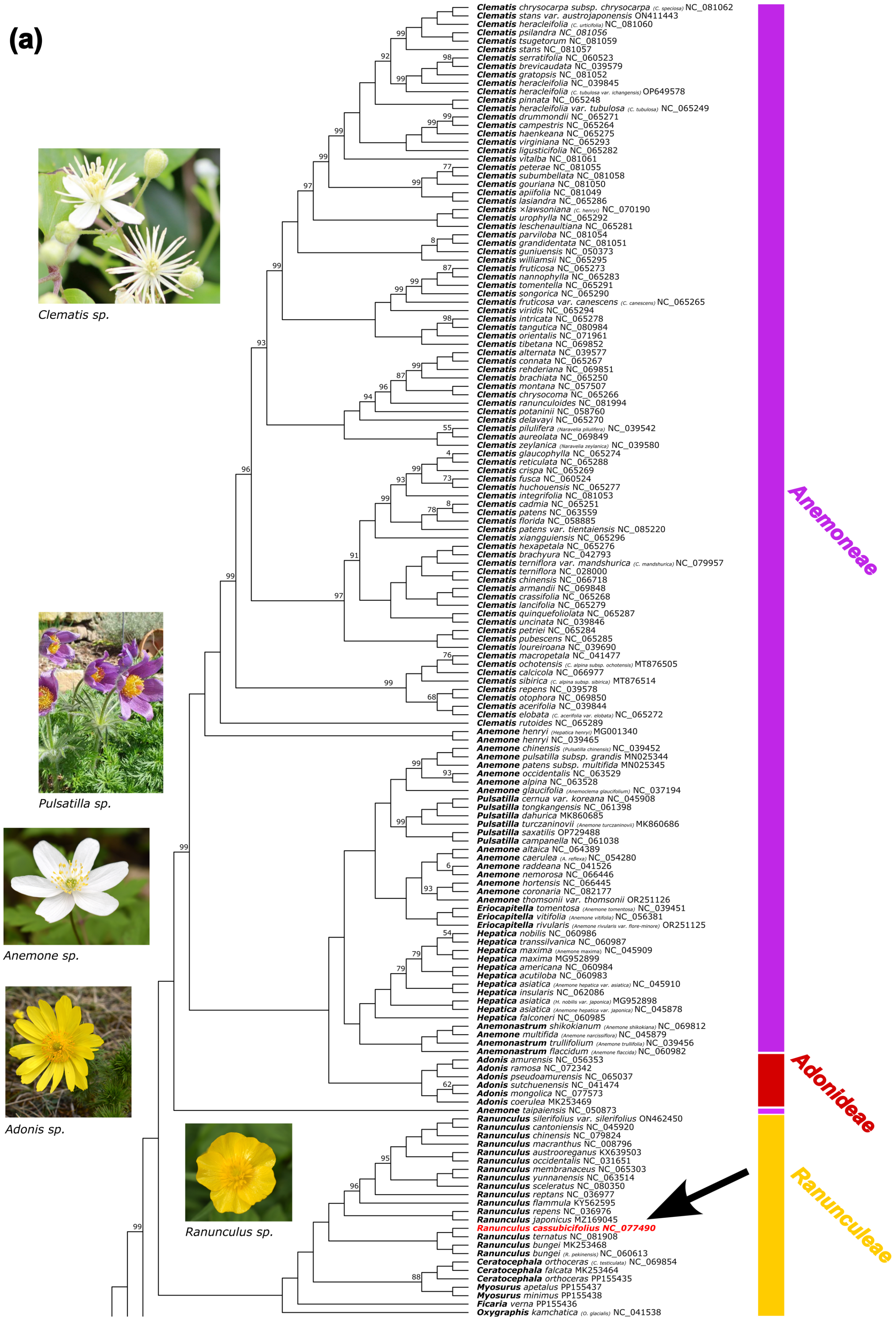
(a)
Clematis sp.
Pulsatilla sp.
Anemone sp.
Adonis sp.
Ranunculus sp.
Clematis chrysocarpa subsp. chrysocarpa (C. speciosa) NC_081062
Clematis stans var. austrojaponensis ON411443
Clematis heracleifolia (C. urticifolia) NC_081060
Clematis psilandra NC_081056
Clematis tsugetorum NC_081059
Clematis stans NC_081057
Clematis serratifolia NC_060523
Clematis brevicaudata NC_039579
Clematis gratopsis NC_081052
Clematis heracleifolia NC_039845
Clematis heracleifolia (C. tubulosa var. ichangensis) OP649578
Clematis pinnata NC_065248
Clematis heracleifolia var. tubulosa (C. tubulosa) NC_065249
Clematis drummondii NC_065271
Clematis campestris NC_065264
Clematis haenkeana NC_065275
Clematis virginiana NC_065293
Clematis ligusticifolia NC_065282
Clematis vitalba NC_081061
Clematis peterae NC_081055
Clematis subumbellata NC_081058
Clematis gouriana NC_081050
Clematis apiifolia NC_081049
Clematis lasiandra NC_065286
Clematis ×lawsoniana (C. henryi) NC_070190
Clematis urophylla NC_065292
Clematis leschenaultiana NC_065281
Clematis parviloba NC_081054
Clematis grandidentata NC_081051
Clematis guniuensis NC_050373
Clematis williamsii NC_065295
Clematis fruticosa NC_065273
Clematis nannophylla NC_065283
Clematis tomentella NC_065291
Clematis songorica NC_065290
Clematis fruticosa var. canescens (C. canescens) NC_065265
Clematis viridis NC_065294
Clematis intricata NC_065278
Clematis tangutica NC_080984
Clematis orientalis NC_071961
Clematis tibetana NC_069852
Clematis alternata NC_039577
Clematis connata NC_065267
Clematis rehderiana NC_069851
Clematis brachiata NC_065250
Clematis montana NC_057507
Clematis chrysocoma NC_065266
Clematis ranunculoides NC_081994
Clematis potaninii NC_058760
Clematis delavayi NC_065270
Clematis pilulifera (Naravelia pilulifera) NC_039542
Clematis aureolata NC_069849
Clematis zeylanica (Naravelia zeylanica) NC_039580
Clematis glaucophylla NC_065274
Clematis reticulata NC_065288
Clematis crispa NC_065269
Clematis fusca NC_060524
Clematis huchouensis NC_065277
Clematis integrifolia NC_081053
Clematis cadmia NC_065251
Clematis patens NC_063559
Clematis florida NC_058885
Clematis patens var. tientaiensis NC_085220
Clematis xiangguiensis NC_065296
Clematis hexapetala NC_065276
Clematis brachyura NC_042793
Clematis terniflora var. mandshurica (C. mandshurica) NC_079957
Clematis terniflora NC_028000
Clematis chinensis NC_066718
Clematis armandii NC_069848
Clematis crassifolia NC_065268
Clematis lancifolia NC_065279
Clematis quinquefoliolata NC_065287
Clematis uncinata NC_039846
Clematis petriei NC_065284
Clematis pubescens NC_065285
Clematis loureiroana NC_039690
Clematis macropetala NC_041477
Clematis ochotensis (C. alpina subsp. ochotensis) MT876505
Clematis calcicola NC_066977
Clematis sibirica (C. alpina subsp. sibirica) MT876514
Clematis repens NC_039578
Clematis otophora NC_069850
Clematis acerifolia NC_039844
Clematis elobata (C. acerifolia var. elobata) NC_065272
Clematis rutoides NC_065289
Anemone henryi (Hepatica henryi) MG001340
Anemone henryi NC_039465
Anemone chinensis (Pulsatilla chinensis) NC_039452
Anemone pulsatilla subsp. grandis MN025344
Anemone patens subsp. multifida MN025345
Anemone occidentalis NC_063529
Anemone alpina NC_063528
Anemone glaucifolia (Anemoclema glaucifolium) NC_037194
Pulsatilla cernua var. koreana NC_045908
Pulsatilla tongkangensis NC_061398
Pulsatilla dahurica MK860685
Pulsatilla turczaninovii (Anemone turczaninovii) MK860686
Pulsatilla saxatilis OP729488
Pulsatilla campanella NC_061038
Anemone altaica NC_064389
Anemone caerulea (A. reflexa) NC_054280
Anemone raddeana NC_041526
Anemone nemorosa NC_066446
Anemone hortensis NC_066445
Anemone coronaria NC_082177
Anemone thomsonii var. thomsonii OR251126
Eriocapitella tomentosa (Anemone tomentosa) NC_039451
Eriocapitella vitifolia (Anemone vitifolia) NC_056381
Eriocapitella rivularis (Anemone rivularis var. flore-minore) OR251125
Hepatica nobilis NC_060986
Hepatica transsilvanica NC_060987
Hepatica maxima (Anemone maxima) NC_045909
Hepatica maxima MG952899
Hepatica americana NC_060984
Hepatica acutiloba NC_060983
Hepatica asiatica (Anemone hepatica var. asiatica) NC_045910
Hepatica insularis NC_062086
Hepatica asiatica (H. nobilis var. japonica) MG952898
Hepatica asiatica (Anemone hepatica var. japonica) NC_045878
Hepatica falconeri NC_060985
Anemonastrum shikokianum (Anemone shikokiana) NC_069812
Anemone multifida (Anemone narcissiflora) NC_045879
Anemonastrum trullifolium (Anemone trullifolia) NC_039456
Anemonastrum flaccidum (Anemone flaccida) NC_060982
Adonis amurensis NC_056353
Adonis ramosa NC_072342
Adonis pseudoamurensis NC_065037
Adonis sutchuenensis NC_041474
Adonis mongolica NC_077573
Adonis coerulea MK253469
Anemone taipaiensis NC_050873
Ranunculus silerifolius var. silerifolius ON462450
Ranunculus cantoniensis NC_045920
Ranunculus chinensis NC_079824
Ranunculus macranthus NC_008796
Ranunculus austrooreganus KX639503
Ranunculus occidentalis NC_031651
Ranunculus membranaceus NC_065303
Ranunculus yunnanensis NC_063514
Ranunculus sceleratus NC_080350
Ranunculus reptans NC_036977
Ranunculus flammula KY562595
Ranunculus repens NC_036976
Ranunculus japonicus MZ169045
Ranunculus cassubicifolius NC_077490
Ranunculus ternatus NC_081908
Ranunculus bungei MK253468
Ranunculus bungei (R. pekinensis) NC_060613
Ceratocephala orthoceras (C. testiculata) NC_069854
Ceratocephala falcata MK253464
Ceratocephala orthoceras PP155435
Myosurus apetalus PP155437
Myosurus minimus PP155438
Ficaria verna PP155436
Oxygraphis kamchatica (O. glacialis) NC_041538
Anemoneae
Adonideae
Ranunculeae

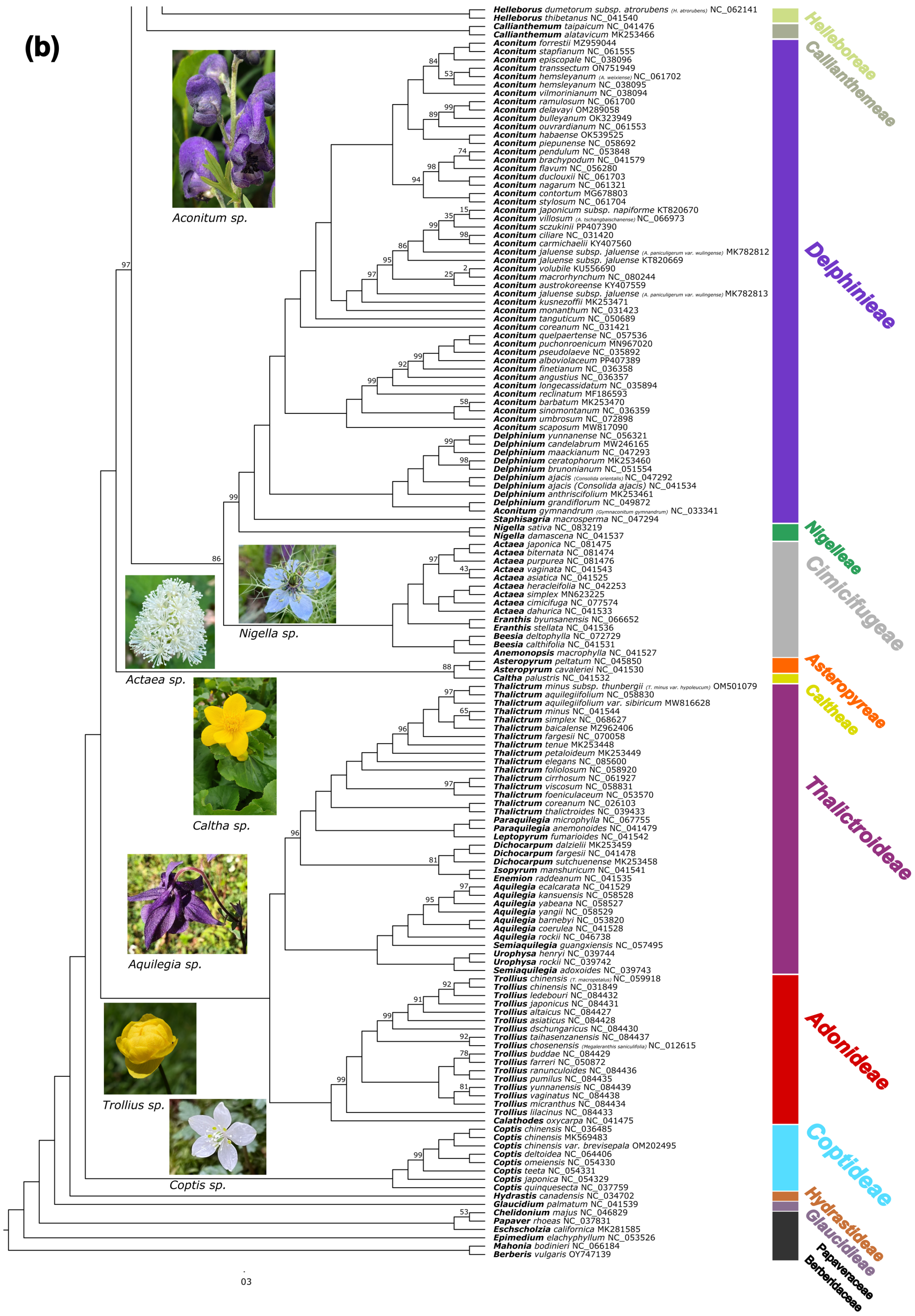


**Figure S3.** (a, b) Maximum-likelihood phylogeny based on 306 plastomes (taxa) of the plant family Ranunculaceae. All branches received full (1) transfer bootstrap expectation (TBE)

values, unless otherwise shown. Quartet Sampling (QS)  metrics are found in Fig. S4. All NCBI species names were checked by gbif.org, and synonyms were replaced by accepted names (synonyms in brackets). The color of clades corresponds to tribe/subfamily names (tribal classification follows Wang et al. (2016) and Zhai et al. (2019), and the plastome of *R. cassubicifolius* assembled in this study is highlighted with a black arrow. Modified images of *Anemone sp.*, *Pulsatilla sp.*, *Adonis sp.*, *Nigella sp.*, *Caltha sp.*, *Aquilegia sp.*, and *Trollius sp.* by Kevin Karbstein and from inaturalist.org (CC-BY-NC), of *Clematis sp.* (photos/344038634), *Aconitum sp.* (observations/175339012), *Actaea sp.* (observations/154486471), and *Coptis sp.* (observations/152014320). See Texts S2 and S3 for more details. The file is attached to the Supplements.

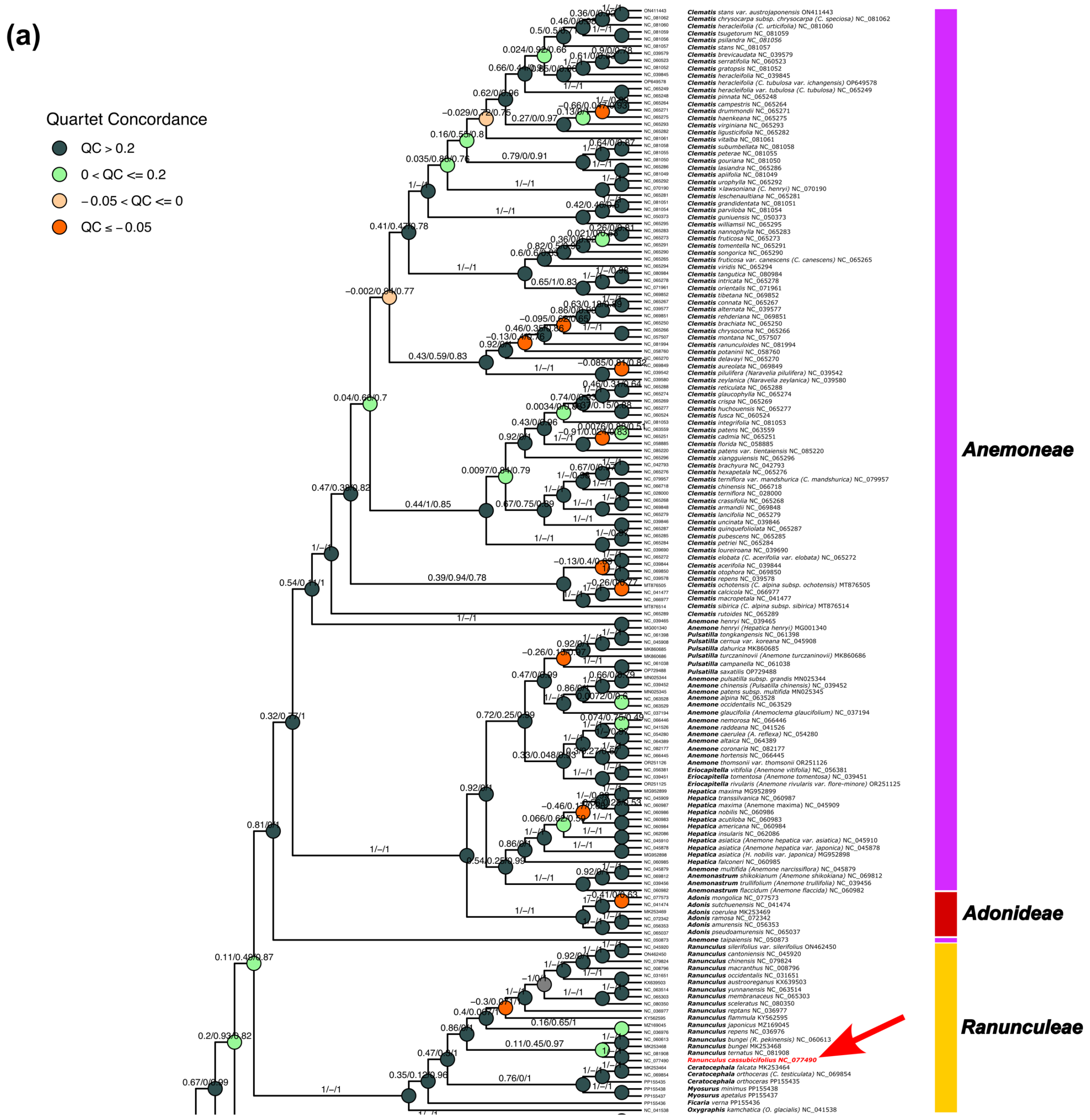

(a)
Quartet Concordance
QC > 0.2
0 < QC <= 0.2
−0.05 < QC <= 0
QC ≤ −0.05
Anemoneae
Adonideae
Ranunculeae
Ranunculus cassubicifolius NC_077490

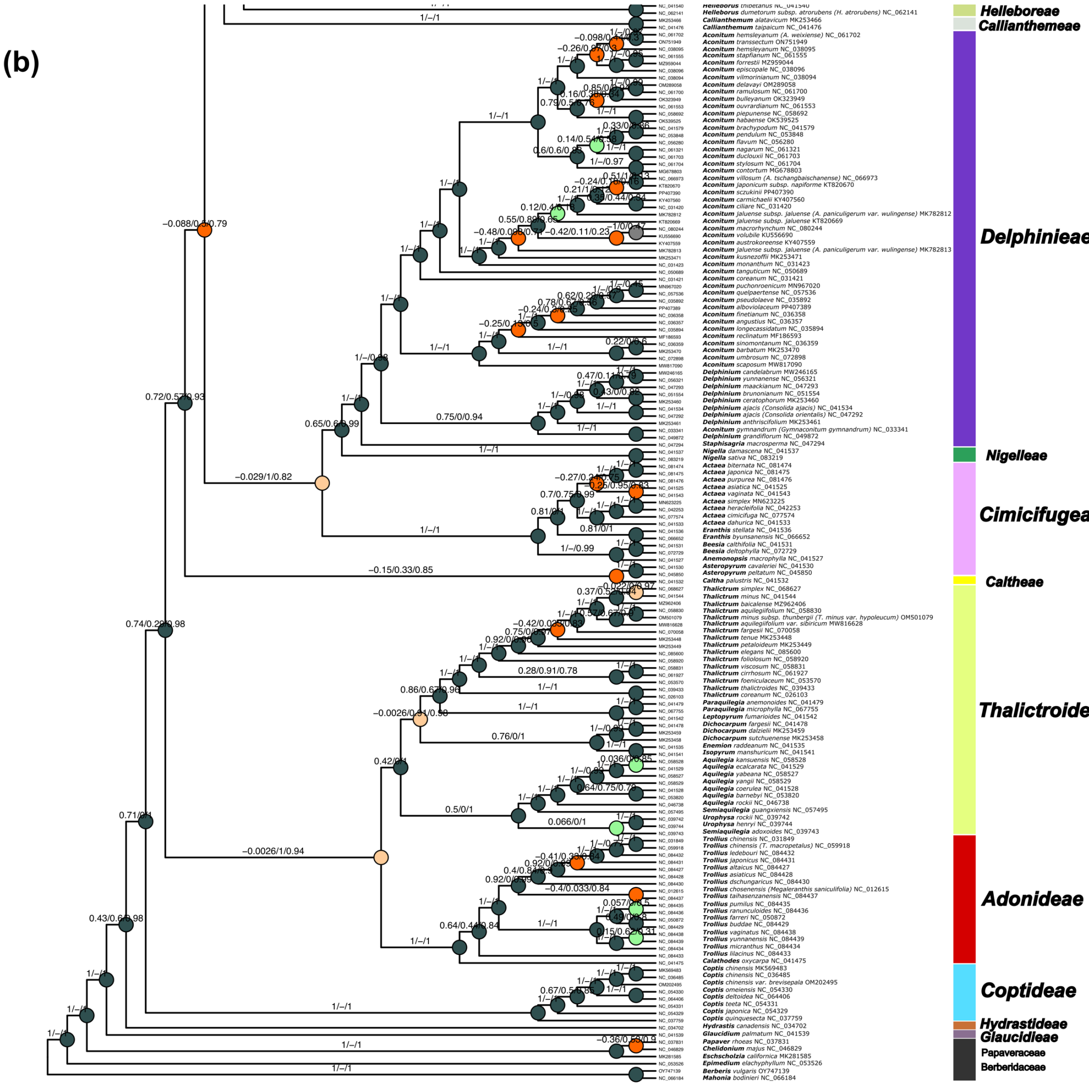


**Figure S4**. (a, b) Maximum-likelihood phylogeny based on 306 plastomes (taxa) and the min90 alignment of the plant family Ranunculaceae. See Pease et al. (2018) and Karbstein et al. (2020) for interpretation of QS metrics, and the Materials and Methods section for more details. The file is attached to the Supplements.

**(a)**

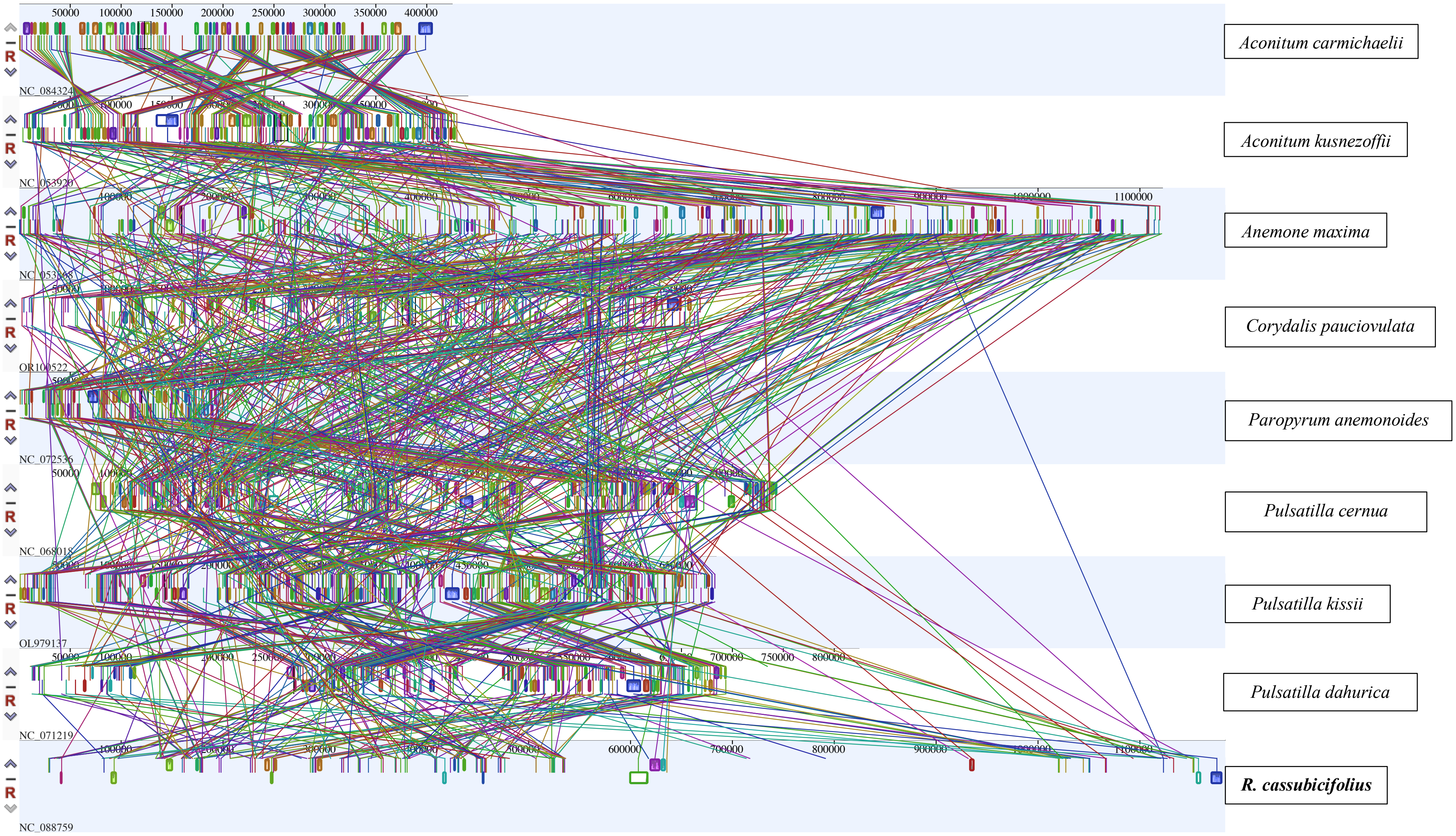
NC_084324
Aconitum carmichaelii
NC_053920
Aconitum kusnezoffii
Anemone maxima
OR100522
Corydalis pauciovulata
NC_072536
Paropyrum anemonoides
NC_068018
Pulsatilla cernua
OL979137
Pulsatilla kissii
NC_071219
Pulsatilla dahurica
NC_088759
R. cassubicifolius

**(b)**

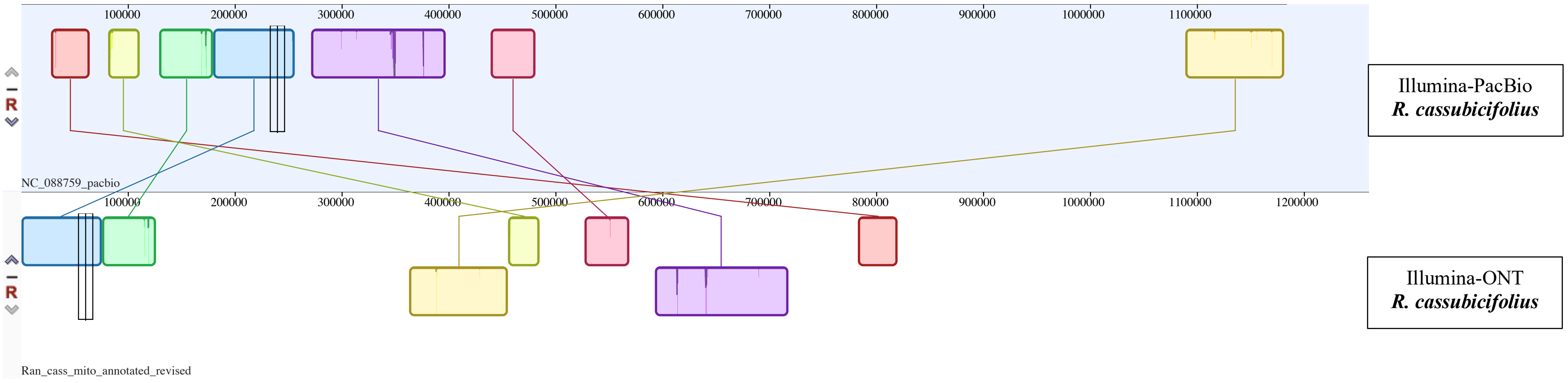


**Figure S5.** Whole genome alignment analysis of (a) all available mitogenome sequences in Ranunculaceae, and (b) of the assembled Illumina-ONT and -PacBio genome sequences of *R. cassubicifolius* (LH040). We performed a progressiveMAUVE analysis (2015-02-25; Darling et al., 2010). Shown are locally collinear blocks, i.e., homologous regions of sequences shared by the genomes under study, without any rearrangements of homologous sequences. The figures show substantial rearrangements of collinear blocks (a) among species but also (b) between individuals of the same species. See the Materials and Methods section for more details.

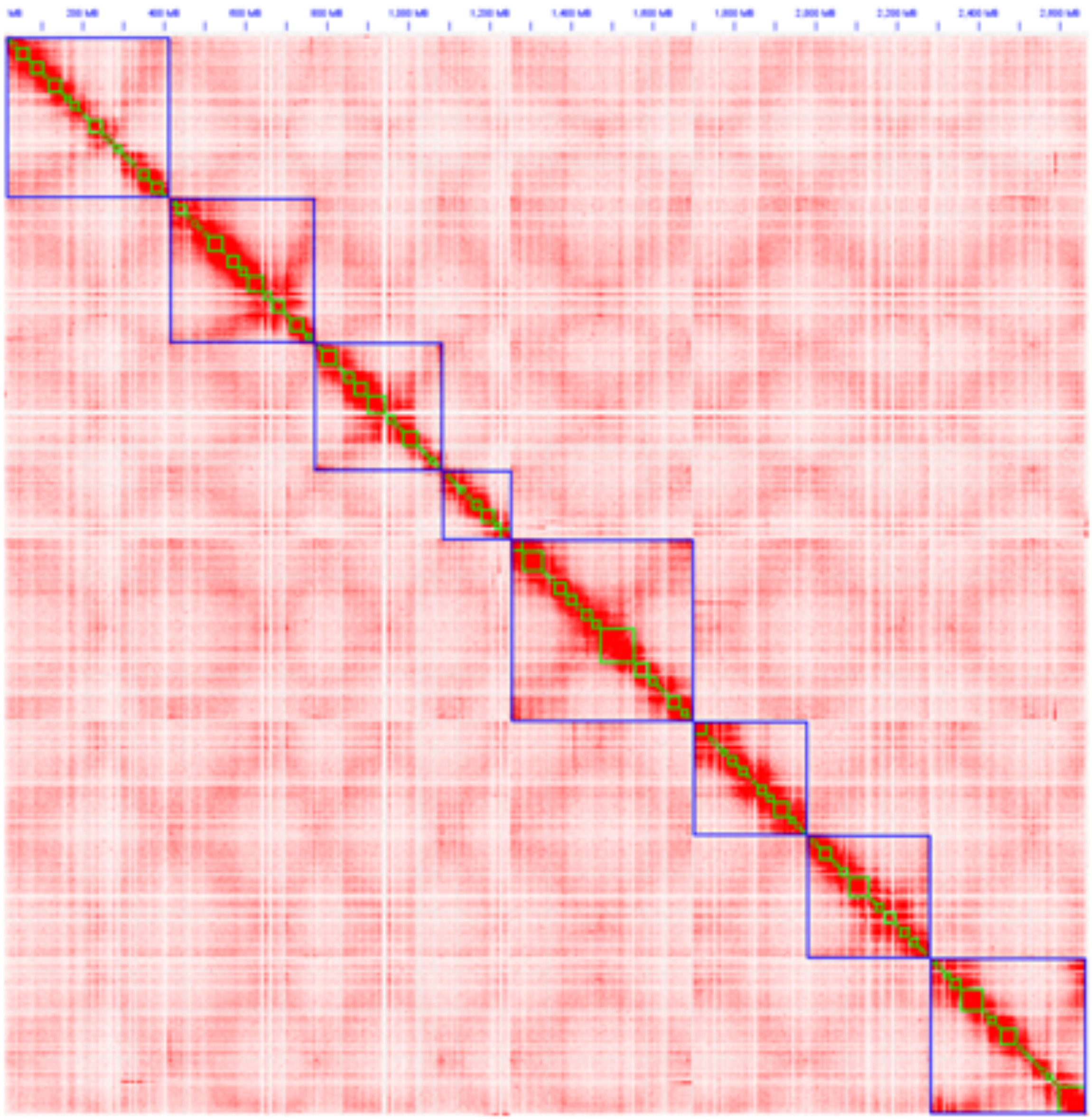

**Figure S6.** Hi-C contact map. Shown is a heatmap representation of chromatin interactions, derived from the Hi-C data. Please see the Material & Methods for more details. The diagonal represents the regions that are close to each other. The color scale indicates the frequency of chromatin interactions (red areas likely indicate high contact frequencies, meaning those regions of the genome are physically close in the 3D space of the nucleus. Blue rectangles along the diagonal highlight topologically associating domains with high intrachromatin contact, i.e., the 8 pseudochromsomes of *R. cassubicifolius*. The figure is attached to the Supplements.

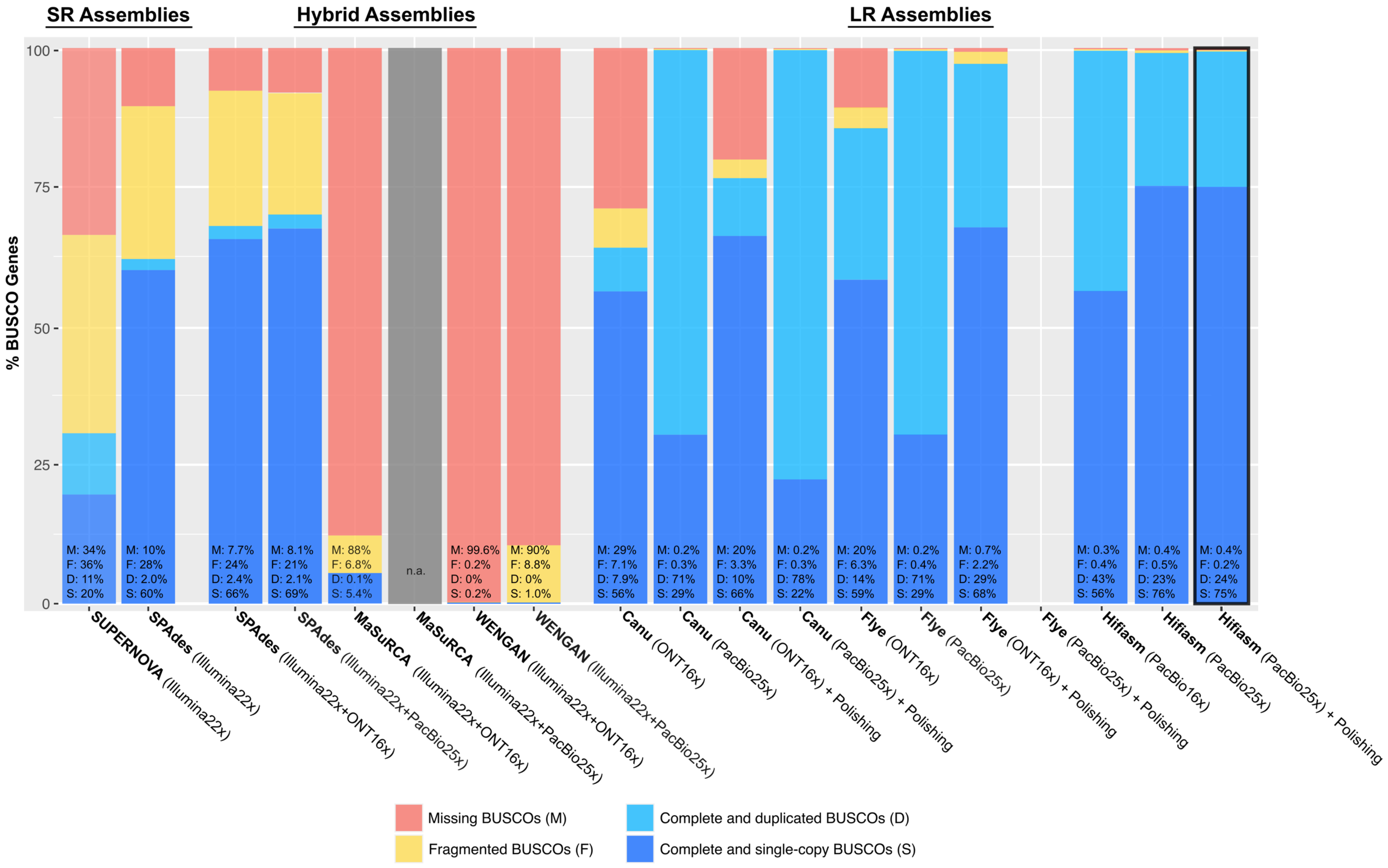
SR Assemblies
Hybrid Assemblies
LR Assemblies
% BUSCO Genes
100
75
50
25
0
M: 34%
F: 36%
D: 11%
S: 20%
M: 10%
F: 28%
D: 2.0%
S: 60%
M: 7.7%
F: 24%
D: 2.4%
S: 66%
M: 8.1%
F: 21%
D: 2.1%
S: 69%
M: 88%
F: 6.8%
D: 0.1%
S: 5.4%
n.a.
M: 99.6%
F: 0.2%
D: 0%
S: 0.2%
M: 90%
F: 8.8%
D: 0%
S: 1.0%
M: 29%
F: 7.1%
D: 7.9%
S: 56%
M: 0.2%
F: 0.3%
D: 71%
S: 29%
M: 20%
F: 3.3%
D: 10%
S: 66%
M: 0.2%
F: 0.3%
D: 78%
S: 22%
M: 20%
F: 6.3%
D: 14%
S: 59%
M: 0.2%
F: 0.4%
D: 71%
S: 29%
M: 0.7%
F: 2.2%
D: 29%
S: 68%
M: 0.3%
F: 0.4%
D: 43%
S: 56%
M: 0.4%
F: 0.5%
D: 23%
S: 76%
M: 0.4%
F: 0.2%
D: 24%
S: 75%
SUPERNOVA (Illumina22x)
SPAdes (Illumina22x)
SPAdes (Illumina22x+ONT16x)
SPAdes (Illumina22x+PacBio25x)
MaSuRCA (Illumina22x+ONT16x)
MaSuRCA (Illumina22x+PacBio25x)
WENGAN (Illumina22x+ONT16x)
WENGAN (Illumina22x+PacBio25x)
Canu (ONT16x)
Canu (PacBio25x)
Canu (ONT16x) + Polishing
Canu (PacBio25x) + Polishing
Flye (ONT16x)
Flye (PacBio25x)
Flye (ONT16x) + Polishing
Flye (PacBio25x) + Polishing
Hifiasm (PacBio16x)
Hifiasm (PacBio25x)
Hifiasm (PacBio25x) + Polishing
Missing BUSCOs (M)
Fragmented BUSCOs (F)
Complete and duplicated BUSCOs (D)
Complete and single-copy BUSCOs (S)

**Figure S7.** BUSCO assessments for different genome assembly strategies of the diploid sexual species *Ranunculus cassubicifolius* (LH040). SUPERNOVA, and partially SPAdes, MaSuRCA, and WENGAN (hybrid assemblies) used Illumina short-reads ('Illumina'). Canu, Flye, and HIFIASM, and partially SPAdes, MaSuRCA, and WENGAN (hybrid assemblies) used ONT/PacBio long-reads ('ONT', 'PacBio'). The polishing of the Canu, Flye, and HIFIASM assemblies includes the following steps: polishing via filtered ONT reads using Racon and Medaka / via filtered PacBio reads using Racon and polishCLR, respectively, and via filtered Illumina reads using POLCA (MaSuRCA pipeline). The best assembly in terms of complete BUSCO genes (S+D) is highlighted with a black square, that is HIFIASM (PacBio) + Polishing. Analyses are based on 1,614 BUSCO genes. See the Materials and Methods section, and Table S5 for more details. See Figure 5 for comparisons with downsampled (16x) PacBio reads. The figure file is attached to the Supplements.
